# The Toll and Imd pathway, the complement system and lectins during immune response of the nemertean *Lineus ruber*

**DOI:** 10.1101/2022.04.26.489627

**Authors:** Andrea Orús-Alcalde, Aina Børve, Andreas Hejnol

**Affiliations:** Sars International Centre for Marine Molecular Biology, University of Bergen, Bergen, Norway; University of Bergen, Department of Biological Sciences, Bergen, Norway

**Keywords:** Toll pathway, Imd pathway, Complement system, lectins, FreD-C, C-lectins, PRR

## Abstract

Innate immunity is the first line of defense against pathogens. In animals, the Toll pathway, the Imd pathway, the complement system, and lectins are well-known mechanisms involved in innate immunity. Although these pathways and systems are well understood in vertebrates and arthropods, they are understudied in other invertebrates. In order to shed light on immunity in the nemertean *Lineus ruber*, we performed a transcriptomic survey and identified the main components of the Toll pathway (e.g. *myD88, dorsal/dif/NFκB-p65*), the Imd pathway (e.g. *imd, relish/NFκB-p105/100*), the complement system (e.g. C3, *cfb*) and some lectins (FreD-Cs and C-lectins). *In situ* hybridization showed that *TLRβ1, TLRβ2* and *imd* and are expressed in the nervous system, the complement gene *C3-1* is expressed in the gut and the lectins in the nervous system, the blood, and the gut. To reveal their potential role in defense mechanisms, we performed immune challenge experiments, in which *Lineus ruber* specimens were exposed to the gram-negative bacteria *Vibrio diazotrophicus*. Our results show the upregulation of specific components of the Toll pathway (*TLRα3, TLRβ1*, and *TLRβ2*), the complement system (*C3-1*), and lectins (*c-lectin2* and *fred-c5*). Therefore, similarly to what occurs in other invertebrates, our study shows that components of the Toll pathway, the complement system and lectins are involved in the immune response in the nemertean *Lineus ruber*. The presence of these pathways and systems in *Lineus ruber*, but also in other spiralians, in protostomes and in deuterostomes suggest that these pathways and systems were involved in the immune response in the stem species of Bilateria.

## Background

Innate immunity is the first line of defense of plants and animals against pathogens (1,2). During innate immunity, Pathogen Recognition Receptors (PRR) are able to distinguish non-self from self by recognizing Pathogen Associated Molecular Patterns (PAMP). In animals, PRRs are present in the main pathways and systems involved in innate immunity, such as the Toll- and Imd pathways, or the complement system (3–5).

The Toll pathway is an immune pathway present in many species across the metazoan tree (4,6–8). In vertebrates, the receptors of this pathway, named Toll receptors (TLRs), are able to recognize and bind PAMPs directly. However, in insects, PAMP recognition is mediated by the Spätzle protein (Figure 1A). Once TLRs are activated, a signaling cascade, in which MyD88, Tube/Irak4, and Pelle/Irak1 are involved, triggers the entrance of NFκB transcription factors (Dorsal and Dif in *Drosophila* and NFκB-p65 in vertebrates) to the nucleus, which activate the expression of antimicrobial peptides and cytokines (7). Besides vertebrates and *Drosophila*, components of the Toll pathway have been identified in multiple invertebrate species (Figure 1D) (9–12). Moreover, immune challenge assays have shown that the Toll pathway is involved in immunity in mollusks and crustaceans (13,14).

**Figure 1.**
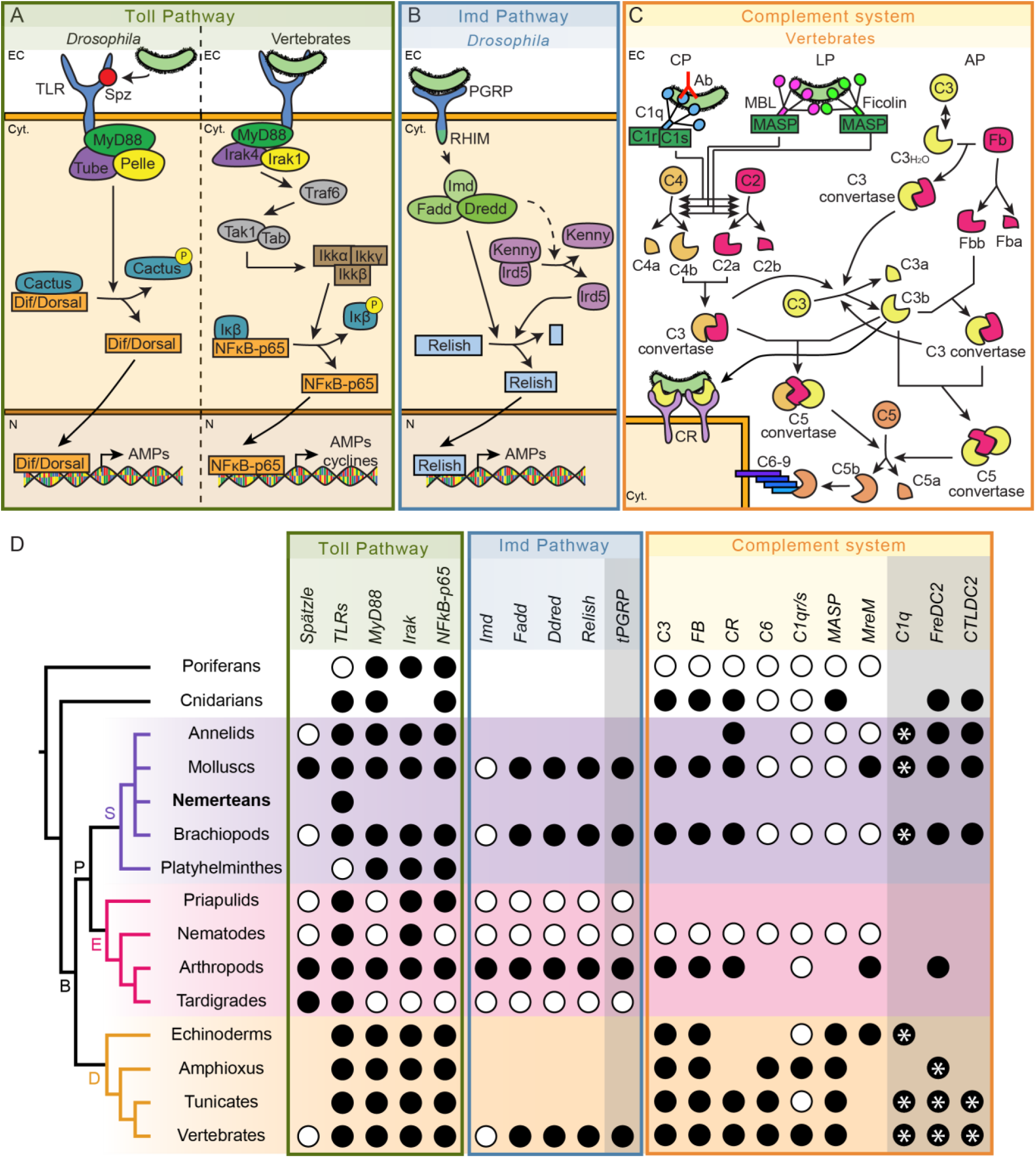
Toll pathway, Imd pathway, and complement system in metazoans. **A.** Toll pathway, **B.** Imd pathway, and **C.** complement system in *Drosophila* and vertebrates. Dashed arrows indicate indirect processes. **D.** Presence and absence of proteins belonging to the Toll and the Imd pathways, and to the complement system across metazoans. Orthologs of components of the Imd pathway have been found in vertebrates, however, these components belong to the vertebrate TNFα pathway, which is analogous to the arthropod Imd pathway. Greyish compartments within each pathway compartment indicate proteins that are uncertain to be involved in the pathway. Black circles indicate that the protein is present for that taxa, while white circles indicate its absence. For C1q, FreD-C, and C-lectin proteins, black circles with an asterisk (*) indicate the presence of proteins with collagen domains (C1qL, ficolin and MBL/GBL, respectively), while only black circles indicate presence of C1q, FreD-C and C-lectin proteins containing coiled coil regions instead of collagen domains (FreDC2 and CTLDC2). Nemertean phylogenetic position is highlighted in bold. Abbreviations: AP: alternative pathway; B: Bilateria; CP: classical pathway; Cyt: cytoplasm; D: Deuterostomia; E: Ecdysozoa; EC: extracellular space; Fb: Factor B; LP: Lectin pathway; N: nucleus; P: Protostomia; S: Spiralia; tPGRPs: long transmembrane PGRPs. For references for panel D, view Supplementary Table 1. Phylogeny according to (39).

The Imd pathway is involved in the arthropod immune response against bacteria (15–17). Bacterial recognition triggers the activation of some long peptidoglycan recognition protein receptors (PGRP-Ls) (18). Activation of these PGRP-Ls triggers a signaling cascade that includes the recruitment of the Imd, Fadd and Dredd proteins and culminates with the entrance of the transcription factor Relish into the nucleus (Figure 1B) (7). The existence of this pathway outside arthropods is not clear. Components of this pathway are not present in other ecdysozoans, such as priapulids, nematodes, and tardigrades (Figure 1D) (11). Moreover, even though no homologous pathway to the Imd pathway has been identified in vertebrates, the Imd pathway shows similarities with the vertebrate TNF-α pathway, as orthologous proteins (e.g. Fadd, Dredd/Caspase8, Relish/NFκB) are present in both pathways (19). However, the Imd protein is absent in vertebrates, and vertebrate PGRPs are not involved in TNF-α pathway activation. In spiralians, PGRPs and downstream components of this pathway are present in mollusks and brachiopods (10,13,20,21). However, PGRPs with RHIM motifs, essential for signal transduction in arthropod PGRPs, and the Imd adaptor have not been found in any of the two taxa.

The complement system is a proteolytic cascade involved in opsonization, phagocytosis, inflammatory regulation, and cytolytic processes. In vertebrates, this system is activated by three pathways: the classical, the lectin, and the alternative pathways (Figure 1C) (22–24). C1q is the receptor of the classical pathway, whereas the lectin pathway is activated by mannose-binding lectins (MBL) and ficolins (25–28). These receptors trigger the activation of serine proteases (e.g. C1r, C1s, MASPs), which lead to the cleavage of the C3 protein. The alternative pathway is activated by the spontaneous hydrolysis of the C3 (23,25–28). C3 is the central component of the complement system, being the point where the three activating pathways converge (23). The cleaved C3 protein can be detected by complement receptors (CR) present in phagocytic cells (29), but it can can trigger the formation of the membrane attack complex (MAC) to induce cell lysis (30). Although the complement system has been well studied in vertebrates, little is known about how this system functions in invertebrates. The core components of the complement system (C3, Factor B, and complement receptors) are widespread through the metazoan tree (Figure 1D) (21,31–34). Nevertheless, although C1q proteins have been detected in spiralians (35), ficolins, MBL and downstream proteins (e.g. C6) seem to be present in deuterostome invertebrates but not in protostomes (35–38). However, C-lectins and Fibrinogen-related domain-containing proteins (FreD-C) with similar domain architectures than ficolins and MBL have been identified in spiralians, suggesting that these proteins could perform analogous functions to the vertebrate MBLs and ficolins (10,21,35). Furthermore, although the serine proteases C1r, C1s and MASPs have not been found in protostomes, MASP-related Molecules (MreM) are present in invertebrates (35).

Besides in complement activation, FreD-C and C-lectins proteins are also involved in a large variety of immune processes, independent from the complement system. FreD-Cs are a family of proteins characterized by the presence of a fibrinogen domain (FBG) (40). In vertebrates, besides ficolins, there is a wide variety of FreD-C proteins (e.g. tenascins, angiopoietins), which also have immune functions. In invertebrates, FreD-Cs have been observed to play a role in bacteria agglutination (41). Moreover, FreD-Cs are also involved in neuronal development and allorecognition (40). C-lectins are characterized for having at least a C-lectin domain, although other domains can also be present (42). These proteins are very abundant in invertebrates and they are involved in a broad variety of immune functions (e.g. agglutination, opsonization, phagocytosis, encapsulation) (43,44).

The Toll pathway, the Imd pathway, the complement system and lectins have been well studied in *Drosophila* and vertebrates. However, although some studies have been conducted in mollusks and brachiopods (10,21,35), studies on these pathways and system in other spiralian groups are scarce (Figure 1D). In this study, we aim to relate the common innate immunity pathways to the immune response in the nemertean *Lineus ruber* (45). Therefore, we performed a survey in the *Lineus ruber* transcriptome in order to detect components belonging to the Toll and the Imd pathways, the complement system, and lectins, and we confirmed the presence of these pathways and systems. Moreover, we studied the expression of members of these pathways and lectins by *in situ* hybridization, showing that they are expressed in various organs and tissues. Finally, we performed immune challenge experiments in *Lineus ruber* in order to study the changes in expression of some of these genes in response to bacterial infection, revealing that all the genes studied, except for *imd*, seem to be involved in immunity against gram-negative bacteria.

## Results

### Presence of orthologs of the Toll-, the Imd pathways, the complement system and lectins components in *Lineus ruber*

We performed transcriptomic surveys in order to identify *Lineus ruber* components of the Toll-and the Imd pathways, the complement system and lectins. All protein sequences retrieved from the surveys are available in Supplementary Figure 1.

#### The Toll pathway is present in *Lineus ruber*

Our previous study revealed the presence of 6 TLRs in the transcriptome of *Lineus ruber* (12). Here, we detect *myD88,* an *irak* gene, and the transcription factor *dorsal/diff/NFkB-p65* in the *Lineus ruber* transcriptome. Domain architecture (Figure 2), BLAST (Supplementary Table 4) and phylogenetic analyses (Supplementary Figure 2) confirm the identity of these proteins. The presence of an ortholog of *spätzle*, however, was not detected.

**Figure 2.**
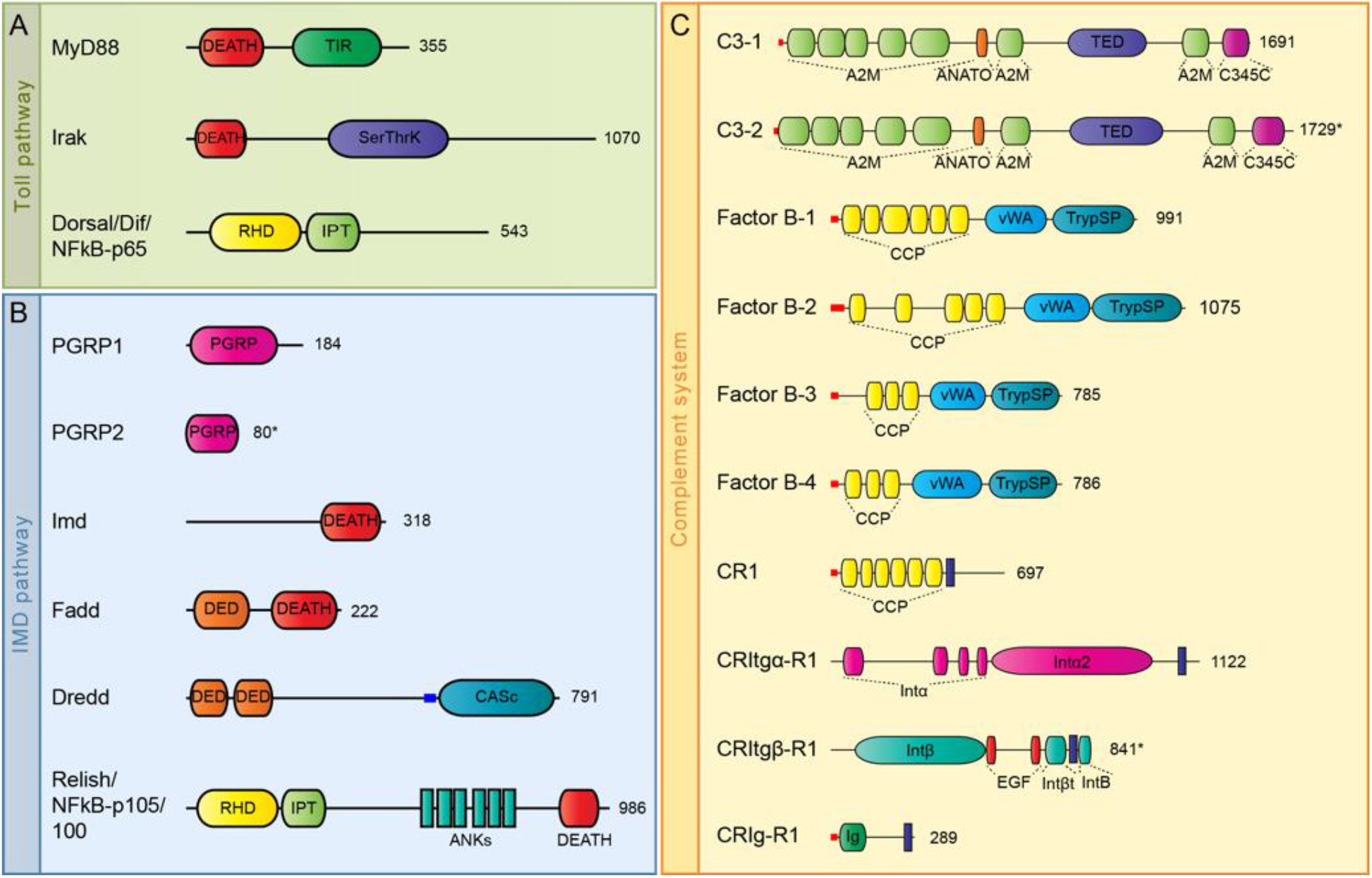
Domain architecture analyses of putative proteins belonging to the Toll-, the Imd pathways and the complement system in *Lineus ruber*. **A.** Proteins belonging to the Toll pathway. **B.** the Imd pathway and **C.** the complement system.An example of each complement receptor type (CR, CRItgα, CRItgβ and CRIg) is shown. For other complement receptor proteins identified in this study, see Supplementary Figure 3. Numbers indicate the length of the protein in amino acids. Asterisks after the amino acid number indicate partial proteins. A2M: α2-macroglobulin family domain; ANATO: Anaphylatoxin homologous domain; ANK: Ankyrin domain; CASc: caspase domain; CCP: complement control protein; DED: Death effector domain; EGF: Epidermal growth factor domain; Ig: Immunoglobulin domain; Intα: Integrin-α domain; Intβ: Integrin-β domain; Intβt: Integrin-β tail domain; IPT: Ig-like, plexin, transcription factors domain; PGRP: Peptidoglycan recognition protein domain; RHD: Rel Homology Domain; SerThrK: Serine/Threonine protein kinase domain; TED: Thioester domain; TIR: Toll-Interleukin 1 receptor domain; TrypSP: Trypsin-like serine protease domain; vWA: von Willebrand factor type A domain. Small blue and red bars represent coiled coils and signal peptides, respectively. Blue rectangles represent transmembrane domains.

*Lineus ruber* proteins belonging to the Toll pathway contain the characteristic domains of these proteins (Figure 2A), which are also found in ortholog proteins of other species. The *Lineus ruber* MyD88 ortholog contains the TIR and DEATH domains characteristic of MyD88 proteins (46); whereas a DEATH and a Serine/Threonine protein kinase domain, typical features of the Pelle/Irak1 and Irak4 proteins (47,48), were identified in the *Lineus ruber* Irak ortholog. Furthermore, two proteins belonging to the NFκB family were also identified, with our phylogenetic analysis of NFκB proteins showing that one of them is an ortholog to the *Drosophila* Dorsal and Dif proteins and the vertebrate NFκB-p65 (Supplementary Figure 2). This protein contains two Relish domains (an RHD and an IPT domains), which are also found in its orthologs (49). The second protein was identified as the Relish/NFκB-p105/100 ortholog, the transcription factor of the Imd pathway (see below).

#### Key components of the IMD-like pathway are present in *Lineus ruber*

Since orthologs of the *Drosophila* Imd adaptor have not yet been identified in mollusks and brachiopods, the existence of this pathway in spiralians has been questioned (13,21). Therefore, we performed a survey in the *Lineus ruber* transcriptome to identify potential components of this pathway. Our results show the presence of two PGRPs and one *imd*, *fadd, dredd* and *relish/NFκB-p105/100* genes.

Our survey identified two PGRPs genes, named *PGRP-1* and *PGRP-2*, respectively. Our analyses show that *PGRP-1* encodes a short protein that contains one PGRP domain, but lacks the transmembrane domains, signal peptide or RHIM motifs (Figure 2B), elements that are present in *Drosophila* PGRP receptors involved in the Imd pathway (18). Furthermore, PGRP-1 BLAST best hit is the short PGRP-S2 of the mollusk *Hyriopsis cumingii* (Supplementary Table 4). Thus, PGRP-1 is a short PGRP, which is probably not involved in Imd pathway activation. Since from our survey we could only obtain a partial sequence for *PGRP-2*, it was not possible to determine if a transmembrane domain and a RHIM motif are present in the PGRP-2 protein. Furthermore, a survey for RHIM motifs in the *Lineus ruber* transcriptome did not detect any sequence encoding this motif. However, the lack of long PGRPs in the transcriptome only shows that these genes are not expressed in that specific stage. Therefore, as no genome of *Lineus ruber* is available, we performed surveys for PGRP domains and RHIM motifs in the genome of the nemertean *Notospermus geniculatus* (50). Our results show the presence of 8 genes encoding for PGRP domain-containing proteins (Supplementary Figure 4A), but genes encoding for proteins containing RHIM were not detected. All *Notospermus geniculatus* PGRP proteins are shorter than 350 amino acids, with the exception of PGRP5, which is 517 amino acids long. According to Dziarski and Gupta, 2006 (51), short PGRPs have an approximate length of 200 amino acids, while long PGRPs are at least the double in length. Domain architecture analyses show that *Notospermus geniculatus* PGRP5 has a signal peptide and a C-terminal PGRP domain, but no transmembrane domains. Therefore, we suggest that this protein is an extracellular protein which is not involved in Imd pathway activation. Furthermore, we performed a phylogenetic analysis of PGRP proteins, including the *Lineus ruber, Notospermus geniculatus*, and *Drosophila melanogaster* PGRPs (Supplementary Figure 4B). This analysis shows that all nemertean PGRPs cluster together forming a sister clade to the *Drosophila melanogaster* short PGRPs and PGRP-LB, proteins that are not involved in Imd pathway activation. Therefore, our results suggest that nemertean PGRP are probably not involved in Imd pathway activation.

Furthermore, we identified an *imd* gene in the *Lineus ruber* transcriptome. The *Lineus ruber* Imd protein contains the characteristic DEATH domain of Imd proteins (Figure 2B). Moreover, our survey also retrieved the presence of a *fadd* and a *dredd* genes. Our analyses show that Fadd and Dredd proteins contain Death effector domains and, in the case of Dredd, one Caspase domain was additionally found, as in the *Drosophila* ortholog (52) (Figure 2B). Finally, as mentioned above, two NFκB genes are present in the *Lineus ruber* transcriptome. Our analyses identified one of these NFκB proteins as the ortholog of Relish/NFκB-p105/100 (Figure 2B; Supplementary Figure 2). This protein contains two Relish homology domains (an RHD and an IPT domains), six ANK repeats – domains that are also present in the *Drosophila* and vertebrate orthologs (49,53) – and a DEATH domain, which has also been observed in the Relish protein of other arthropods (54,55).

#### The complement system is present in *Lineus ruber*

Our *Lineus ruber* transcriptomic survey revealed 2 *C3* genes, 4 *Complement factor B* (*Cfb*) genes, and up to 26 putative genes encoding for complement receptors (CR) (Figure 2C, Supplementary Figure 3). Genes encoding for ortholog proteins to the vertebrate membrane attack complex proteins (C6-9) and for the serine proteases C1s/C1r/MASP/MReM were not identified.

Our analyses show that the presence of 2 *C3* genes in the transcriptome of *Lineus ruber* (named here *C3-1* and *C3-2*) (Figure 2C, Supplementary Figure 2, Supplementary Table 4). Domain architecture analyses show that these proteins contain α2-macroglobulin domains, an anaphylatoxin domain, a thioester region, and a C345C C-terminal domain (Figure 2C). Furthermore, we also unraveled the presence of four *cfb* genes encoding for 4 Factor B proteins (Factor B-1 to Factor B-4) (Figure 2C, Supplementary Figure 2, Supplementary Table 4). Domain architecture analyses show that these four proteins are composed by complement control protein domains (CCP), a von Willebrand factor (vWF) type A domain, and a Trypsin-like serine protease domain (TrypSP) (Figure 2C). However, *Complement factor C* (*Cfc*) genes, which are homologous to *Cfb* and have been detected in some protostomes (e.g. arthropods, brachiopods) (21), were not found in the *Lineus ruber* transcriptome. Additionally, we identified up to 26 putative genes encoding for complement receptors (CR) with similar domain composition than the vertebrate complement receptors (Figure 2C, Supplementary Figure 3). Similarly to vertebrate complement receptors CR1 and CR2 (56), 6 proteins were found containing multiple CCP repeats and a transmembrane domain (CR1 to CR6) (Figure 2C, Supplementary Figure 3). Furthermore, in vertebrates, integrin-α (CD11b) and β (CD18) proteins assemble to form the complement receptors CR3 and CR4 (57). Here, we also identified 4 transmembrane proteins containing integrin-α or β domains (CR-Itgα1 and 2; and CR-Itgα1 and 2β) (Figure 2C, Supplementary Figure 3). However, whether these proteins heterodimerize to constitute complement receptors in *Lineus ruber* is not assessed in this study. Moreover, the vertebrate CRIg are constituted by one or more immunoglobulin domains and a transmembrane domain (58). Here, we show the presence of 16 genes encoding for proteins with similar domain composition (Figure 2C, Supplementary Figure 3).

#### Putative activators of the complement system

In order to investigate the possible pathways by which the complement system could be activated in *Lineus ruber*, we performed a transcriptome survey to identify genes encoding for proteins containing FBG, C-lectin, or C1q domains, since these domains are present in vertebrate proteins involved in complement system activation (27,28).

Our survey reveals 14 genes encoding for proteins containing a FBG domain (FreD-C1 to FreD-C14). While all these proteins have a FBG domain, only FreD-C1 contains a CCP domain (Supplementary Figure 5). Moreover, although the remaining proteins do not have any other domains, FreD-C3, FreD-C4, FreD-C7, and FreD-C11 contain coiled coil motifs and, therefore, belong to the FreDC2 subfamily (35). Proteins belonging to this subfamily have been suggested to form multimeric proteins, similarly to the vertebrate ficolins, that could activate the complement system (35). Our results also show the presence of 39 C-lectin genes in the transcriptome of *Lineus ruber* (*c-lectin1* to *c-lectin39*). These genes encode for proteins with a large variability of domain composition (e.g. leucine rich repeat domains, von Willebrand factor type-A domain, Complement control protein modules) (Supplementary Figure 5), being some of these domains also found in vertebrate C-lectins (42). Additionally, although some C-lectin proteins constituted only by a sole C-lectin domain were also identified, no proteins containing a collagen domain or a coiled coil region together with a C-lectin domain were found. This suggests that C-lectin proteins would not be involved in complement activation in *Lineus ruber*. Furthermore, we identified three C1q genes (*C1q-1* to *C1q-3*) in the *Lineus ruber* transcriptome. These genes encode for proteins formed by a collagen and a C1q domains and, therefore, they are C1qL proteins that could activate the complement system (Supplementary Figure 5).

Together, the findings from our transcriptome surveys in *Lineus ruber* confirm the existence of the Toll- and an Imd pathways, the complement system and lectins in this organism. The Toll pathway proteins MyD88, Irak and Dorsal/Dif/NFκB-p65 were identified; as well as the Imd pathway components Imd, Fadd, Dredd and Relish/NFκB-p105/100. However, no putative receptors for this pathway were identified. Our results also show the presence of the necessary components to constitute a functional complement system, since C3, Factor B, and complement receptors were identified. Additionally, the presence of coiled coil motifs in FreD-C proteins and collagen domains in C1q proteins suggest a possible involvement of these proteins in complement system activation.

### Genes with putative immune functions are expressed in a variety of tissues in *Lineus ruber*

In order to study the expression of the aforementioned genes, whole mount *in situ* hybridization (WMISH) was performed in 40 and 60 days *Lineus ruber* juveniles (Figure 4). As all nemerteans, *Lineus ruber* have an eversible proboscis used to catch prey (59). *Lineus ruber* nervous system is formed by a brain and two lateral and a dorsal nerve cords, as well as cephalic nerves that emerge from the brain and extend to the anterior area of the head, innervating the frontal sensory organ and eyes (59–62). Furthermore, this species possesses a closed circulatory system that consists of two lateral and a dorsal blood vessels that run parallel to the lateral and the dorsal nerve cords (59). By the anterior part, these vessels are connected near to the brain and form a cephalic vascular loop surrounding the proboscis. The mouth is located in the anterior area of the trunk, opening to the gut, which is extended to the posterior part of the animal. These organs and systems are already present in *Lineus ruber* juveniles (Figure 4A).

The results of our whole mount *in situ* hybridization reveals that both *TLRβ1* and *TLRβ2* are expressed in the lateral nerve cords as well as in the brain and cephalic organs in 40-day juveniles (Figure 3A, B and B’). Similarly, *imd* is also expressed in the brain and the lateral nerve cords (Figure 3D). Furthermore, the complement *C3-1* gene has found to be expressed in the gut and the blood both in 40 and 60 days juveniles (Figure 3E and F). Although at 60 days of development, the expression of *C3-1* is strong in the cephalic vascular loop, at 40 days it is very faint in this region. In 60-day juveniles, *fred-c1* is expressed in the brain, the ventrolateral nerve cords and the cephalic nerves (Figure 3F); whereas *fred-c5* is expressed in the blood (Figure 3G). Expression of *c-lectin2* was detected in the brain and the cephalic nerves at both stages analyzed (Figure 3H and I). *c-lectin3* is expressed in the proboscis area of the head (Figure 3E). *c-lectin5, c-lectin9*, and *c-lectin10* are expressed in the nervous system, with *c-lectin5* expressed in areas of the brain and in the cephalic nerves (Figure 3F); *c-lectin9* in a pair of brain lobes and in the frontal organ (Figure 3G); and *c-lectin10* in the brain, the two ventrolateral nerve cords and in the eyes (Figure 3H). *c-lectin11* expression was found to be expressed in the gut (Figure 3I) and *C1q-1* in the brain, the ventrolateral nerve cords, the cephalic nerves, and in the frontal sensory organ (Figure 3J).

**Figure 3.**
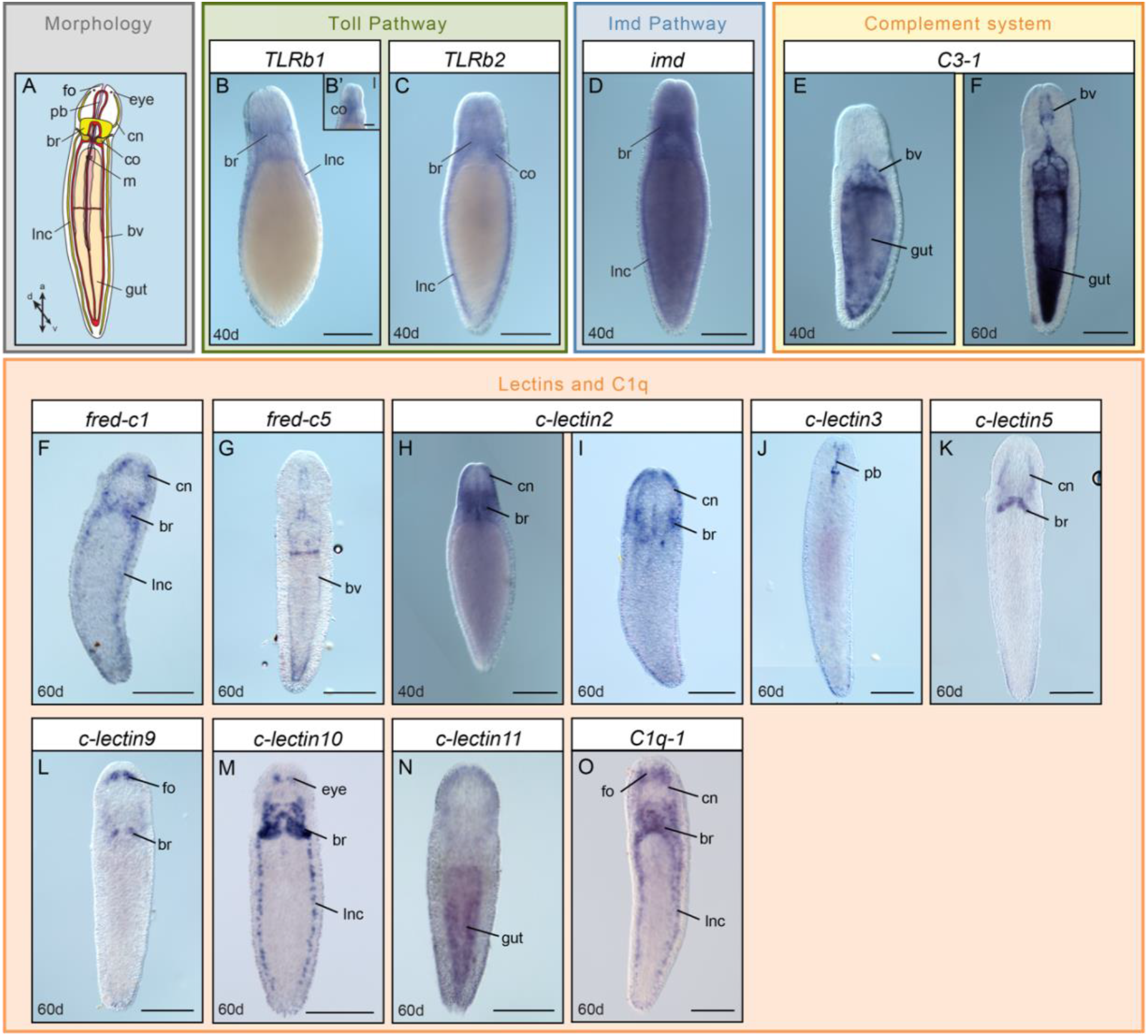
Expression of immune genes in *Lineus ruber*. **A.** Schematic representation of *Lineus ruber* anatomy. **B-O.** Whole mount *in situ* hybridization (WMISH) of *(B-C)* genes belonging to the Toll pathway; *(D)* to the Imd pathway; *(E-F)* of the complement system and *(F-O)* lectins and the C1q family. The name of each gene is indicated above each panel. Besides panel B’, all panels show dorso-ventral (d,v) views, being anterior (a) orientation to the top. Panel B’ shows lateral (l) orientation. Scalebars indicate 250μm, with the exception of the scalebar in B’, which indicates 100μm. Abbreviations: br: brain; bv: blood vessel; cn: cephalic nerve; fo: frontal sensory organ; g: gut; l: lateral view; lnc: lateral nerve cord; m: mouth; pb: proboscis. Scheme drawn after (59).

**Figure 4.**
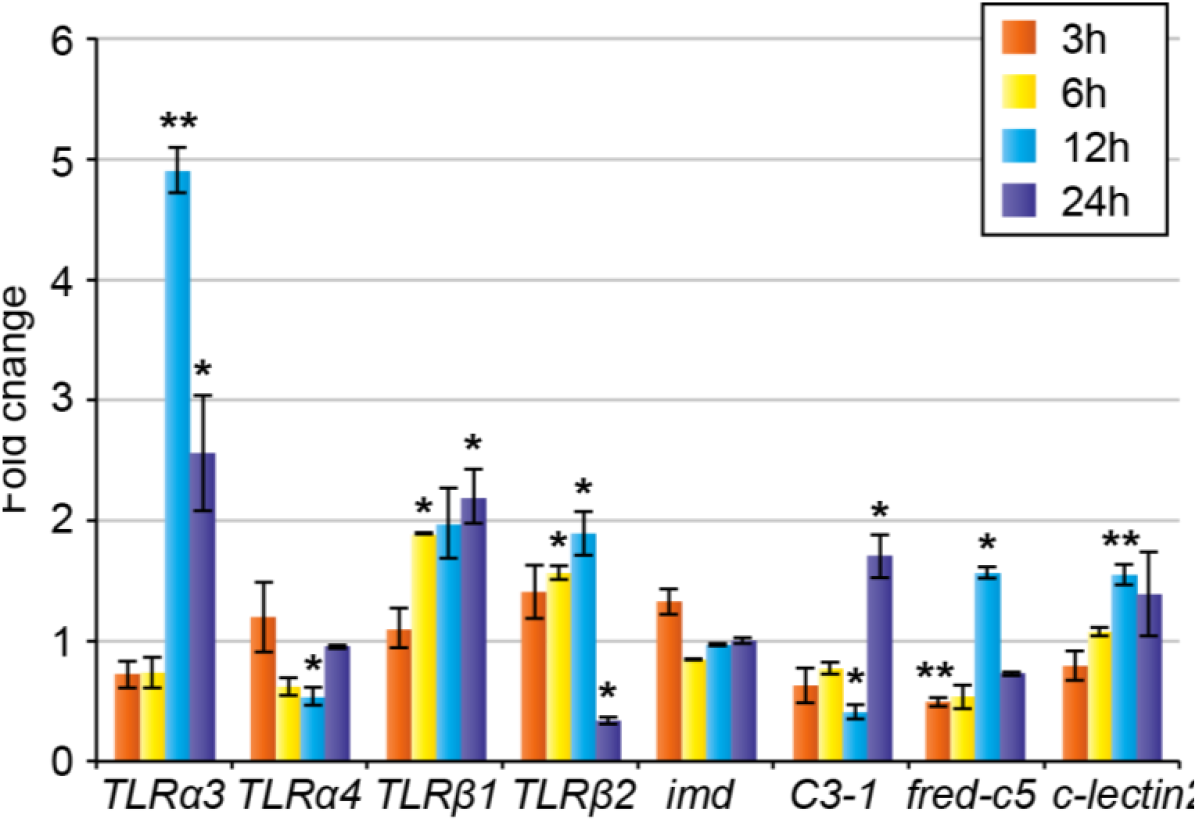
Relative expression of immune genes in response to infection to *Vibrio diazotrophicus*. Expression of immune genes in the different timepoints was normalized to the gene expression levels in the control animals. The fold change was calculated using the 2^-ΔΔCT^ method (63), standardizing to 1 the expression level for control animals. Asterisks indicate significant differences in gene expression between control and infected animals, being evaluated by performing statistical ANOVA tests (* indicates p-value <0.05; ** indicates p-value <0.01). Bars indicate the standard error between infected biological replicates.

Moreover, we also performed *in situ* hybridization for genes encoding other components of the Toll pathway (*TLRα1, TLRα2, TLRα3, TLRα4, myd88*, and *dorsal/dif/NFκB-p65*), the Imd-like pathway (*relish/NFκB-p105/100*), other FreD-Cs (*fred-c2, fred-c3, fred-c4, fred-c6*, and *fred-c7*), C-lectins (*c-lectin1, c-lectin4, c-lectin5, c-lectin6*, and *c-lectin7*) and the C1q family member *C1q-2*, but no expression was obtained. This could be explained by the absence or very low expression levels, non-detectable by WMISH, of these genes in healthy juvenile animals.

### The degree of expression of immune related genes is altered in infected *Lineus ruber*

In order to understand whether the candidate genes were involved in immunity against gram-negative bacteria, we exposed healthy adult *Lineus ruber* specimens to *Vibrio diazotrophicus* for 3h, 6h, 12h, and 24h. The expression levels of the *TLRα3, TLRα4, TLRβ1, TLRβ2, imd, fred-c5, C3-1*, and *c-lectin2* genes were evaluated at those timepoints performing quantitative real-time PCR (qPCR) and compared between control and infected animals.

Our results show that the expression of most genes was not significantly altered at 3h of infection, with the exception of *fred-c5* (Figure 4). *TLRα3* expression levels were upregulated in infected animals at 12h, remaining increased at 24h of infection. In contrast, *TLRα4* expression was downregulated at 12h of infection, and never upregulated at the timepoints of study. *TLRβ1* expression levels were increased at 6h and remained such at 12h and 24h of infection. Interestingly, *TLRβ2* was upregulated at 6h and 12h of infection, but its expression decreased by 24h of infection. *imd* expression did not vary at any of the studied timepoints. The complement factor *C3-1* was downregulated at 12h of infection but its expression levels were increased by 24h of infection. *fred-c5* expression was downregulated already at 3h of infection; however, its expression increased at 12h and reached similar expression levels to control animals at 24h of infection. *c-lectin2* expression was upregulated at 12h in infected animals, while its expression dropped to similar expression levels to control animals at 24h of infection.

Thus, our results show that the gram-negative bacteria *Vibrio diazotrophicus* triggers an immune response in the nemertean *Lineus ruber,* in which the Toll receptors, the complement system and the lectins *fred-c5* and *c-lectin2* are involved. However, although the Imd pathway is involved in defense against gram-negative bacteria in arthropods (15–17), the *imd* gene expression remained unaltered in all the studied timepoints.

## Discussion

### The Toll pathway is involved in the gram-negative immune response in *Lineus ruber*

The Toll pathway is a pathway involved in immunity that is present across many metazoan lineages (4,6–8). In a previous study, we unraveled the presence of 6 TLRs in the nemertean *Lineus ruber* (12). In this study, we identified the presence of the MyD88 adaptor, an Irak protein and a Dorsal/Diff/NF-κB-p65 protein in this nemertean species, but no Spätzle protein ortholog was identified (Figure 2A). Therefore, we suggest that TLRs in *Lineus ruber* are probably activated directly by the pathogen, similar to other spiralians and deuterostomes (7). Considering our results under the scope of the existing knowledge on the Toll pathway in other animals, we suggest that, once the *Lineus ruber* TLRs are activated, a signaling cascade in which MyD88 and an Irak protein are involved, trigger the entrance of the *Lineus ruber* Dorsal/Diff/NF-κB-p65 into the nucleus (Figure 5). Our results show that *TLRβ1* and *TLRβ2* are expressed in the nervous system in juvenile *Lineus ruber*, suggesting that these receptors could be involved in immunity and/or development of this tissue (Figures 3 and 5). Additionally, upon exposure to the gram-negative bacteria *Vibrio diazotrophicus, TLRα3, TLRβ1* and *TLRβ2* are upregulated; whereas *TLRα4* expression did not vary or was downregulated (Figure 4). These findings show that at least three TLRs in *Lineus ruber* are involved in gram-negative bacterial response. Although the Toll pathway is not involved in *Drosophila* defense against gram-negative bacteria, upregulation of TLRs in *Lineus ruber* against gram-negative infection is in agreement with findings in other invertebrate species, such as other arthropods and mollusks (13,14). Furthermore, *TLRα4* could be either involved in other stages of infection against gram-negative bacteria or involved in the detection of other pathogens (e. g. gram-positive bacteria, fungi, virus), or not be involved in immunity.

**Figure 5.**
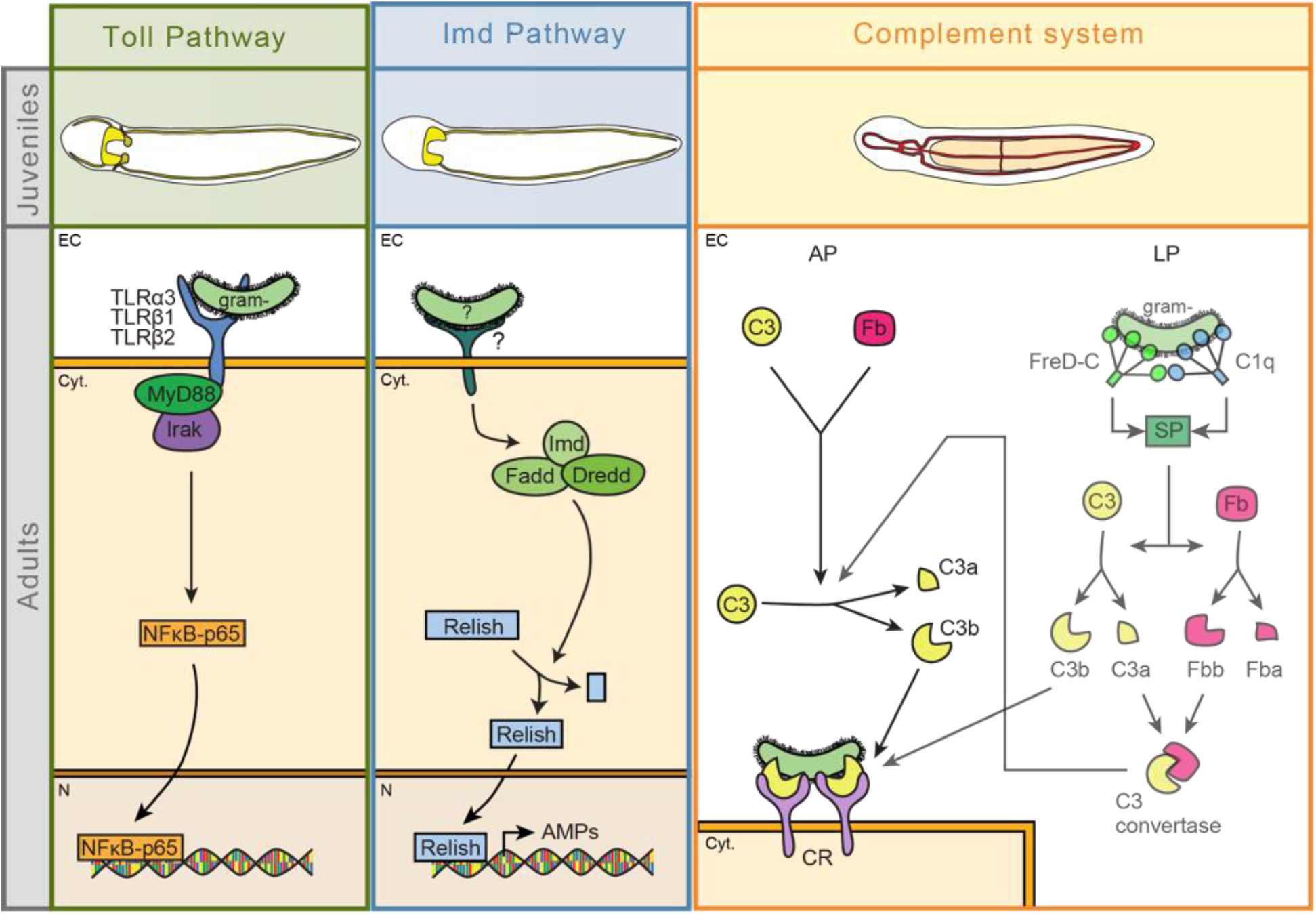
The Toll pathway, Imd pathway and complement system in *Lineus ruber*. “?” indicates uncertainty about the identity of the agent activating the pathway or the receptor involved in it. Semi-transparent schemes indicate the uncertainty in the existence of this pathway. AP: alternative pathway; CR: Complement receptor; Cyt.: cytoplasm; EC: Extracellular space; Fb: Factor B; gram-: Gram-negative bacteria; LP: Lectin pathway; N: Nucleus; SP: Serine protease.

### The Imd-like pathway is present in *Lineus ruber* but it seems to not be involved in immunity against gram-negative bacteria

The Imd pathway has been shown to be a pivotal pathway in the defense against gram-negative bacteria in arthropods (15–17). However, no orthologs of the Imd protein, the key component of this pathway, have been found in spiralians and vertebrates (6,10,13,19,21). In this study, we surveyed for components of this pathway in the transcriptome of the nemertean *Lineus ruber*, identifying for first time an *imd* ortholog in spiralians (Figure 2B). Besides *imd*, we also found the downstream components *fadd, dredd*, and *relish/NFκB-p105/100*. Except for the Imd protein, orthologs of proteins belonging to the arthropod Imd pathway have also been identified in mollusks, brachiopods, and vertebrates (6,13,19,21). Among these proteins, transmembrane PGRPs compatible with Imd pathway activation are present both in brachiopods and mollusks (10,21). However, although our analysis identified 2 PGRPs in the *Lineus ruber* transcriptome and 8 in the *Notospermus geniculatus* genome (Supplementary Figure 4), we did not find evidences for them to be the receptors of this pathway. Therefore, although our findings might indicate the existence of this pathway in nemerteans, this pathway would be activated by other receptors than PGRPs (Figure 5B). Furthermore, our results show that *imd* is expressed in the nervous system in juveniles (Figure 3 and 5). However, although this gene is upregulated after gram-negative bacterial exposure in arthropods (6,16,17), exposure of adult *Lineus ruber* to the gram-negative bacteria *Vibrio diazotrophicus* did not result in differences in expression of this gene (Figure 4). These results indicate that the Imd pathway is probably not involved in gram-negative response in *Lineus ruber* or it is involved in other timepoints of infection not tested here. Furthermore, its involvement in the immune response towards other types of pathogens cannot be excluded.

### The complement system is involved in *Lineus ruber* immunity against gram-negative bacteria

Previous studies show the presence of a complement system formed by C3 and Factor B proteins in spiralians, such as mollusks or brachiopods (Figure 1D) (10,21,32,33). Our study shows that two C3 genes and four Cfb genes are present in the transcriptome of the nemertean *Lineus ruber* (Figure 2C). Furthermore, up to 26 putative genes encoding for complement receptors proteins with similar domain architectures than the human complement receptors (56–58)) were also detected in our *Lineus ruber* transcriptomic survey (Figure 2C; Supplementary Figure 3). Since C3, Factor B, and complement receptors constitute the core components the alternative pathway, the presence of these proteins in *Lineus ruber* suggests the presence of this complement pathway in this species (Figure 5C).

Although the lectin pathway is thought to have been emerged in early chordate evolution (38), recent studies suggest the presence of this pathway in spiralians (21,35). Here, we show that multiple FreD-C and C-lectin proteins are present in *Lineus ruber* (Supplementary Figure 5). Although no collagen domains have been found within these proteins, domain that is always present in vertebrate FreD-C and C-lectins activating this pathway (64,65), four proteins with coiled coil regions and an FBG domain were detected (FreD-C3, FreD-C4, FreD-C7, and FreD-C11). Previous studies have shown that FreD-C and C-lectin proteins containing coiled coil motifs are multimeric proteins that are also present in other spiralians and suggested their involvement in complement activation (10,21,35,66). Therefore, it is plausible that *Lineus ruber* FreD-Cs could activate the complement system via the lectin pathway (Figure 5C).

Furthermore, the classical pathway to activate the complement system is also considered to be a vertebrate innovation, since antibodies of the adaptive immunity are often involved in this pathway (38,67). However, as activation of the complement system by C1q is not always antibody-dependent (22,68) and C1q proteins containing collagen domains are widespread in metazoan species (10,32,35), it has been suggested that C1q proteins could activate the complement system in invertebrates in an antibody-independent way (21,35). Here, we found 3 genes encoding for C1q proteins containing a collagen domain in the *Lineus ruber* transcriptome (Supplementary Figure 5), suggesting that these proteins could be putative activators of the complement system. However, although the complement system could likely be activated in

*Lineus ruber* either by FreD-C and/or C1q proteins, the mechanism to circumvent the lack of the serine proteases MASP, C1r, and C1s and cleave of C3 and Factor B to form the C3 convertase has yet to be elucidated. It has been suggested that MreM could perform this function in other spiralians (35). However, we did not find MreM genes in the transcriptome of *Lineus ruber*.

Furthermore, our results show that the complement gene *C3-1* is expressed in the blood and the gut in *Lineus ruber* juveniles (Figures 3 and 5). Additionally, we show that this gene is upregulated in adult *Lineus ruber* after exposure to *Vibrio diazotrophicus* (Figure 4), suggesting that *Lineus ruber* complement could be activated in response to gram-negative bacterial infection. This is in concordance with previous studies showing the up-regulation of complement components after exposure to gram-negative bacteria in other invertebrates, such as mollusks, cnidarians and invertebrate deuterostomes, (32,69–72). Additionally, upregulation of complement system components in invertebrate deuterostomes and mollusks has also been observed to occur after exposure to gram positive bacteria (71,72). However, in this study, activation of the complement system by other pathogens (e.g. gram-positive bacteria, fungi) was not assessed and, therefore, cannot be excluded.

### FreD-C and C-lectin proteins, likely not part of the complement system activation, could also be involved in immunity in *Lineus ruber*

FreD-C and C-type lectins are proteins with high structural and functional diversity (42,43,73). In our study, besides the presence of FreD-Cs putatively involved in *Lineus ruber* complement activation (see above), we show the presence of 10 FreD-Cs and 39 C-lectins with domain architectures not suitable for this function (Supplementary Figure 5). Therefore, these proteins must have other functions than complement activation. In here, we show that both *fred-c5* and *c-lectin2*, expressed in the blood and the head nervous system respectively (Figure 3C), are upregulated after gram-negative bacterial exposure (Figure 4) and, therefore, they are involved in immunity against this type of bacteria. Other FreD-Cs and C-lectins are expressed in various tissues (e.g. gut, anterior proboscis and nervous system) (Figure 3), suggesting that they could be involved in immunity in those tissues.

## Conclusions

In this study, we identified key components of the Toll and Imd pathways, and the complement system in the nemertean *Lineus ruber*. The presence of the complement system C3, Factor B and complement receptor proteins indicates that complement could be activated by the alternative pathway; whereas the presence of C1q and FreD-C proteins with characteristics resembling to those ones that activate complement system in vertebrates, suggests that this system could also be activated by the lectin pathway. Moreover, the upregulation of genes belonging to the Toll pathway and the complement system after exposure to *Vibrio diazotrophicus* suggests that these pathway and system could be involved in immunity against gram-negative bacteria in *Lineus ruber*. We demonstrate the presence of the Imd pathway in spiralians, identifying the Imd protein in *Lineus ruber*. However, expression levels of *imd* were not affected during *Vibrio diazotrophicus* infection, suggesting that this pathway might not be involved in defense against gram-negative bacteria in *Lineus ruber*. Additionally, lectins, probably not involved in complement system activation, could be involved in immunity (e.g. *fred-c5, c-lectin2*). Overall, our results demonstrate the presence of immune pathways involved in defense against gram-negative bacteria in *Lineus ruber*. However, further research is necessary in order to elucidate the role of these pathways in response to other pathogens (e.g. gram-positive bacteria, fungi) and other putative roles in the organism.

## Material and methods

For more details, see the supplementary materials and methods.

### Animals and bacteria

Adult *Lineus ruber* (45) were collected in Bergen (Norway) and kept in sea water at 12**°**C in the animal facility. After oviposition, the offspring was cultured in the same conditions. 40 and 60 days old juveniles were fixed with 4% paraformaldehyde for 1h at room temperature and stored in methanol. The bacteria *Vibrio diazotrophicus* were purchased from ATCC and cultured in Difco™ Marine Broth (Fisher Scientific).

### Bioinformatic survey of immune genes in *Lineus ruber*

Hmmer profiles were built from alignments of the protein of interest using HMMER software v3.2.1 (www.hmmer.org) and blasted against the *Lineus ruber* transcriptome and the *Notospermus geniculatus* genome (50). The sequences obtained were validated by BLAST (74) (www.blast.ncbi.nlm.nih.gov). Protein domain architectures were analyzed using SMART (75) (http://smart.embl.de), hmmer (76) (http://hmmer.org), and NCBI Conserved Domains (77) (https://www.ncbi.nlm.nih.gov/Structure/cdd/wrpsb.cgi).

### Gene cloning and whole mount *in situ* hybridization (WMISH)

Fragments of each gene of interest obtained by amplification of cDNA libraries using specific primers were inserted into pGEM-T Easy vectors (Promega, USA) and transformed into competent *E. coli* cells. WMISH were performed as described elsewhere (78).

### Immune-challenge experiments in *Lineus ruber*

Immune-challenge experiments were designed using a similar approach to other previous studies in which immune-challenge experiments were performed (72,79–85). 64 animals were randomly distributed into 8 groups of 8 animals each. 4 groups were exposed to *Vibrio diazothropicus* (10^8^ bacteria/ml), while the other 4 groups were used as controls. Prior to infection, both control and immune-challenged animals, were injured with a sterile needle in order to facilitate the penetration of the bacteria in the infected animals. At different timepoints of infection (3h, 6h, 12h and 24h), the animals were frozen in liquid nitrogen and stored at −80C.

### RNA extraction, DNA synthesis, qPCR, and data analysis

mRNA extractions were performed individually for each animal using TRI Reagent™ Solution (Thermo Fisher Scientific) and 1-bromo-3-chloropropane (Sigma). cDNA was synthetized using SuperScript™ III First-Strand Synthesis System (Invitrogen). qPCRs were performed using specific primers for each gene (Supplementary Table 2) in Roche LightCycler 480 real-time PCR machine using Lightcycler 480 Sybr Green I Master kit (Roche). *Actin* was used as a reference gene (Supplementary Figure 1). Expression of each gene of interest was compared between infected and control animals for each timepoint and the fold expression was obtained using the ^2-ΔΔCT^ method (Supplementary Table 3) (63). Data was analyzed with the Light Cycler 480 SW 1.5.1, Microsoft Excel and StatPlus:mac LE v7.

## List of abbreviations

CCP: Complement Control Protein domains
*Cfb*: *Complement factor B*
*Cfc*: *Complement factor C*
CR: Complement Receptor
FBG: Fibrinogen domain
FreD-C: Fibrinogen-related domaincontaining proteins
MBL: Mannose-Binding Lectins
MreM: MASP-related Molecules
PAMP: Pathogen Associated Molecular Patterns
PGRP: Peptidoglycan Recognition Protein Receptor
PGRP-L: Long Peptidoglycan Recognition Protein Receptor
PRR: Pathogen Recognition Receptors
qPCR: quantitative real-time PCR
TLR: Toll receptor
TrypSP: Trypsin-like serine protease domain
vWF: von Willebrand factor
WMISH: whole mount *in situ* hybridization

## Declarations

### Ethics approval and consent to participate

Not applicable.

### Consent for publication

Not applicable.

### Availability of data and materials

The datasets supporting the conclusions of this article are included within the article and its additional files. Sequences obtained in the genomic/transcriptomic surveys are available in the Supplementary Figure 1. NCBI accession numbers of genes for which their expression was studied by WMISH and qPCR are the following: *TLRα3*: ON146341; *TLRα4*: ON146342; *TLRβ1*: ON146343; *TLRβ2*: ON146344; *imd*: ON146345; *C3-1*: ON146346; *freD-c1*: ON146347; *fred-c5*: ON146348; *c-lectin2*: ON146349; *c-lectin3*: ON146350; *c-lectin5*: ON146351; *c-lectin9*: ON146352; *c-lectin10*: ON146353; *c-lectin11*: ON146354; *c1q1*: ON146355.

### Author’s contributions

AH and AOA designed the study. AOA performed the genome and transcriptome surveys, the domain and phylogenetic analyses, gene cloning, probe synthesis, immune-challenge experiments and the immune challenge experiments, including the infection of the animals, RNA extraction, cDNA synthesis, quantitative-real time PCRs, and data analyses. AOA also wrote a draft manuscript. AOA and AB performed the whole mount in situ hybridization and imaging. AH and AOA revised and contributed to the writing. All authors read and approved the manuscript.

## Acknowledgements

We want to thank to Carmen Andrikou for instructing and guiding AOA in performing the immune challenge experiment and for discussions. Furthermore, we would like to thank other former and present members from the Hejnol lab for helping with the *Lineus ruber* collections.

## Bibliography

1. Nurnberger T, Brunner F, Kemmerling B, Piater L. Innate immunity in plants and animals: striking similarities and obvious differences. Immunological Reviews. 2004;198(1):249–66.

2. Honti V, Kurucz E, Cinege G, Csordás G, Andó I. Innate Immunity. Acta Biologica Szegediensis. 2015;59:1–15.

3. Zhang Z, Long QX, Xie J. Roles of Peptidoglycan Recognition Protein (PGRP) in Immunity and Implications for Novel Anti-infective Measures. Critical Reviews in Eukaryotic Gene Expression. 2012;22(3):259–68.

4. Aderem A, Ulevitch RJ. Toll-like receptors in the induction of the innate immune response. Nature. 2000;406:782–7.

5. Mayer S, Raulf M-K, Lepenies B. C-type lectins: their network and roles in pathogen recognition and immunity. Histochemistry and Cell Biology. 2017;147:223–37.

6. Hoffmann JA, Reichhart J-M. Drosophila innate immunity: an evolutionary perspective. Nature Immunology. 2002;3:121–6.

7. Valanne S, Wang J-H, Rämet M. The Drosophila Toll signaling pathway. The Journal of Immunology. 2011;186:649–56.

8. Nie L, Cai S-Y, Shao J-Z, Chen J. Toll-Like Receptors, Associated Biological Roles, and Signaling Networks in Non-Mammals. Frontiers in Immunology. 2018;9:1–19.

9. Peiris TH, Hoyer KK, Oviedo NJ. Innate immune system and tissue regeneration in planarians: An area ripe for exploration. Seminars in Immunology. 2014;26:295–302.

10. Gerdol M, Venier P. An updated molecular basis for mussel immunity. Fish & Shellfish Immunology. 2015;46:17–38.

11. Mapalo MA, Arakawa K, Baker CM, Persson DK, Mirano-Bascos D, Giribet G. The unique antimicrobial recognition and signaling pathways in tardigrades with a comparison across Ecdysozoa. G3 Genes|Genomes|Genetics. 2020;10:1137–48.

12. Orús-Alcalde A, Lu T, Hejnol A. The evolution of the metazoan Toll receptor family and its expression during protostome development. BioRxiv. 2021;

13. Toubiana M, Rosani U, Giambelluca S, Cammarata M, Gerdol M, Pallavicini A, et al. Toll signal transduction pathway in bivalves: Complete cds of intermediate elements and related gene transcription levels in hemocytes of immune stimulated Mytilus galloprovincialis. Developmental & Comparative Immunology. 2014;45:300–12.

14. Li X-C, Zhu L, Li L-G, Ren Q, Huang Y-Q, Lu J-X, et al. A novel myeloid differentiation factor 88 homolog, *Sp* MyD88, exhibiting *Sp* Toll-binding activity in the mud crab *Scylla paramamosain*. Developmental & Comparative Immunology [Internet]. 2013 Apr;39(4):313–22. Available from: http://dx.doi.org/10.1016/j.dci.2012.11.011

15. Lemaitre B, Kromer-Metzger E, Michaut L, Nicolas E, Meister M, Georgel P, et al. A recessive mutation, immune deficiency (*imd*), defines two distinct control pathways in the *Drosophila* host defense. Proceedings of the National Academy of Sciences [Internet]. 1995 Oct 10;92(21):9465–9. Available from: http://www.pnas.org/cgi/doi/10.1073/pnas.92.21.9465

16. Bao Y-Y, Qu L-Y, Zhao D, Chen L-B, Jin H-Y, Xu L-M, et al. The genome- and transcriptome-wide analysis of innate immunity in the brown planthopper, *Nilaparvata lugens*. BMC Genomics [Internet]. 2013;14(1):160. Available from: http://bmcgenomics.biomedcentral.com/articles/10.1186/1471-2164-14-160

17. Zhou Y-L, Wang L-Z, Gu W-B, Wang C, Zhu Q-H, Liu Z-P, et al. Identification and functional analysis of immune deficiency (IMD) from *Scylla paramamosain:* The first evidence of IMD signaling pathway involved in immune defense against bacterial infection in crab species. Fish & Shellfish Immunology [Internet]. 2018 Oct;81(July):150–60. Available from: https://linkinghub.elsevier.com/retrieve/pii/S1050464818304145

18. Kaneko T, Yano T, Aggarwal K, Lim J-H, Ueda K, Oshima Y, et al. PGRP-LC and PGRP-LE have essential yet distinct functions in the *Drosophila* immune response to monomeric DAP-type peptidoglycan. Nature Immunology [Internet]. 2006 Jul 11;7(7):715–23. Available from: http://www.nature.com/articles/ni1356

19. Myllymäki H, Valanne S, Rämet M. *The *Drosophila* Imd signaling pathway*. The Journal of Immunology [Internet]. 2014 Apr 15;192(8):3455–62. Available from: http://www.jimmunol.org/lookup/doi/10.4049/jimmunol.1303309

20. Zhang S-M, Coultas KA. Identification and characterization of five transcription factors that are associated with evolutionarily conserved immune signaling pathways in the schistosome-transmitting snail *Biomphalaria glabrata*. Molecular Immunology [Internet]. 2011 Sep;48(15-16):1868–81. Available from: http://dx.doi.org/10.1016/j.molimm.2011.05.017

21. Gerdol M, Luo Y-J, Satoh N, Pallavicini A. Genetic and molecular basis of the immune system in the brachiopod *Lingula anatina*. Developmental & Comparative Immunology [Internet]. 2018 May;82:7–30. Available from: https://linkinghub.elsevier.com/retrieve/pii/S0145305X17305979

22. Bajic G, Degn SE, Thiel S, Andersen GR. Complement activation, regulation, and molecular basis for complement-related diseases. The EMBO Journal [Internet]. 2015 Nov 12;34(22):2735–57. Available from: https://onlinelibrary.wiley.com/doi/abs/10.15252/embj.201591881

23. Merle NS, Church SE, Fremeaux-Bacchi V, Roumenina LT. Complement System Part I - Molecular Mechanisms of Activation and Regulation. Frontiers in Immunology [Internet]. 2015 Jun 2;6:1–30. Available from: http://journal.frontiersin.org/Article/10.3389/fimmu.2015.00262/abstract

24. Ricklin D, Reis ES, Mastellos DC, Gros P, Lambris JD. Complement component C3 - The “Swiss Army Knife” of innate immunity and host defense. Immunological Reviews [Internet]. 2016 Nov;274(1):33–58. Available from: http://doi.wiley.com/10.1111/imr.12500

25. Girija V, Gingras AR, Marshall JE, Panchal R, Sheikh A, Harper JA, et al. Structural basis of the C1q/C1s interaction and its central role in assembly of the C1 complex of complement activation. Proceedings of the National Academy of Sciences [Internet]. 2013;110(34):13916–20. Available from: http://www.pnas.org/lookup/doi/10.1073/pnas.1615704113

26. Clas F, Loos M. Antibody-independent binding of the first component of complement (C1) and its subcomponent C1q to the S and R forms of Salmonella minnesota. Infection and Immunity [Internet]. 1981;31(3):1138–44. Available from: https://iai.asm.org/content/31/3/1138

27. Matsushita M, Fujita T. Activation of the classical complement pathway by mannose-binding protein in association with a novel C1s-like serine protease. Journal of Experimental Medicine [Internet]. 1992 Dec 1;176(6):1497–502. Available from: https://rupress.org/jem/article/176/6/1497/24746/Activation-of-the-classical-complement-pathway-by

28. Matsushita M, Endo Y, Fujita T. Cutting Edge: Complement-activating complex of Ficolin and Mannose-binding lectin-Associated Serine Protease. The Journal of Immunology [Internet]. 2000 Mar 1;164(5):2281–4. Available from: http://www.jimmunol.org/lookup/doi/10.4049/jimmunol.164.5.2281

29. Ehlenberger AG, Nussenzweig V. The role of membrane receptors for C3b and C3d in phagocytosis. Journal of Experimental Medicine [Internet]. 1977 Feb 1;145(2):357–71. Available from: https://rupress.org/jem/article/145/2/357/22132/The-role-of-membrane-receptors-for-C3b-and-C3d-in

30. Rosado CJ, Kondos S, Bull TE, Kuiper MJ, Law RHP, Buckle AM, et al. The MACPF/CDC family of pore-forming toxins. Cellular Microbiology [Internet]. 2008 Sep;10(9):1765–74. Available from: http://doi.wiley.com/10.1111/j.1462-5822.2008.01191.x

31. Prado-Alvarez M, Rotllant J, Gestal C, Novoa B, Figueras A. Characterization of a C3 and a factor B-like in the carpet-shell clam, *Ruditapes decussatus*. Fish & Shellfish Immunology [Internet]. 2009 Feb;26(2):305–15. Available from: http://dx.doi.org/10.1016/j.fsi.2008.11.015

32. Wang L, Zhang H, Wang L, Zhang D, Lv Z, Liu Z, et al. The RNA-seq analysis suggests a potential multi-component complement system in oyster *Crassostrea gigas*. Developmental & Comparative Immunology [Internet]. 2017 Nov;76:209–19. Available from: http://dx.doi.org/10.1016/j.dci.2017.06.009

33. Gorbushin AM. Immune repertoire in the transcriptome of *Littorina littorea* reveals new trends in lophotrochozoan proto-complement evolution. Developmental & Comparative Immunology [Internet]. 2018 Jul;84:250–63. Available from: https://doi.org/10.1016/j.dci.2018.02.018

34. Sekiguchi R, Nonaka M. Evolution of the complement system in protostomes revealed by de novo transcriptome analysis of six species of Arthropoda. Developmental & Comparative Immunology [Internet]. 2015 May;50(1):58–67. Available from: http://dx.doi.org/10.1016/j.dci.2014.12.008

35. Gorbushin AM. Derivatives of the lectin complement pathway in Lophotrochozoa. Developmental & Comparative Immunology [Internet]. 2019 May;94:35–58. Available from: https://doi.org/10.1016/j.dci.2019.01.010

36. Azumi K, De Santis R, De Tomaso A, Rigoutsos I, Yoshizaki F, Pinto MR, et al. Genomic analysis of immunity in a Urochordate and the emergence of the vertebrate immune system: “waiting for Godot.” Immunogenetics [Internet]. 2003 Nov 7;55(8):570–81. Available from: http://link.springer.com/10.1007/s00251-003-0606-5

37. Suzuki MM, Satoh N, Nonaka M. C6-like and C3-like molecules from the cephalochordate, amphioxus, suggest a cytolytic complement system in invertebrates. Journal of Molecular Evolution [Internet]. 2002 May 1;54(5):671–9. Available from: http://link.springer.com/10.1007/s00239-001-0068-z

38. Nonaka M, Kimura A. Genomic view of the evolution of the complement system. Immunogenetics [Internet]. 2006 Sep 9;58(9):701–13. Available from: http://link.springer.com/10.1007/s00251-006-0142-1

39. Dunn CW, Giribet G, Edgecombe GD, Hejnol A. Animal phylogeny and its evolutionary implications. Annual Review of Ecology, Evolution, and Systematics [Internet]. 2014 Nov 23;45(1):371–95. Available from: http://www.annualreviews.org/doi/10.1146/annurev-ecolsys-120213-091627

40. Hanington PC, Zhang S-M. The primary role of Fibrinogen-related proteins in invertebrates is defense, not coagulation. Journal of Innate Immunity [Internet]. 2011;3(1):17–27. Available from: https://www.karger.com/Article/FullText/321882

41. Yang C, Wang L, Zhang H, Wang L, Huang M, Sun Z, et al. A new fibrinogen-related protein from Argopecten irradians (AiFREP-2) with broad recognition spectrum and bacteria agglutination activity. Fish and Shellfish Immunology [Internet]. 2014;38(1):221–9. Available from: http://dx.doi.org/10.1016/j.fsi.2014.03.025

42. Zelensky AN, Gready JE. The C-type lectin-like domain superfamily. FEBS Journal [Internet]. 2005 Dec;272(24):6179–217. Available from: http://doi.wiley.com/10.1111/j.1742-4658.2005.05031.x

43. Lu Y, Su F, Li Q, Zhang J, Li Y, Tang T, et al. Pattern recognition receptors in Drosophila immune responses. Developmental & Comparative Immunology [Internet]. 2020 Jan;102:103468. Available from: https://doi.org/10.1016/j.dci.2019.103468

44. Pees B, Yang W, Zárate-Potes A, Schulenburg H, Dierking K. High innate immune specificity through diversified C-type lectin-like domain proteins in invertebrates. Journal of Innate Immunity [Internet]. 2016;8(2):129–42. Available from: https://www.karger.com/Article/FullText/441475

45. Müller O. Vermivm Terrestrium et Fluviatilium, seu Animalium Infusoriorum, Helminthicorum et Testaceorum, Non Marinorum, Succincta Historia. Copenhagen, Leipzig: Heineck and Faber; 1774. Vol. 1, Part 2.

46. Medzhitov R, Preston-Hurlburt P, Kopp E, Stadlen A, Chen C, Ghosh S, et al. MyD88 Is an adaptor protein in the hToll/IL-1 receptor family signaling pathways. Molecular Cell [Internet]. 1998 Aug;2(2):253–8. Available from: https://linkinghub.elsevier.com/retrieve/pii/S1097276500801367

47. Shelton CA, Wasserman SA. *Pelle* encodes a protein kinase required to establish dorsoventral polarity in the <i>Drosophila<i> embryo. Cell [Internet]. 1993 Feb;72(4):515–25. Available from: https://linkinghub.elsevier.com/retrieve/pii/009286749390071W

48. Li S, Strelow A, Fontana EJ, Wesche H. IRAK-4: A novel member of the IRAK family with the properties of an IRAK-kinase. Proceedings of the National Academy of Sciences [Internet]. 2002 Apr 16;99(8):5567–72. Available from: http://www.pnas.org/cgi/doi/10.1073/pnas.082100399

49. Ghosh S, May MJ, Kopp EB. NF-κB AND REL PROTEINS: Evolutionarily Conserved Mediators of Immune Responses. Annual Review of Immunology. 1998;16(1):225–60.

50. Luo Y-J, Kanda M, Koyanagi R, Hisata K, Akiyama T, Sakamoto H, et al. Nemertean and phoronid genomes reveal lophotrochozoan evolution and the origin of bilaterian heads. Nature Ecology & Evolution [Internet]. 2018 Jan 4;2(1):141–51. Available from: http://dx.doi.org/10.1038/s41559-017-0389-y

51. Dziarski R, Gupta D. The peptidoglycan recognition proteins (PGRPs). Genome Biology. 2006;7(8):1–13.

52. Kleino A, Silverman N. Regulation of the Drosophila Imd pathway by signaling amyloids. Insect Biochemistry and Molecular Biology [Internet]. 2019 May;108:16–23. Available from:https://doi.org/10.1016/j.ibmb.2019.03.003

53. Dushay MS, Asling B, Hultmark D. Origins of immunity: Relish, a compound *Rel-like* gene in the antibacterial defense of *Drosophila*. Proceedings of the National Academy of Sciences [Internet]. 1996 Sep 17;93(19):10343–7. Available from: http://www.pnas.org/cgi/doi/10.1073/pnas.93.19.10343

54. Shin SW, Kokoza V, Ahmed A, Raikhel AS. Characterization of three alternatively spliced isoforms of the Rel/NF-kB transcription factor Relish from the mosquito Aedes aegypti. Proceedings of the National Academy of Sciences [Internet]. 2002 Jul 23;99(15):9978–83. Available from: http://www.pnas.org/cgi/doi/10.1073/pnas.162345999

55. Keshavarz M, Jo YH, Patnaik BB, Park KB, Ko HJ, Kim CE, et al. *Tm* Relish is required for regulating the antimicrobial responses to *Escherichia coli* and *Staphylococcus aureus* in *Tenebrio molitor*. Scientific Reports [Internet]. 2020 Dec 21;10(1):7013. Available from: http://www.nature.com/articles/s41598-020-63872-1

56. Ahearn JM, Fearon DT. Structure and Function of the Complement Receptors, CR1 (CD35) and CR2 (CD21). In: Dixon F, editor. Advances in Immunology [Internet]. Academic Press, Inc; 1989. p. 183–219. Available from: https://linkinghub.elsevier.com/retrieve/pii/S0065277608606549

57. Vorup-Jensen T, Jensen RK. Structural immunology of complement receptors 3 and 4. Frontiers in Immunology [Internet]. 2018 Nov 26;9:1–20. Available from: https://www.frontiersin.org/article/10.3389/fimmu.2018.02716/full

58. Helmy KY, Katschke KJ, Gorgani NN, Kljavin NM, Elliott JM, Diehl L, et al. CRIg: A macrophage complement receptor required for phagocytosis of circulating pathogens. Cell [Internet]. 2006 Mar;124(5):915–27. Available from: https://linkinghub.elsevier.com/retrieve/pii/S0092867406001243

59. Punnet R. Linneus. In: LMBC Memoirs on Typical British Marine Plants and Animals. 1901. p. 57.

60. Beckers P. The nervous systems of Pilidiophora (Nemertea). Zoomorphology. 2014;134(1).

61. Gąsiorowski L, Børve A, Cherneva IA, Orús-Alcalde A, Hejnol A. Molecular and morphological analysis of the developing nemertean brain indicates convergent evolution of complex brains in Spiralia. BMC Biology [Internet]. 2021 Dec 27;19(1):175. Available from: https://bmcbiol.biomedcentral.com/articles/10.1186/s12915-021-01113-1

62. Martín-Durán JM, Pang K, Børve A, Lê HS, Furu A, Cannon JT, et al. Convergent evolution of bilaterian nerve cords. Nature [Internet]. 2018;553(7686):45–50. Available from: http://dx.doi.org/10.1038/nature25030

63. Livak KJ, Schmittgen TD. Analysis of relative gene expression data using real-time quantitative PCR and the 2-ΔΔCT method. Methods. 2001;25(4):402–8.

64. Sastry K, Herman GA, Day L, Deignan E, Bruns G, Morton CC, et al. The human mannose-binding protein gene. Exon structure reveals its evolutionary relationship to a human pulmonary surfactant gene and localization to chromosome 10. Journal of Experimental Medicine [Internet]. 1989 Oct 1;170(4):1175–89. Available from: https://rupress.org/jem/article/170/4/1175/24167/The-human-mannosebinding-protein-gene-Exon

65. Ichijo H, Hellman U, Wernstedt C, Gonez LJ, Claesson-Welsh L, Heldin CH, et al. Molecular cloning and characterization of ficolin, a multimeric protein with fibrinogen- and collagen-like domains. Journal of Biological Chemistry [Internet]. 1993 Jul;268(19):14505–13. Available from: http://dx.doi.org/10.1016/S0021-9258(19)85267-5

66. Li H, Zhang H, Jiang S, Wang W, Xin L, Wang H, et al. A single-CRD C-type lectin from oyster *Crassostrea gigas* mediates immune recognition and pathogen elimination with a potential role in the activation of complement system. Fish & Shellfish Immunology [Internet]. 2015 Jun;44(2):566–75. Available from: http://dx.doi.org/10.1016/j.fsi.2015.03.011

67. Fujita T, Endo Y, Nonaka M. Primitive complement system - recognition and activation. Molecular Immunology [Internet]. 2004 Jun;41(2-3):103–11. Available from: https://linkinghub.elsevier.com/retrieve/pii/S016158900400080X

68. Albertí S, Marqués G, Camprubí S, Merino S, Tomás JM, Vivanco F, et al. C1q binding and activation of the complement classical pathway by *Klebsiella pneumoniae* outer membrane proteins. Infection and Immunity [Internet]. 1993;61(3):852–60. Available from: https://iai.asm.org/content/61/3/852

69. Poole AZ, Kitchen SA, Weis VM. The role of complement in cnidarian-dinoflagellate symbiosis and immune challenge in the sea anemone *Aiptasia pallida*. Frontiers in Microbiology [Internet]. 2016 Apr 22;7:1–18. Available from: http://journal.frontiersin.org/Article/10.3389/fmicb.2016.00519/abstract

70. Clow LA, Gross PS, Shih C-S, Smith LC. Expression of SpC3, the sea urchin complement component, in response to lipopolysaccharide. Immunogenetics [Internet]. 2000 Oct 5;51(12):1021–33. Available from: http://link.springer.com/10.1007/s002510000233

71. Wang G, Zhang S, Wang Z. Responses of alternative complement expression to challenge with different combinations of *Vibrio anguillarum*, *Escherichia coli* and *Staphylococcus aureus*: Evidence for specific immune priming in amphioxus *Branchiostoma belcheri*. Fish and Shellfish Immunology [Internet]. 2009;26(1):33–9. Available from: http://dx.doi.org/10.1016/j.fsi.2008.09.018

72. Peng M, Niu D, Chen Z, Lan T, Dong Z, Tran T-N, et al. Expression of a novel complement C3 gene in the razor clam *Sinonovacula constricta* and its role in innate immune response and hemolysis. Developmental & Comparative Immunology [Internet]. 2017 Aug;73:184–92. Available from: http://dx.doi.org/10.1016/j.dci.2017.03.027

73. Doolittle RF, McNamara K, Lin K. Correlating structure and function during the evolution of fibrinogen-related domains. Protein Science [Internet]. 2012 Dec;21(12):1808–23. Available from: http://doi.wiley.com/10.1002/pro.2177

74. Altschul S. Gapped BLAST and PSI-BLAST: a new generation of protein database search programs. Nucleic Acids Research [Internet]. 1997;25(17):3389–402. Available from: https://academic.oup.com/nar/article-lookup/doi/10.1093/nar/25.17.3389

75. Schultz J, Milpetz F, Bork P, Ponting CP. SMART, a simple modular architecture research tool: Identification of signaling domains. Proceedings of the National Academy of Sciences [Internet]. 1998 May 26;95(11):5857–64. Available from: http://www.pnas.org/cgi/doi/10.1073/pnas.95.11.5857

76. Finn RD, Clements J, Arndt W, Miller BL, Wheeler TJ, Schreiber F, et al. HMMER web server: 2015 update. Nucleic Acids Research [Internet]. 2015 Jul 1;43(W1):W30–8. Available from: https://academic.oup.com/nar/article-lookup/doi/10.1093/nar/gkv397

77. Lu S, Wang J, Chitsaz F, Derbyshire MK, Geer RC, Gonzales NR, et al. CDD/SPARCLE: the conserved domain database in 2020. Nucleic Acids Research [Internet]. 2020;48(D1):D265–8. Available from: https://academic.oup.com/nar/article/48/D1/D265/5645006

78. Martín-Durán JM, Vellutini BC, Hejnol A. Evolution and development of the adelphophagic, intracapsular Schmidt’s larva of the nemertean Lineus ruber. EvoDevo. 2015;6(1):1–18.

79. Deris ZM, Iehata S, Ikhwanuddin M, Sahimi MBMK, Dinh Do T, Sorgeloos P, et al. Immune and bacterial toxin genes expression in different giant tiger prawn, *Penaeus monodon* post-larvae stages following AHPND-causing strain of *Vibrio parahaemolyticus* challenge. Aquaculture Reports [Internet]. 2020 Mar;16:100248. Available from: https://doi.org/10.1016/j.aqrep.2019.100248

80. Lu Y, Li C, Zhang P, Shao Y, Su X, Li Y, et al. Two adaptor molecules of MyD88 and TRAF6 in *Apostichopus japonicus* Toll signaling cascade: Molecular cloning and expression analysis. Developmental & Comparative Immunology [Internet]. 2013 Dec;41(4):498–504. Available from: https://linkinghub.elsevier.com/retrieve/pii/S0145305X13001870

81. Lv Z, Zhang Z, Wei Z, Li C, Shao Y, Zhang W, et al. HMGB3 modulates ROS production via activating TLR cascade in *Apostichopus japonicus*. Developmental & Comparative Immunology [Internet]. 2017 Dec;77:128–37. Available from: https://doi.org/10.1016/j.dci.2017.07.026

82. Ren Y, Pan H, Pan B, Bu W. Identification and functional characterization of three TLR signaling pathway genes in *Cyclina sinensis*. Fish & Shellfish Immunology [Internet]. 2016 Mar;50:150–9. Available from: http://dx.doi.org/10.1016/j.fsi.2016.01.025

83. Russo R, Chiaramonte M, Matranga V, Arizza V. A member of the *Tlr* family is involved in dsRNA innate immune response in *Paracentrotus lividus* sea urchin. Developmental & Comparative Immunology [Internet]. 2015 Aug;51(2):271–7. Available from: http://dx.doi.org/10.1016/j.dci.2015.04.007

84. Wang M, Yang J, Zhou Z, Qiu L, Wang L, Zhang H, et al. A primitive Toll-like receptor signaling pathway in mollusk Zhikong scallop *Chlamys farreri*. Developmental & Comparative Immunology [Internet]. 2011 Apr;35(4):511–20. Available from: http://dx.doi.org/10.1016/j.dci.2010.12.005

85. Zhu F, Sun B, Wang Z. The crab *Relish* plays an important role in white spot syndrome virus and *Vibrio alginolyticus* infection. Fish and Shellfish Immunology [Internet]. 2019;87(October 2018):297–306. Available from: https://doi.org/10.1016/j.fsi.2019.01.028

