## Supplementary Figure 1 for "The Toll and Imd pathway, the complement system and lectins during immune response of the nemertean *Lineus ruber*"

>MyD88 Complete protein sequence
MASSNPISDADEELEELLRGAPVRCLNSSTRSRLSLFLNPTRSIPARCGYLPDWRGVAELRDFQYQQIQNFQLSPDPT
GRLLDEYEFGESPDNSVQSLLDILKTLERFDILEEMKRDIERDVNKFRRSQQRVQVPTVTSNQPDPLPFIDDFLVPEDR
FIVVDDSEDNVTIFDAFVCYTKDPSATNDRDFVKEMVRILEGQFGLKLYIMERDLLPGTNELCVSAALIEKRCKRTILVIS
QAYLRSPECEVHTRSALCFSPTCHNKKILPIFIERNVKAPSVLRPMNAVDMTMRDEQIMEWSWRRIAAALRMPLILTHE
DKRDEKKLEMISFKLRERPKLDSLEKTNEQAQKSDSTV*
>Irak Complete protein sequence
MSLAITKPSQWRPRGKKRNNSIKFLSDLPFSVRHKISKSMDSGPHPGYARLAGCLGYSQLDIANFALEFKKMDGSPM
MALFHDWESRNPTMACVFRYFNMAERYLEMDILKRFVPEHYRSEVPNNSDAIDVDAMFDSDDKDDAAGGKDVKPDL
NENESKKLPEEWKGRPGMPRENSWDDESNDHPNEKRKISGEFDGSSINADPKVSENKLSPTAPPIQDLDKIKGLSLD
TKKDSMSTGQCPNSLEKCTDPPMSSGSGSPDKACSQEPSLKNENGTNGSKDHQAKLVKLSFATVHSFLYEELKAGT
NNWSKSCKIGEGSFGEVFMVNNLGNTSTTFAVKRMKPSGALGIPEKSSEKFDHFITEITESRKYNHKNLILISGYALGP
ELCIVYEYMSKGSLMDNLKCKGDLKPLLWKDRIAILRGAARGLNYLHVLDEKRPLIHGDIKSDNILLNQHFEPKISDLGL
AKHALRANEMGEYTHVTLDNTYMHRNSAYLPKSFVQHPTLKRLSIKTDVHCFGVVIMETLTGLEAKTRKPFSQSMQLD
LLVDIVHDCKDRTERLHLLDRKPSDFYPGELLDDLFKLAMKCTSDKKKEQPCMKEVLEAIKDIEKRKVDIDGKIRQGVR
LEKSAPPAVKNNPLDVQLIRHIDRQQAIMTTSRPDQSLQLQQSFQPGIPSFNAGPAASGYSKSLPVRTQQTSAAPSGG
MSHYQYSLRSDTSDSVDMYQSSESMKCSSGMRQFQGVVGGNTPRTPLPLEDCSVPLPTTNCPQTHGKIRTVTGINH
VPPGRKSLDNGDQQQLHGHPPDNRQERRSPRYVSLPAMNENSPNNEHVKCSLPSKLNDEVQQQISPSLVNRQDIH
QQNFRPTGAKSLPVMKNDKPAGSEPTSYSLPPNALPPTQGQFPSAGDSHCEAPPGRKARHQCEISEQEILEIKRKKP
LSFLSDVSDGLFPFPDEYESIDGEDSHEEEEWDEDEDANKQLPSLPPGLSRPPTFQINPPRACEESDERDGHISVSDD
DMVVPIAQPGISVPPPGGADKMNREGGENQLTQEFETITLSSLSDNGEHESLTLPPAIYREPTS*
>Dorsal/Dif/NFkB-p65 Complete protein sequence
MLGKQGMSAASIAAGSGRGGRNKSNGKMNSQVPVLNNDPWVQITEQPKSRGLRFRYECEGRSAGSIPGERSQIER
KSFPAIKIHNYQGTAVVVVSCVTKDNPPKSHPHSLVGKGCDKGVYTQKIKSHDMVATFPHLGIQCAKRKDVDGALKQ
REGIRVDPFSTGFKHKANQIDLNVVRLCFQVFLPDSNGRFTRIVAPVVSNPIYDKKAAGDLVICRLDKHAGSVRGGEEV
FLLCEKVNKDDIKVRFYEEDEDGIPIWEGWGEFGASDVHRQFAIVFKTPSYRQRNINKPANVWLQLIRPSDEETSEPK
QFIYKPEDHDPEYILAKKKRKVASSLDPLFDPAHLASIPSTVPHGEPSAGDTEMMEISNPSVKERVKLKASRNIQTKKE
DSAQSTSAAGMDQYPLYDMSGTTDAALVQQMVHTQGYQIQAPPGGGGDLYYSYPPGDVQTVSTEGMTHQQILDSL
AQGGIEQVNSSDSAELRRFVQENFPNMQSIDAEGNISLPGSLLASMEPLDVAAHVDVSNVGEQTAQFHNLQTVNTTQ
QQS*
>PGRP-1 Complete protein sequence
MSRTACVDVTRQGWHARPPKSTSPLNTPVGMVFIHHSAMQRCSNFDECAAQVRAIQNFHMDVRGWDDIGYSLLVG
EDGKVYEGRGWDRIGAHTYGYNDVAFGICVMGNFTTVTPNSQALDAVRNLIACGVRDGKVTTDYKLYGHRDSKKSQ
TECPGQAFYNVIRGWPHYDTHGPPPDPQPIGAK*
>PGRP-2 Partial protein sequence
IDVVKGIQNYHMNTKGWHDIGYSFLVGDDGFIYEGRGWKTQGAHTSGFNNKAIALSFMGDFTSQTPSTMAIAAAKAMI
AC
>Fadd Complete protein sequence
MDGKDYSEIDKVKEYREMLVKLSRNIDAKAFKELKHICDVKKARAEKMNDALDMFEYLTERGELSANNVANLIELLEKL
ERIADAQLVKDFRNKFGGVKAQYYNGSLKGTENITEKTITQPHDEVDACADSNLNAAFEFLREQLGHKWRELARTQG
LSQVDIHAIEHAYQRDLKEQSYQALLLWRKKQKQNATIASLEADLRKRHFNQMANDIASQKYLSV*
>Imd Complete protein sequence
MALMQNPKQSVEVAKDPKPSVLEVDSVVIGSSSEIGDGDGEIGDVKAGLDCLGANAGDQDQGTGSDIGELTQRTSQ
MSVRPKDKQPELNNVLPGADSHDEIDAARAIPFNTYGAGGQANAVGEETKTIAVMKQGTKEIKYTERKRRIHQYDKPA
QYPIPAFNNCQFNSAVITGGTFGDFNQTYQSMHERNQPALDVLDNHVVTDHAVSLSDEFADLLNCVEQISIDDILDIAE
EIGGNWVALGRKLETKTATMDNIKCQCHYVDQQEMCVLLLEEWQERVSEDAVISEVTKALIDLKCKGEDVVPAILEKL
SDRHKCQ*
>Dredd Complete protein sequence
MQADAVGPSEDFRLTIKQISDDLDSDEVLTLKYLCNDYIPSKRLEEIKTGLDLFDVLCNKNLVTAEDTMYLAECLYSIQR
MDLLLGHLRMTANTVRQMINSSRPLLDPFRVMIMNVIECMDSSEMRELAFCLDEYLPKKYTKYTSGYTLLMALERSGQ
LAPDNLHLLERLVEKTVEDENALIAIRNYKASRHETSSKVPGDTTTHQKLVNNLVIVPQAPPRLVVVQNGTGAGGTGG
PRQPLPVDSGQKLGQQPFDDPINTSPSDATPGGASGDHQVETPCSAEAVGGKVHLPNDPCCGEVDQKLPSLLDEPG
GVACSKQPVPGEEDYRKIGGATGGVAFSKLPVGLPGEEESENMEEATGGGDLEPPDKKKQQENSTSSFEGATMAT
GRQGDEVGKTNLYLPPSKQWTPAETAAIQTKEEPMLPSFLNEPEHVGSVASQAGELGPAPPDNEPVVEEIEAALPSIL
ANFQDTDDEEPHPTEAADEDSIRVPPTQSDLPEGLEDGPNIAPGGDYIRAKTFEEMERELQTLREEIYKRENDMPSYK
INANPRGVCIIINNVNFCAVPGGKRLKDRHGSDVDQENLRFIFNKLKFVIKPYKNLTRDEMMGVVKQAAHQTDHSKYD
CFAFCVLTHGSKGNLYGVDGLPADVMTITDNFKGLNCPTLQGKPKLFFIQACQGHINQIGFKTRMEEDSDNNYGETPS
LLPNEADFLVGYATLPGFVSYRSETHGSWYITMLTDMLDKYSDRHDLLSIMIKVNEEVAKASAMIEGGIYKQSPMPWP
TLRKKVFLKVSD*
>Relish/NFkB-p105/100 Complete protein sequence
MSDSDDSIELGDGGAKVTHKISKGVTSVGDPYIEITEHPQSRGIRFRYKSEGPSHGGLQGESSTRGRKTYPSIKIHNYE
GLAMIFVTLVTDQDPPQPHAHELVGKNCKDGRCGFEIRINRAMNITFPNLSVQHVTKKSVLKTLEQRIRENRYPFQEIP

QTLQDEEITSIQEEAKKRANSIPLSVVRLQFEVLLKPDGKAHRRLPPAYSRPIYDSKSPAAAALKICRMDRQAGCCRGN
EEIFLLCDKVQKDDIRVRFYEEDHNGDVKWEEEGNFAPSDVHRQYAIVFTTPPYHNQNIDQPVEVKIQLVRKSDEEFS
DPKPFTYCPQRDKEEIQRKRRKKIQVFPSYNDGGAEGEDRENYQDSNVRSVRTLGSGGSASTNVPTRQNSLPNAST
ARLPTVLDSNVVADCAAVFERRQELPTKRLIGHKMNKVDSYDGPEEADSYTGPNIEPEFKVEPFKLRVDHGNRESRV
MGEKDVGCQTELHPEEELKDRYTANYKGTLAYSIAERSTAGLLQYAVTSDAKSLLHVQRYLLGVKDEEGDTPLHIAIV
NKNKDVAKAMVEVAATIQDADFLNTTNTDGQSAVHLAVIYKQHEVFDSLLRCHADPSIQDKDGNTAAHIIAMEDDKKC
MEVLLKYRDNKLYKDNVVDMSWANKKNYEGLTPLHLAVRSRSSFCVKALLKAKVDLNIEDAKSGRTALHFCADVNNIA
ILGVLLTEGNPDVNARDYDGLTALHIAVSRGNIGMAAMLMLRGADPTIESNRTAENSPKDDEGEEGATAQPDEVRGET
PMDFAKQLKNVENDKKKLIRVLEGESYTDVSSVAKGMKEAAAVKFNPYDDLPPIDSLYHSLTDLGVHVLPQGDIVDLN
FGVRYKLALVLDKDNSARGWRGLAEKIELDALIEPLGREKSPTMALLDNYEVIDGTVAKLREALKAINRAELIKVLDQYE
KIIASKRRSKPSGPKSLDSGYTSVAASASATKSLPTVTGNGNKSRKD*
>Factor B-1 Complete protein sequence
MSRVVYLGLLSLAYILQSYLAVGVCNNPPPLQNGRVRLLDGWGHTRMRAVYECNQGYFLAGPRKRRLAVDTFTREC
VRGGWLMPSELPRRLLCLLKSKCHANPTSPGNGQMVGHCCDNLDTVTFTCNQGYKRAGAEILQCSEGRWNGSAPV
CTSGQVCRSPQKPNGIVESRRSGQTFIPNDAISYKCDTDYYFEKKVEREDDYDSYYDEVSEYADEGAEFIIYDDIARKI
KCLGNGQWSMSFPDCHAFKCEPPAPIRNGFIDFQGPESSQYDGGRVISYSCGQKYILVGPSHRKCSMKAGGWTGN
DPQCIQIRCPTPEAPEHGYIDNTRLNKEGDTIHIHCIRGYTLQGSSSMTCGAYADWSGTTPTCHKADIGVGRCQAPVIP
FGGRKRNSNSGNSYQLNETVKYSCYGSNRLIGPEEVRCTSEGWMPNRQPKCQVPFSYDNPAELAQKFTRLFDKLEN
KTSEIENNVIGRGRYLDVNSETGLDIYFLIDSSASVTETYFRKAVNFVKALVKKVGVTSQPNGARVGVTTFSDDVAKQF
GLNNGFHTSQNKPLEKVVELLDNLKYSQGSTNIRGALEFVRNQMLPMSLASRTGAKQYLFLLSDGDDNMGGDPIPEA
KELKKSTKGYPDMEIFCIKIGLENDSEKKIKNEKKLREISSEIEMNKDGTDPHFFRLKDFAAFDILVETIVQRSPDYSECGI
AGVKTGISSANIINGEVAFEHAWPWMASILEKNENEAGLGAFKHTCGGTLINSEWILTAAHCFQKELYEKNLDKIRVKL
GSNLWKLESDTDAIEVKSMHIHRGWKVDINKMHPSHATWENDIALLRLAASARLSNKTRTICMANNSDEHNALFYSGA
AKGVVTGWGFYQNRSVGTDEKPKSRSDNLRQVELRVVQKDSEPSCKPPAKEKIVCAGSSGLADSCFGDSGGPLMV
STARDANGERRWYQIGIVSYGKGCGLFGDQIQYGRYTDVRHYQDWIKNTIKNVTESANPSI*
>Factor B-2 Complete protein sequence
YAPRQQLNERTPREHKGTTAMSSMFLAITAILALTVTSTQANCPNPLPPSNGLVSPETGQFTPGNIVRVFCRPHYKFD
GTNETMRVIRCLPGGVWESYPQCRATITCSTPPPKAGGTWILTGRGPGDRHSHTRARANFQCNPGHIFAGPRNNSV
TQARMQDLVWYCNNDLQWAGRWIRRMNMPGYRCLAEADCNSDPGDITHGSRTGSCCANNNAITYSCDNGYHLIGR
KTITCINRKWNTKPYCSASRVCTKHDLAIGTLVEYPLESDKFEPGTTAWFFCEHGYFFRRIETEVTSDDSYDYGASAT
QTKVTFDTGNRSLDCTGPRWSGGPTPACNHVFTCPVPTVTNGQWNTNQPRIKHGEAVTITCNPGYIELGPASIKCSID
KANDKDGWNLDPPSCRQIRCARPVRPERGIVSLKNGTDIRAIGAVIEISCNHNFELDGSSIRKCMTNGRWSGNDATCQ
PTRAQDISHCPVPVKPYGGRKFGSSYNKGNVVTYECHNDHISRGPSRIVCSDAGQWKPVGLPRCEALFSFDDREQL
AGNFGQVLDAMVKSDEYDPIRGRRIELNAPEGLDITFLIDSSGSITQKDFNDSIEFVKLFVEKLGVSEKSNGARIGVASF
AAEPEREFHENNGFNANPPKSVELVKNLLDNLKYQSGDTNITGALAHFKSNMLPLNLEKREKAKQYLFILSDGDYNRG
GSPVLIAEKLRKGENEVEIFSVKIGSDKETAEAKKNTLVLIEMSSEKEMNGAGTDPHFFHLQDYATFDYMVQELLNGSI
DYSECGVAGKTGITLGNIIGGNLAQPRAWPWMAAIMASQADNLEFKHHKQICGGTVLNGEWILTAAHCFQGDTYNNIK
RVVVKLGTNNMKQFQESNPINVQDVRVRSMHIHQEWIVDAVDKKKYDNDYSNDIALLRLARKISFSNRIRPICLIPKTTP
NKGITNKIAVVTGWGRPGNESLPADQRSEKLRSDELRQVRLPVVTQEEDPRRCINRYPKEKVICAGWKNKVMDSCW
GDSGGPLMIELGTENNKPHWNQIGIVSNGIGCAQVDGLTDAAYGRYTDVRQYRDWIDATIQHAGA*
>Factor B-3 Complete protein sequence
MMLLALLLVGTIIEGSVGAPSRSDDGRITCVMKNFHIGEGANVIASVFAGETMTLKCRKGYYFDRVLQDIDYSEDITSTG
NDYFYDTAPRVLACQGDGRWDKPIPTCHAFTCPRPDTPSHGAIRPDRQTIFSANDTITYVCEKGTVLVGYSSQTCDM
AWSGWSYGSKPPNCITVRCPSPEPPRNGWITNRNRDIDSTIHYYCNRGYERYGAPNQTCQANGKWSETTPICKAAE
RQVRACENPVVPPGSITRGNSILIGEKVTHRCYWPEFLKGPDTRTCGPLGWSPSSEPECEALHAFDNRDTLAKTFFSI
VDRFSSSSEQNTINGLSSRRYDLFFLIDSSHSVSPDHFTLAKNFVRHAIDKLYLTTSGTAIGLASFSTKPEVEFDRSGGI
HADPGRAKPLIFEKLENLKHIDRDTNITGVLEFVRKVMIPMTSNARGAEDVPKVIFLFSDGDATCGGNPIQSAKKLKDL
GVEIFVIKIGTVRNDKTRSILEGVSSEADFPRHYPHVLRIKDNSDMEYMVANFKDYSTARTIFPQFPTCGVAGDVHYTK
SYQIGGDAANSTTWPWIMAISKIHSDNTAEFACTGTLISDQWVLTTVQCIDDRTGAHFVATSASSPDKFDGEIVKQTFR
YGTMPSDVALLRLQHPLKFGPYVRPICLPNFRPTDELDDKFGDHAVTIGWNQIAKRPKLIQAAVPIVKDWRCRGLAPG
AGYCAGLNTSVNGTCTQDIGSPLMIKSRMDRKTREVHWIQIGIAWQSDGCTATGGNTGRFNRLERFVQWMKTTMNS
TTN*
>Factor B-4 Complete protein sequence
MSPRQLIAMFVVCLLFVVGLLTSHASAACDDYNYADDCNLDATCYRPLIVHGDSKGSDGTSKDIYNIGERMSVVCNED
YLIRGDQEATCQGDFTWSWSVEPKCVPKPPPAAECTTLKEMASGMYALVNRTEKIGSIYSKIYVNCYVDEDYYLHGP
GVYTCEGTTWKGNNFCLKRRVCPRITAPVNGFITIGGLESRRKGDEIHFGCKQGYTLMGDRRWKCEESLRWSGYNV
FCEKDLGIAGMARGIQENIIDKFKQQTVSASNRGRLSASNVNGIDLFFLVDRSDSISAGDFKLAQDFINMLINFFGISSK
GGATRLAVISFGTKAKLELSFSMIYTIEECLIKAKQNGENIQDHHAPWATKCLNVTEPSLIGYNPKCLGLIPEDNSTAINN
TEIAMSLKELVKRKINLIQSSGGGTNIFEAINMVNRIATRCDAEKAVFLMTDGRDNMITDLKLDAVLHRIRRSVNEIFGIGI
GQDLRMDQLKAIASPPIARHVFRLRDYVELNELKDLILTTENDYSACGVSNVQRTGVVVLNSTIVQEAKQRSWPWLVK
ISVRQGAFICAGALVAKNYVLTAASCLHDENGTLLQRVRSDSPISVMAGYYDVNDMSGQQTTSAEELKVYTRFDIRRP
RENDVALIKLHDDVKFDPFVRPLCLLGSTDSILFHQSDRKWLVGWNTSNTESLMTAKPWERTMELHGKKTCSRELPY

TTSHSYLQCMGDRHQKRGYCAGDKGAPIVVQARDIDGKPIYKVVALVEDGSECMRPGKFTVVTQLTPGFMAWINTNI
QN*
>C3-1 Complete protein sequence
MLLFQALGALFAAAFVAAQQQPTVFVVFPNLMRFDSLEEVIIDYSSGAGADSASCSVSLQNYPDRDNTFSSKDVVVRS
GAQEKVYVITKLTDLVRGSLSKLNYVYVRVQCNSLGFNKEAVVLLTGTKGYLFLQTDKPIYKPKEEVAIRAISLNPDMLP
ANTDFTMDVLNPNQTIFQRFEGKAEKGFFSKKIKLPDVTIQGNWSVRASYKNQFQPQSLISNVSFEVREYVLPTFSVNI
KTPNFILRADLEVQVIVKAKYVFKEDVRGRLTIVFSLVKKDGSSQTITIKTYSDFHGEKTVALPKDDFKNKWFPDVGSRL
VVDATVTEEASGNQETTSDSSAMFVNSPYVVSFERTTKYFKPGLPYNLKLALKYASGGVAGGVRMEVSATAKETGR
NDRVNVFRNDAESEQFITDEEGWKLITLDVPANAEDLKVKVNTTDQHAASTGEITLKPQAAARNNFLSIQLINNPSRVP
VGQKLEIRVDFTENVNKHVTFMVLTRGKVVSSYSTPVNGISKHENIDITPQMAPKARLVVYYVSSGNPPEVIADSIVLDV
SKTCKEELELSHPLGGGNQDYRPKENVKLTIKSATDSMVSLLAVDKAVYLLSERDILSRDKMFNTIMDLDLGCGPGSG
SNVGEVFRDAGLYVLTNRVNVGTPRRMSPECKQKIRKRRAAERVRRGLHRNNRIIATLNKYSNAARACCEDGLDQQV
QQDQRSCAAIAQTMVHGSCRNAFLHCCQLEKDANTPSGRSGADDEEEINENDIDIDRREDFPETWFNELLEMTRDQ
EILNRALPDSITTWVISGVSVSASEGMCVAKPINVTAKKEFFVHADLPYAAVRGEQLEVKATVYNFGRNRITTDVWME
GNRNKLCFVGNSKGRTRKQRVVLEPKTGKKVTFPVVPLEVGDIDITVTAMTQFTGDQVIRKLKCAPEGVTRYATFSTR
LDPEGIMGGSGSGGSSAKVTIDIANKRQTIEIPLKFPGNAIPGTKVCSVSISGSLLGPMVPSQDLNMANKIMQPMGCGE
QTMMFMGPLVYTTRYMLRTNTLSGELEQQAYKYILSGFQRELTYKHPDGSYAAFQHRGSSSWLTAFVAKVFCQASQ
FPNVGKAVAPHAMAAIKWLLGHAQRADGSFYEKLSVIHTDMFGQSRKNDVTFTAFVVEAISKCGNVNGIDITSGDIATK
LGKSVAYLVQNRNSQNKPYEQAILTYALALQNNPHSVAANMKLRGMSNLDPVTGMRYWNPDGSVVPENTTPTWYK
VKPSAYAIETASYALLAQLAIGGAAEDYFQYSGPIASWLIRQRTKTGAFVSTTDTVIGLEALSDYALRVDSRCDLNMHV
RVFAGNNLNKNFQLTKADAYKPYTITENLPIDGKLFMVATGTGIGQVQVECQYNSPEIRGEVCTFDLKVTSEEGNAPV
EELNNEIEQVKQDRADGIPRDVGGVERFVNEIKIDFSYIGAGAKVGMSILDVTLFSGFSPVLIDLKTLERNGVIDRYEVV
SSSVLMYIGEVKKGVTTTVTFRAKQDQDVKDVQPAAVKLYDYYEPDKHHCTKFYHPKQNSPLLATICDKNDEGRKVC
QCAASACTEFWGFPQGKRPHLKTKDELINDACNIYEYAVKIRAGPTEVCGPLSKMEATVEINIKAGIVVYSAGDKIQFA
WNAKCENPSFTAGNSYLVMGKDGAEYVDENGRGSMRYVLDSSSHVVEWYG
>C3-2 Partial protein sequence
EGYKMLNPPLLLGAFVLLAATYLTDASSNLFVASPNLLRVNSDEDVWVTLHHDQNTQATVKVYLENYPDRGGRFSEK
TITVGKDESKKVTMRVGWDNIPQGSQQTYVYLVAECQTLGFKKESVILLSKKSGYLFLQTDKPIYNPHEKVNIRVVSLD
AEMKPSDLEITLDVINPNGDVIKRYLKQQATNGFVNVEFNLPKILVYGNWTLQASYKDRKQPETTKTSIEFEVKEYVLPT
FSTKVKTLDDFILESQSEIQYKITTNYVFGTPVTGRGKVYYYLVYPGGNEHQIGTLEFDVTGDYVGSIKKGELGNRWFP
EEGTRFLIKVAVTEGATGKQEKASDSSAVFVKSPYVISYERMPKFFRPHLPFTVKVDLTYPNKAVAAGIPISIEALVTVG
GRQLGAHGNQVTTKDYVTEENGRQDVTLDIPDNAADLKIKVTTHDPDLAVLHQAEKPILLAKHTSGSNEFMVLQRRTT
GREIKINHNLEIRADFTNNRASKKVNFMVVARGKILQHFVKLVNDRTTHENIQVSPLMSPQARVIAYYFTGNEIVSDSLV
LDIVKSCGNELTITTNKNEYQPKDNVEITIAAGKRSTVSVLGVDKAVYLLSEKDILTRSKMFKKMLEMDIGCGPGGGKD
TADVFKQAGLSVLVQSGLIAIPRRNSEKCPQNMFRRRRDVMETYSDIMARLSDQDKVCCNRGKADSVDATRSCQQL
TSQYGSSRRSCSAAYLECCVAETRRVSGGDRGRSGGDGDQVQLGSFENVQVDIREDFKETWINELINFGPNEESIN
MRLALPDSITTWVISGVSVSPDQGMCVAVPTNITSRKPFFVHVDLPYAAVRTEQLEVKATVYNLGERDLSRVLVSLKA
DKSICYAAESGKYTYRHEISVKAKSATSVYFPIVPLKANDIPSEISVKALSAFGGDEVIKKIRVVHEGLKTFSTQSVRLDP
EEIMGDAGSGVRGWEVDINKAEKRQTVKIPLKFPENTIPGTEKCSVSIHGNLLGPMVPSGDFDISKRIKLPAGCGEQN
MITMGPLVYATRYLLRIGKLTGALEADAYKFITKGYESQLTYKKKDGSFAVWQPARSSSWLTAFVTKVFCQASTFDQI
GQKVREHVVGAVNWLFKNAQDPQSGKFVETYKVYHREMQGQTLTNELTLTAFIVECISSCKEVINLSDREFAGPMQK
AVQYIERNRDTLLGNPYGQAILAYALTMTNSQYKTEANQLLWDMSSFDTVTGMRYWNLDGQAVDHSAPVWYTKPPS
AAAIETASYALLTQLAISDEERYFTNSGPIASWLIRQRTKTGSFSSTTDTVLGLEALSEYAFKANSRSTLDMRIRFYASGI
NFGVNENLELKEASALIPILIENNLPIGDILNVEATGTGMAQLQVECTYHAPPKDDQGCPFTLEVDVTEYKATSADLGPL
DAVPAQSLPDRSRFGGFRFKRADDGGTSSYVATIQSRIRYNGGQKVGMSMLEVSLFTGFEAYLDDLKKLKKIGKVDD
YEITGSSVKFYLKEISNIEDTFVKFRATQKYKVGKTQPVAVKVYDYYEPNSRQCTKFYHPHAGSALLAQSCEESQGTK
VCACVAGTCAEYWGFDRAHIPRGKRFKLRTRTELYQDACNIYDYVVKVKAGQRFINKPLVEMQAEIEHVIKLGRDNFD
TGETINFVWNSECTYPLMEDGVSYLVMGKDGKEYIDKNNRAKTKYILNSSSHVVQWYGQERRGYMDALNAFAGKLL
REGCGQ*
>C1q-1 Complete protein sequence
MKKTLLLVLLFICVVSSRKEGRKKQRHGGDNTLNDEGGSCDLEISCKNTDGSFPVKLPIKGPRGPPGSAGSRGDKGD
PGEPGLPGLPGKSAANTQKVAFFAGLKDNVGPIKKDVDVSFDKVMTNVGGAYDAKSGKFTAPVNGTYTFTVVVAAQ
GRQKAAVKLIRNHDMIATVWAESIPYWASASNTAVLHLKKGDQVWLQVLQRASYLHGYMYSTFSGHILFED*
>C1q-2 Complete protein sequence
MCRMKIYKFQIITFLSLCLLTIGGMGITNVLGPDGKTDKNTKDKSCISDTCKCFQGPAGAPGVPGVPGMHGTRGRDGS
KGEKGNFGQPGLPGDVGGPGQPGTRGKKGNKGEKGDTGEQGYTGNKGEPGNSGYNGLKGAKGEPGQSAGAKIA
FSASRGKKLGPVLQDTIVTFDKIFSNVGDGFDVYTSHFVCRHNGTYIFMTHILGQDNRDAFAWVMVNKNHQLPLHGD
GRAGYGTGSNTIILHLNVDDHVWIQLNKNSALLNDYSTFSGYLIFED*
>C1q-3 Partial protein sequence
RREVRMTTRRLVPLLLLAVFLFANATHNNKHGRQQREAAKDADQGTCQQCQCGPGPPGPAGPAGMPGNHGNNGN
NGAPGAPGMPGEKGEQGAEGIKGKRGSKGEPGTPAKGPLKIAFSVQRNLTYGKFGYDFDISYDHVITNVGEAFSSYT
SHVTAPVDGIYVFMFNTLGGNGNTLVSLMVNGNKRASAWGYHTDKNPYPNAGNQIVLQLLRDDRVWLKLAAG

>FreD-C1 Complete protein sequence
MKTFACLAFFALAGCVLPLHHAVQACRAPQACAIKPKRIHNGIFARVSDPNQNDAFAFSCNNGFALHGKAIVTCCNGT
WDAFPTCVAIDCKAHYDSGNTKDGAYMIYPGGQATPILVYCDMTNGGWTVIQRRKDKSVDFFRTWADYKAGFGQYN
VNFWIGLDNLHKLTVQNSEIYFFMKASSTENRYALYSTFRVADEARKYRLSVSGYSGTAGDSMKHHNNMMFSTVDAD
NDLYTDSCAGLYKGAWWYNTCHHSNPNGLFLNGPHDTRYAEGIEWYTYKGQHHSMVYFEMKIKPNMLPWGS*
>FreD-C2 Complete protein sequence
MKWTFAIAVLSSALFVISSASSSFSQTTTLIDQWTRTHSGCKCSQDDQSMACPCCVEGACQCGVFSGSRSGKCARC
DSTSFISDCSTTTSQSDNQPTDSNGWTGRMECPCPEKTPTRPTTCACCVNNGCRCNDDPYQCTQCGREDECLVKP
ATNSMSSPGDTSPHFSYAHIDSQSYPPRGDTPAGTGGGDDYDQSSDYRQPTTGGDDDKDQTSDGGNGQTNGGSN
NYGNYDYGAGGDGYDQQRVTRPPLFLSTSPTSSRPHVYTSKFINQIGLKCVTVTTCYKPTMTTMATPTTTTTTPTTTP
TITTTTSITTSPTTLPPTGPTWIDRPNGDCSTLAATGTGNGVHEIHPSRREPFDTYCLDGWTVIQSRLGPEEDFDRDW
QDYKTGFGDPSTGNYWVGLENIYHLTNSRRYRLRVVLKTFGHLAYWIEYDHFFIESETEGYRLHIEGLSQGNAGDGF
RSRGFGSRRFHNGMKFSTKDRDNDNSGISCARKYRSGWWFNNCFNANLNGMYMSPRRKGGRYAFCGENGCVAW
LPIHNRLSMEKVSMMIKLEEDDTVPRCGGQSDRFGQGNRLRKVRQWKQRPGCDWRRFWVDSAPGQSAQEHGKH
GWRLRRLRNGS*
>FreD-C3 Complete protein sequence
MAIRVFLAMAVSVCVVATTGRKGDNTCAYTFQVWKPDQDVISILDRLDLKQRNQSMHFNTELLRLRNDVLHELLASKN
ISHSIEKQVMEQKLENVKLRMQQQEIKMSIMSMKDRNEFVGGSRAGRVPNQLGGEDSMQNEEQIKTDVGSLQAESA
WLKNSIGNLEAEWILIKREIQELRNGGTKTTTDTGNIGTDIAELKSMMHNVQADTLLLKKQQNDLKKENIRLNAMILRIEN
ASDDYLKFQTRMRKSTSRIQEQLHEMRFENSDLQHELITLKSTNRKLRDRLGESANNITDPYSSKDGKSVYPDISTGT
QSDRAASLTLRSTDTPRDCDEIYKRGYSISGVYQIRPDRSPHLDDVYCEMFNGSGWTVLQRRVDGLQKFNRRWLEY
KFGFGNNYGEFWIGNEKLTQLTNQKKYVLRVDMWDWEGKRYYAEYDHFLVESATEKYRLHVSNYHGNAGDSLSYH
NDMAFSTGDMDNDLHARNCADENQGGWWYQSCYTSNLNGVYRTSWYSGGGQKYANGIVWYTLKESDLYSLRKVE
MKIKPYRR*
>FreD-C4 Complete protein sequence
MLEKMAGKCSILLVLHVLIWTAYGQGGVSQKDPGKRLQEVEDRIQWLTKEHRNMTNVVAKIHHRLHTTHHKMTEIKEE
ELLRGKLITDMRDQLEKQSYAIDRLQQDNQQMRGVILQLSQAVQNIRGQPLGTTLAPIVKTTKSTLPTTPRDCQDVFN
GGSHYAGNYYITVEPTGTKSSFKVCCQMTDGAGWTILMRRMDGSIDFFKKWDEFKRGFGNLEAEFWLGNDKIHELT
SQGDYMLRIEMESWEGKKYYAEYDHFSIADEADKYRIHVSGYHGNAGDSLTSYWEDHNNQAFSTRDRDNDDRFYD
NCAEHYHGAWWFKSCFESHLNGIYYRKGEHNNYFVRNGIQWNTIHLHSSLKYAAMLVKPSTKASDPSRRNDTVD*
>FreD-C5 Complete protein sequence
MEKVLEYIIFLNLAVFITSLQASGYQRSYSLCTNIEYHYMSPGLQTRYQCQMFKRIMEEIEGDRVHSENVLSHLQTLLYS
IDQKIDTMKSGISQDIYQSAAMRDFPRVSSNIISFQNNRDVPLLEDCSDLYEYGVRISGVYPIKITDGSIVIVYCDMETDG
GGWTVIQNRIDGQVNFSRNWEEYTHGFGNPNSEYWLGNTHLNKLTISRNYTLRIDIGDWENGTAYAEYTNFRVEGIE
ENFRLYINGYRGTAEDSFKGYHNGMQFSTQDSDNDLWFGHCAEKDGAGWWYRSCGYSSLNGRYYHMGKIPLGPD
GMVNRGVLWHSWKQDYLYSLKSSTMKIRRKRKS*
>FreD-C6 Complete protein sequence
MELRLSSRHIGCTILIVLCFLSAHVSATYYYPYYNYYHRQRPTKCAYSLVVNEVSGSKCPELMQRLQADQATARLQKE
AAPRSETESETKKIREELRNFEVTFYKEMSRLKDLNMTVKIQQDAMVDTQKYVRKKVLNATRFDLLIKNMDKKIEKQW
TYIQDLESKVFEVMTSLGELNLMLQKRLPKLNNLDTKIITVESAAKVKNCGLGANTKYYRDCLHIYLSGYRKSGIYYVTPI
YSACPIPVWCDMDTPGGGWLVLQRRKDGALSFDRTWLEYRGGFGDLNKEFWLGNDNIFLLTNQDTFRLRVDLWDF
KGSRVYAEYERFIVDGERTNYRLHIGNYSGTAGEGLLRHDGADFSTPDRDNDVWEKYHCAKEWGAGWWFTNCWF
AFLNGLYSNSTNVKYRGVAWNDWKNEQLMAVEMKIVPIL*
>FreD-C7 Complete protein sequence
MKLLDTILFILSTTYMQTLASSSGDQRDKRAGPITKEEAFYTMVEGLHRQINDLYKDRDIDRKHFQELEDKVYHHQKTI
HKLRVELHGLNEQQTTNQDKIQNLQGLIENQESNFQNIRTAQTLLGQEIKGLLEVQANFTENVSRLEQIVLKRAEEKEN
FSDAVYDMNKEIKTICDSIQRQKSDLEQLQFTFQHVIEAEFGKPKHKSKSKTRSKKKGEKKPTSIPILDELSSSGSGDLE
LDVKVTESPKTDKSNQTVVGDLDLNSNDIDSGSGDLDLDPDLEKVNPVTPTQETPATPTMFTSTDSEDKVDITNNGIL
HVFSNDFAERFIEQVNAIDDLKVKVKTLNITVKNLEEQFLEIPTERAQTGADLFRDMFSNLTQQMLTLQQLNTVNIPNIT
NTFSNQKELDKLLTVVMETKNSINDLARDAFDEKIQNEAKLSKIDFKIFQMNKTVSTAVRRLEKYKERTRDTNDDFGDQ
IAETSEKNLKLEQRLAHVEVNVFNASLQKCLQKDRDNDQDKQIVNIIQDLQTLITVSEKQRRDLRRLRVLVEENNMNND
LTTKVIGDLLGLTKNLTIHIPELLDAKAEVDNFVEHLPLDCTEIYHKGIRHTGIYLIRPLQQRGSVGTYCDMDNDDGWTLI
QRRLDGTTDFNRDWYDYKFGFGDTKSDFWIGNDNIYYLTIQDDYMLQVDMWDMNGVYWYALYENFYITSSEQDFILT
VNNYTGNATDSMSYSNGMKFGTPDVDNDASSTHCASFYHAGWWYKHCQFSNLNGRFDIGMVWFNYEKDEWMQM
KKTTMKIKPRNSSD*
>FreD-C8 Partial protein sequence
TNHSIWAFYDDFKVKSETEKYRVTFTPDEFNKQRWRGDAIMPGQADVDMNNTEFSTYDNDNDGNSAENCATTSKAG
WWYGPGCTNANLNAPYYNKSYWNGLLWPQITWGTVTNTSQGFSRVTMCLQPYDVEFGDKILVI
>FreD-C9 Partial protein sequence
NESENYKVTLTPSSQNKKTWHGDAIMPGDPDLDMNNKLFSTFDGDNDGGPDNCALNRKAGWWYGPGCTDANLNA
KYYNATNFQGLLWPQLSWGTMTNASDGIKFTVLCLQ
>FreD-C10 Partial protein sequence

DKPFLARCDQRTTLIQAQGNDRAPLNNMNRTWMEYAQGFGANFKDFWLGNNYIHAMTSGDRPFTLEVEFRAVIGNG
KSSNVQKYWYVNFKVANESEKFRVRADRNPNIMGV
>FreD-C11 Partial protein sequence
AEAQKAKEAEAQKAKEAEAQKAKEAEAQKAKETEAQRVKDNEAQKVKDAEAQKAKGAEGKKEAGEAAKPSKATPTV
NSPAIVNGERILLYQKGVTSGMKDCFSHFVNGRKRDSGFMIHPLGMNGQPVKVWCDMTSGGWTVIQKRLSGSLDFY
KSWQDYKEGFGDIFGEHWIGNDLIHFLTNQDKYSLRIDMMDWQRNKVFILYKDFWIDSEAEGYRLHVGGYSGTVGDA
LAKHNGKKFSTKDVDNDEVGSHVKQFKEWDGSCAKRFSGAWWFYKCYQSNLNGKYYRGGDVPDKKFDGVAWKP
WKGPKYSLKRVEIKIRPKDASEVVLR*
>FreD-C12 Partial protein sequence
MSSPKMPHLPVKLAFLLVFPFFISAASAATKTKFNMFKCGDDNGIPTAYIKAKHQVRSKLNCAVLCSKYGDVCENMHY
VPSIKDSENCFIGSAVNLNRGYGTCTGLVNTTGAVSFYRKVECGGGSINQATGACECPQGYGGSGCTTRINDCHDTT
TKAWHTVWAYPPGAPVPFQVRCYSQITFIMIRNWGSQATVSFNRSYEEYAAGFGDPNKSDFYLGNKNIHYLTNWTR
GQHISLKLYRHQKAAPFTRYYNHMKVMSEADQFRITIANIQNGQATDTDGDALLSTTPGLSIKEMPFTTYDKDQDNWA
DGNCASKFGGGWWYNNCTAAPLTAKLMNSDQITAGGGPDN
>FreD-C13 Partial protein sequence
MLYTASSTYLVRTTTTGFRLDNTLVTLATPSITVTTSRRHNNQQFSTRDNDNDRFGRSCPRMYSSGWWFHNCMVAN
LNGLYYEGGEYKSHRKDGIMWQPWRYETSLKKTEIKVRPANFLKNQQKGIDVQSVKP*
>FreD-C14 Partial protein sequence
FLHIHINEIDRIFLSKMQIFVILKTLFPIFLFFRCLLGHRVVDTFGSKFFQLPDCPVDKARPAPSSNRFRVESDLECARKCL
WAYEDTCRWMNYIEPDKVTGENCELGNTPANAVCNDLEDRPGYHYHEKGDFCLNQSTFVKSSLQCTCVPGWTGQR
CNYRAFDCSDHYYVHLWNSGRQYLHIKARYSTNNYTVHCDLTTSLSGASGNTCIQRQGYSSGENFERNWNDYKKGF
GNLATDFWLGNDIISEITTKNSRKYRLTVWLQGHNNSYAVSTNYDDFLVKNESENY
>C-lectin1 Complete protein sequence
MKTKQGLGMWAFALVVLVVVIEEAAAQGVTAGIFRKPYVTPNDLLLKSLDEAKEACVAYGMVIATEDQVKALITQNGG
SYTRDPDACAWMANGKTASPGGTALANCTMATTVADVWCIEAPDYTCPKDWSWLPGSMSCFKYTATSTTFAEAKT
NCSELGGYLATIESLEEQAALEKMAVAGPAFIGISGNPAQWNDGSMRPAPLPTGIEFSNPGATDECAGVDNAAAAAK
KWTKLDCATKRPYICEMDLGMAPHYTVSFETSYVTTGAGISGNCRANIIPVVLSSELKKKYAAEKNLLCTAMTGVVVS
GPSVGIFTVQVTFPNATKAVLVQRVIETFKITSKNYALTLGCARYLEEKLPVPIPKIDAGAGCPVLEISGKRHTTLDGGPS
CDPLQGAYDMKRSCLPVDKACTAFSTKATDKGLHAEQRFIEYETSKPAATDRYVMTEEKCAAICKNYKSFTCQSYSY
TLASGVKPVKAVCKISPQRASMYETGPAAITGTSYYERVCPLFKEAPVSRRMSITQTFNYKPADKPECATSKSGADLY
VAAVSLQESFIANTNVALKAALASECGFITGMPSFKVTGFGKRIPTIDITGGSVQFNDFFSFEPTTPLVPSALISDCIAKF
TAKQDLLKPTTKYDLGPLCKKAELVPTPTNPTLVLVPTDAATDCKYADKRPFPSSVYFLPLFGTNTTCLIERPPGYKYV
KGRCTKTGTIKALDKVNRPACTAKCEAEGDTCTAFFWNPDTLLCETLQGGCTDELNVTKLEGYYTKESLTCRKSKGF
TTVSTYVKDFFAMYIGLPASKTRDELCGDESYVDIQNEFSSTTRTLKTSCGTVEMFSLEIATAVAIGYRNMTCMNGVG
WPQIGEPSTCEVKKCKELVYKYLDIDVIPALVNGNGALFTAAMKHNFVRRQCLDMMKTCSKMTQFAGVEGLSLIFTIFK
SVYPQIEQFPGPKDDCQDTFFTDEVVKFNNFKASSTKGVQIVAADLCTMFSNLKASMYLVDGCQPSVKLSSQMLIEAF
TQIMTGVCEKSTVECLKSTYMQCFIKRTEDLTLAFASGKGMESKICFNNTKENFNNDLQGCLSNGQGCTTTSVSSVSL
LYPSLVEEFFSLCDKSELTKLNVFYQLMTSQFEIEESKECLNAQAIVCLKSLNKIMIEGLTAGEDAASQICSAYESTKVCL
TTNYKKCSPITRLNLLETLMKKRYEMIAAVKVCSFTETAACTTAYQNITEEFTIATKAVSAGTYDCTKLSVLKASVATFIS
KSTCYELKTLYVQASLDTIEYFLRDRCSACAKTDALKNYIDYQNAFLDAVLNMWNYTTSSICTGQMSLTVFQSTLGDIT
KNCNVGGARLTAISQSTALYQGAFCKKKDFTWLVSVTETCKVDVVEACFKSYKEVYLVKLAAGDEQSVKQVCMLYQS
ISACVVAAIKTCNGQTQARFLAGIAYLRSMLIVTCELKPVEKECNLVKANQCLTEFRANFETYRDLVVGGKTPKAEAVC
GSAKLTNTCIKTSTLKCSDKENGMVMYFEKTFELYNLAIKKECDGLTPQQCDLSDSFKCLVTYAENVTETLMMVDRRT
AFLCNTKVTEKVAECAATTTLDCGERSTSLIATVKGLERSTEQICQQLVCRQDEAKANILFLLDASGSVNNKDKTNFGK
CKSFVKKVVEKLEIGENMIRVAGSTYASSVVRVFGFNRYYEKENVVEAVEKIVYTEGGTYTHVALANAVTDINNIKKER
GDKVPSIVIIMTDGKSKSEKKTLEAAEELKRIATVYSIGIGNNLRAREVQAMASNSNNLYTAESFDKLDDIISSISQSITCE
GFGQTPGDLKDPATCNLTEATACLATFEKELFIGFTKEGNDVLKVTCLSYKKTMACLKVALLECSGSTQLRLVEAFYLL
IQSSMEVCKTFLPAPKDTCKVTEAMECFNELKKTISKVNGAYEKVCVEINEKQNCVYDRIAGCTDFETYLWVSKIHAEV
YFVKKSKKCEGKILPPQVTGKECDDSKATKCMFEFEELAVQAAFTNSFGTLCLKLVEFKECILGSSEECTSEQTASRIN
RFNIKYALVSPDCVATTESDECKFKEIREKMEAFAKNMLGSLMLNMTTNKNAICNDLHGILQYTVDNMMKCSLVMKSQ
IDDMARMLINQLKITCPNIVYQPVVRPTCDIVDVQQTLRKFNATVFANMAAGTFEFCAVQASTKTVIDAGLKNCPDEGV
TAVTNAWATLVTKIGTSCNQRFDIDFGSTTRITVYEGETRILKYKFIGCTSALCQGEGNCQVKIFYTLTNLRGDAKCGN
KERFLVQGAFSGTSCGFDLTDQNCGNEFDLVLRSRVFFVNQKTQIREVVFYAQVTINGVVAYKKLETKIQVTFTHMRG
KPCRIWGDPHIETPDAKNPNKINKFNIFPEGLFVILKYKTVIGRNIEGIFQNCWKGRKVSCMCALYIQCGGGVVKFGEC
RRNTPPKKDPKCTKNQRKGFSVPKCDKKKPAYSPFMEVETMYNGNIDQCTIIKRSAYAYDVYQPHGRKISVSMRLPS
NLRVVEIFIYPLANDNTETEGVCRATSTTWKQFEMSSKWNSFLAAENWAAYVSPVTSPTCLCTVTGEVEKNTLQINEE
CDGFQISKECNFSKQKTTYVTKALWDTARLIGITSTARKKRAVATSPFAIDPNATPQAVSWDGSRNFTEKKATDACNK
AYDDYKAGVLRTCLAFTKQDITNDLDQCVMDIKMSDTDRFCEPSAAELWKSCNNIRSADTAYMNSPAGQAVAAQLAT
LSCPSGCSGSSKGKCVAGVCDCVVPWGGEACDVNRNDPPNIIAVGTNGGVCDPGLANCSMVMIEAANVYDNPFLTC
HFMEQKYKETGKTDVGATGTRRGELVTGSVRCILPKAVPYLIAISNDGKTTSSAYDYLVYSPKCETCDITAGTCKPVTT
PVCRVNSVCYQDGEKLKENNCFYCNVKESTTKLSISTSDRQCNAVPIKKEGLDLTDIIIISVVVAGIVIGLLALLIYKFICKP
KMDANKKAKELEKEQADNLHRSQEPQPSPEARYAEPPQGTPGSLE*

>C-lectin2 Complete protein sequence
MDFTRNILVIIGLLVIFGYEEIGAKERRDVVGCEPHWHHFHKACFHVNKSKETFDQAEKSCRAVRGSLASIHSSRELDF
VENITFSHLTSVWVGQHMVNFTVRWTDNTTLDFQPDGLTYLTAEKACMVLNEDPKGFARMDATNCSQYLPFVCRKS
ADSKCKDEDIEWLNVKGRSFPKHNIKKIDDVDNMGKCKRICLRQSEFSCLSVNYLRAKKYCILSSDTMKSSGIDLIRSR
QFTYAEWSCGDQKDKAKSLATLRLGNIRASDQEQDSDDEPPAATEPAQQKPKDAVCREHYQLMAGVCIRASFKAMT
YRDAKQHCEKEGGRMSNILRMQNIPDIGGIYRDLDRKLRKSSLTEHAFWLGATWTKSTQLVWADPDEVQSPNWSRG
HPSSKYECVIVTEKEKWETRPCDSRFSFICKRDATTTREDVEYFSLKGLFTQLANLTNARMPSCPKDFPQKRHHRFIL
PPTEGYLQNYRSRTAMCKHLYSEYRENSVFIASIRAGLWKHRKDSDPIPIIVGPDTKGRSRYRRQAQAWRLLKPDVYY
RQTGIPLPCHHLINKTIIHGLGWRNTNTMQYTWSETFTSKTFTQLRCMDEKLLRKIFLSARDLTIEELKSLPSTVYRLFS
PEFYRQLDMEHRWAYCAGLLMSKPTSHDVKYAIEDKNCRDIMTYSNMSRLLPVSVTANEYIMSLFCDYTWKERNGAS
CRLHDIEAYEFCFKIGGIRYYDLRPKCMSKQFTAILMDRVKSAISAKGQHDRLQVYAGQILLVHDFVTSSSLFCLLLDNI
DDIRVRRRLLQLPMAKDWILECDIPSNLRGSCSQTFNGFDDPRVQRVAIKGLAKVEKPISNRTTRVILALCARADCPDEI
LEKLDPEMVFENMDACRPNGCWTDPAVGAKVARILRLVRTYIRHRPLTVHDLLKLKCVLPGMGRHFLFQQKREQVVT
FAIMALQHTPLDRRTANQLARVVISREQHRLMTKKPSPNALNQIYDYFKPLKRLLPYVDPDLVDEMDPDVCSKFLPDL
AQCTDKCDAIRPEWFLAWKERAMAGRGWSLHECHEIIQSCNGGKFLSPIDIYECAKAFPSSIFHIENKKFLNRHMCYA
VLNVQSQRWKEPLEEWSARAIRQELGRLVIGLHPENITRILVRAVERDLVAKQTKPVDADDYDEDAYRGKKDLRRLRT
GRVRGKRNAEDDDDIETKEEGETALDYILLFASYFPQLRSTQRYAVMDALNDLAFSLPSDLDENSFRMIRKIVLNDGTG
SLIRSILMASRDKQETIKKICRGIKIPYIDKAFLADIISLYKKAVGTTMLDEDDILALDQCAFGLNSNDIRSLTDRGILEMCS
SGVHYYDWDDETMSMLARRTLDALHVNDVADMKMDAFRNCSSVITLIPGDDVRLIRNLKMRCQAFTVLKHGYWLRR
TTLSTRIERGELKLDVSPANAMYWYKKWEPIRKMWDGHIQQGWAASTNACSRVKDEARRMKRMKRRHRKKRRYRR
NVFDISHEKGYKMDQSQTCNDLRKLGDAVVEQGKPIFTSLRRHDFLDCLDFLGSREGYSKESVDHLAMLAKKHGGET
SSWHPQFTLELGCIAQGLSEDDLRRLPLNDSSVMESLGSFGGWSHSQLKAIHLAYLQVSGKTHSSLSGADLTRMNSI
VCGISPYDILLINPKAYRQAAGTIGEASSCTLDQQRAWAQLAKKAFGTDVTKWQRETIGRIGSVVSGLPDEDIRSLNRQ
QIDAISPKVIRALSPEIFRAFSIAQLKCFDRDQVRQINGEQLQALSREKRKTVLSLSTRRTANIYSGNDDEIMNEMAPIIS
NLD*
>C-lectin3 Complete protein sequence
MKATLWILTLHKIKMKVGLLFTLSWIILISEICECSAEANVTDLICPSGFLRFQRSCYRFSQMRASWLEAQGFCKAMGA
NLASIEYWQENRYIVGYIKKHHSKFKNKAFKRSFWIGGNDFANEREWRWVETQEKVRYQNWYHGQPDNAKNSENC
MELEKKYKYQWNDQSCQLRNHFICEIPLHVYYKK*
>C-lectin4 Complete protein sequence
MDFMLKIPAASAALVLHCFFWAFVVSRLAQGVHYCSPTPGNVVTISGESSGEIKTLLNGESSYGDKRDCVWVIDAGVD
KRIWVEIVSSKLQWAPSGAICEGYDYMTIRNGSSSTATEIISWCGNHKPHSVLSTGRYLYLQFKTNDQNIYEYAGIYLR
FGTYDHLSCPPISSKWSNGNASLCITVFQEDITWISAQQHCGWVQSNLIKIYSADMAKFVTNQIVQKTRQSSYWIGLH
DIARENSFLWIDGTKPTYNRLSGKNGVDEDDCVSVSTSSNIWSENNCDTEKPAYACSMFVGGPFPMYNIPKEGEYPE
EPQEKVSPLLIGLITGGVFILIAVIIIIGVIVYMKVIKKGRQRREKRVQEIELDNAAQSMQPESSRRSENVETENDSIFHVR
SPAMQSAGATSYSNAHTCNPTAPPASPLPAYVPNPQPSSQHIAPPGYDEAIGQPVFQYSASNI*
>C-lectin5 Complete protein sequence
MATTLAILFAVLAVQSSLADAYLPECPSDYFTIRSSCYKFIDGFKWYKDARMVCAQNMGSLMELQTRTQTQLLQDYM
QTNLRYFVDHTRTDIWLGGFKQENPHLWPERKWVWLSNTTKLLDAYQYTNWEGQKSVEEEKRYCLSATYNNLKWM
PRNCNSELSYFCQINRKHAWFTPTKERYPPCPAGFIRFGTSCYKFLFSHQRHSDARLDCSKHMGDLVKIESKEENEFL
LDYIKTNLYDVPLNGFWIGADRFDWRTKYGAVWVWLSDNKLLNQSRFSNWGSGEPHSDTSRNCLYVDHNDLKWHG
SVCNVKRQAICEIQLKADWSPNDPTKLGLRTCDAAIFEYDVSATFAMAVFALFTVYLPWS*
>C-lectin6 Complete protein sequence
MYQTRRETRRNGIQLRSSWIGSTWLVFSCAILATLDVSWACTPCHSSCRCTRKLDQTCVVDCSSMRLSSPPASNELP
ESTTISELLLQENQIVTLAPDAFKAFTNLKKLDLSSNSLTTLPDGIFQHLTPQTHLDISYNEWKCDCGLIWLQTSITTNIIIF
DRLAQTTCSKPNKLHGRTIGSLAKSELVCVPADLVSCYLDKDTAITTPVFFVDNGAPTKTTEKKCNDFCFRENHQYGII
YDNNNCACGVHAKADLPLCSCPDLLTSPEPNTGAGLCNKKYLKSVFGVSAEIRFGNTPVISTPSSFTFTVTSSLEVRQ
EDPLSYTWAVQGTYLSSALYTITRPSFTYNFLHTGAYILNVTITAGARHETASLPISVASPVKIFPDCPKYWNIDHSINAT
LSSFSASTGRVVWERSREDGAKKIHHPTCPTGSLLQGGYCFTIDTSPVSWQAAENSCVGKGGHLVALTSAELHSSLK
ANVFDTASGSEYWTGLNDRSYNGVFTWSSGGSLGSYTNWGTNQPDEANPPEDCVVLNKSGSDWQWHDRSCDAS
LPFVCQVRLGTKTRSSGVHVIGAGDFGWNETFTEVTEVAKSADSKEKTVIIFPGLWFEGIAEIRAWEFMTTTLSETAAL
RLQVYSPSCDSEEEKLVPPGCKDIASPYSICDSNPPNCPTRSDCPSYQQLCLLDNTCKNLTQPCNCQETKGYTGCSA
ATPQRHVGIIPTYQLKKEFIVELAKGSSKLIRLPISRLAKKNHVIGISYNSPDNPIKCVKDTSSSLRQNYLLSKQSEWVNL
KDAPDAFSKATWADDQVCSLRVIANTYTTSDFPSEITAKNTFAGAPGVYTFKASVGNELGSSDSECNTTVMEPVEGIE
LLSPINVAAGTDTSKLPPQILELGIPQYFLIKISRGTHVSASLKDAGMIQMPFSAKCPDTFAPSLCSKPKTDIEKPFSAVE
RTYSNLSTAPSSITFTVQNQVSSKQLVLKVKLCERISDLVIINNNGVIQANKQATFYGKISKGSCASYLWELGGVVIQGK
SSSNQLNRIFGSSGTFNLKLTASNDINTVTQTLAFHVVTYADPKDFRFISPPLEGLVGIPVSFKVSVVVNINKLFSILMDY
GDGSTKASITFQASQSPYESTQKHTFAKEGLRTVKANVSDAGSWVVAEHILPLRFPKLTVKVTTVDLNVATGKLVSFV
ANAEVETRGYNLEYTWQFGTGTAIVASKIKEYSHTFKNTGTYLVSVTVSTGFNNASDSLSIVVVEAIAGLALKTPDNIFE
TTAANFTASVTAGTGIKYYFNTGVATLPMSTSPEFVHTYPKQGTYTISVTANNSVSQEVVSKSVTVFHITYLKLIEVTGP
STCSRVNFALDFTANVVHVSPKDLEFIWEFGDGESKTTTGDPIAEHTYRSASDFTLNVTIKSAPNKSVQGTRIVCVQQ
QITKITLTPNVTVVGVDSNHPGTIAVGVAVTPASETQSFTWTFKGVSETGLPVTKKLFSLATAGKYNVSLRVENKISSQ

TQVLVLEAQTIITGVKLHNDLIETRYVKTSKNVAFSVTAVSGSDMVYSWIFGDGTTVDKVKEASQTHAYMSSGDYKAT
VKASNLVSQEDVSLDVSVQDAVTGTKLIVDTSAVAVNQQVWFTASATSGTNRSYEWKLCSSCPTITGDAVKIGNKYTS
TGTHTAKVKVFNKVSEETATVDVKVLDRISGLKIVMNVSVSTRVAKGSTVEMDATCSQGSDLSYGWIIYHTGSANTKH
AFTAKTIKFTYREIGSYNVQLTVSNALGKDDATMTIVVQETITGLDIRTTKTYTEPRELIAFTASITSGSDVSFEWDFGDG
VEKSVPQNTIDHKYASRGTFEVVLTAKNFVSTLTKSLTVGIEEKVASVGITGCCNTVLAAGVSATMSATVAAGSDLVYS
WNITGNTGKPMTFNGKAVSPTFPTNGSYTINLKVHNHISSINVDKTVNVQTKIDKVQLVLQQDKDKLFVDQMVNLGVT
TGQGSDMKYKWIVNDDVDDRLTGSSMSRKFTDQGVYRIKVDVHNDISSNTAQIEITIKQLLCKMPKLSTIPATKLEVSK
SQDFDLEVEVDTTGCTVYTVTFLWTVYASDSCDNLKSKPVVNLPAVGTTTDSALHVSSRTLSVGLYCIAVNVGYSDTP
VDLPAKFEVEVTPSPLRAVIEGGSERSHGNSATLLVESGSYDPDIWPSGTTGLKYKWTCAAVPPTPSLTNTCFKQPTD
GATINIPTASLVNRARYEFTLTVSDGGSRSKVTRQKVTVNLKPIPDVYVSCLSCRHPGSYDVLTNRPLAMKGACKNCN
GKTVTYLWTVIRSDEHRLEITGANSATGNGKPSLVINPGVLSNAYSYSVKLTIKDTVANLDGYALVQLTAAPLTVVGIDC
QIKPKVTAVLQDPVSVICTGAKASVKPNRFRVSMKWTDSANSKREIVMYYGLNPNTKVYLTHIGRKATSSVTVEVAME
NQDGHIKLIKSNVIQVTELILPSGVTRSGYLRKKVETAFTELKRKGDTVKLLQYSVALVVELNKDTGASKKEREDKMFIR
ECLLEGLHVATVTSLGRIGVMGPILDRLTSVQDEFDSKKNEVSHILSGINGYLEKHLTYGSEATLTNPNSLANAISNLLGI
YLSSSTKQQQLKEIMKNSHSIFLCYVLAKVPEESSVEFNTTEFAYSGTKTVVGSMSKSVQFKRGAIGLQPGMIPGDKG
REVARVAMANRKNPFRWGFKSGYPVTSGVLHFTLYGSDGTEIPIKDLPKEKAVTIEFVKEGSSANVARGKPDTYSPTE
NVKYTNRTLLPKDSINILVNSSKLGLTTPGAALSVQVRFTMANSESFPTAGIEVHSFLGLGFKPSRTKYTRYKSIKSSM
MVATEDYRNYTFFLNQSSFDPTKDYHILIQNNNTVSINLSVGIYVSSCQFYTAQETWSNEGCEPLDTSTSTVTVCSCN
HLTAFGASVLVPPNAIDFTDLIRIDLATNPVAIALCSAILFVYLLVLIWARRSDIRDEKKAVVVPLCGKDGPFRYEISVYTG
MMPGAGTTAQVGLKLYGEYDKSDAKHLCRPNAFQRNGHDVLVVASQRNLGDIWQIRVWHDNTGGHPTWFLSHIVIR
DLQTNSHWYFRVSSWLTINAPYGVVEKKVKAASSHELGKFSLRFRSEAASGFSEKHVWTSVFEHRDRSRFTRVQRV
TCCVTLFYTFMCVNAMWYGVLKDQDDRWKGIWEDTFTWEELVVGILSSLMVFPVNILIMQIFRKSKVKLDALKRQPKN
ETIEMDDLCDIPRQGSIIYDNNGNPCRLKRTEALMKIIREGSSSDSMRTKDTSDIGTWSSTSMSSSAKHREDLAKRLPL
NRDDPKLWSYDHILMWPEALPSWVEHENDSPILRNSRRKQSTQDSGNGSPEKDKRQPRRPSIARTARKMSIDTLLKS
CEEDLQEEEKEIKAGKRVSMADTKSTSTESTNTRKFSLTMRRSRRVSTNTSLTNGGTETIPSTFAEDDAWSSHQTEM
SEVSEFGSSYGLGSRRTTLISNGVSDTFDRRSTFCRNPSEGEFSIGREETFDLDSASMARSASIIASYGSRSFTSSSS
GPSKLNRTTSQSLIRIPEVNHRKKQCLLPHCCVYLAYSLCAALSILSIIMVILYGHKFGRDVALKWLLALFISFFQSAFITE
PLKVVFIAIFLATITKQADQRVEDNVEVSPILEYNEKIKEDKMKPIKGFALLQALDEGRKVHRMQLIVRQSIAYFLFIWLIF
VINYANHNAKAYHLTENVYNTLAKSSLPVSGRTLKSFQSQNDIWSWLETIAVPYVHSENTTDSYLYLLGVMRLRQIRAT
KDNCTVELDHIYNARFRSDGCYGKRLFDEDTASYNEMWSPADDGIWRHETTDDWTHQGIYHSYEGGGYIKSLSSSL
PASQSIITALKQGNWINDRSKAVFVDFTIYNANVDLYAVVTVLFEMGGHAGMTSSIEIHTTRFFNFSFDVFNYYDIFMILL
IPFFAYLLVHVTLTIKEERKSYAKNVWNWVEFALTLVMGASLAMYMYLLHHSAAFLAMQRQGSGFFNFGPLAYHHAVF
KRLNAVMLFILFFKITHQFRFIKIWSNYGTALGRSLARLLWVFLLFTILWLVYGMLGYLMLGPGVFAFRSFGASILSLLLTI
RGSMDFSKPYDYNPWFAGFFFFTYVFLVFGFIFGLIIAVLQKSYHLSKSRTKYPCTLDTKDYEMIDFMLERFKLWAGIN
KTQKVSRSFRKVKFKGMPSVSTRSTYSSNDASTSGSGQSASTTSFEVLERMCRESQYTDQMVERLAPSWEEVVAK
MEKLLQDDHEEAKLCSNIARRMIDSTYDTARADIPANTKAPVKPAGLALGWRSKSSIKGFIYSAESNKTPRPPSPPSQP
QGIMKRPSAIPKPVNQSVTIATRPVVWPRTSKPRSSFSTTSSSAGRSSISTISSGHRGSFSTTISEVSTGKQSSSSHSVI
SSASNASGNSTKSDSHMSVVSVRSNASSDDGRLPKSVRYSVDSKPPVPPKATQSVSASMPTPAIRNNTSARPWAAR
PRRPQHVLKSVSKNAW*

>C-lectin7 Complete protein sequence

MIFWRSVVNVELVLLVIVCCLSGRVIIVRARSKRKACDDDAPSVRNGFSRFAGDSMVYTCPRWYRFPDGNKTKSIQCT
EDGWPPIDDCMPGRCPEDWVRYRNKCYGFKKDATMNFTEALNACGSKDIKNVTLNHLATSLRNSQLERKGPVEYFK
RMIHSYIKNNRGHPQSISGQFWIGLYRDPGSKDWLWADDRQAVPVIPPERPDARYAYIDVSFHKDDANVTLAFDSGD
KNYYYMCEQHISCPGIDDNNDDLFEQRSNQLITNDRVCPQVSQLEATCFDGLWRMSSCPKPTVMPGEEVEIIIPCLPL
PSSESEQALSNTTKVGAVVVVRCAKGYEFADRTTEREYACSTKGHWVPKITQCVKIHRCPALNITGDINCTRTGEVSP
GALSDCSCPSGYMFTDYTVEKRFFCQEDASWSDVLPECVESGKYCSYPGNVFNADLLNNDTLFYVNSSVTYKCREG
YRFPDEYKSKEFTTKCLSDGRWDLHHDECHLVHCPSIPYVIMKKEEHAADMVNSSVPTFIPERIDPIYAAEVNITCLEE
NQAFPDGTITRIIQCNEDGEWQPAVPLCMPMETGPHIRIKRYVPPPMEAPGSEKIGSVAIVFIAIFFTGIVVLDLTSLGQS
CAILYHNISGCNERMVDRTEERDYEEDEKEKMNTTREDDILDVIENGQTVTAKCFDDTSLCFVDEVSDSGGEDARSD
MHENMAVLELDRHSVCTQTDDSSFTEENHVNHDTVV*

>C-lectin8 Complete protein sequence

MNRQVTCYFLVLLILEGVHLSYVCPPELCPSRSDAVASRLEGGFNPFFEGDIVLDKMDSAERWWRRQSRRRGMRSR
RETVKQKKRLWPKAIVPYTIDPRLPPTTKENIAEAMLHYERFTCLRFRLRKESDKDYIRYSYQPGCWSYIGRQGGPQR
LVLGRGCEKFGTIIHETGHALGLWHEQSRLDRDKYVKVIKDNVSPRFLKDFGVVGKKSSTSRGYPYDYESVMHYGEK
YFTINGKPTLEVIGIGKTLGLKIGQRAGLSALDAAQLGDMYQCQTKNKRVCDSTWVEYKTSCYKFFRKPPKQFGGAVD
YCKDRNSELLSIDDRDEDIFVKRYLTTHRSDTDIWRTGGRKDNYYGWIWQHSDGRSHKRMSYQKWADKEPAGGKTT
LVLKKEPNKDTFKWRAEWAGSYRQLPAYSYGFICERDTTAQGCFRSRFRDGRDYRGDINYSDQGLVCQKWTSQWP
HQHTALDENPHKNTQNGLGDHNFCRNPEGRKAKPWCYVTKQKVIWQYCNVNLCRIG*

>C-lectin9 Complete protein sequence

MIMKWILSLVAVLVVMAITESEYNTAPFKRTELICHDDDITKNEAWAYSNRSGNCYQLFNKQETFDDAKTECTKIGGSL
AEIKYEEEQLFLNVVLERDWIAHWVYGTREDGKNFTWGEAKKEMTYYNFVGNTQPTDRPTETCLVMTKNYVPPNGA
AAIFAQWTSADCNKKNNYICQKKAKRCPKLPEADHTVPKPLTPYIELTIKYSCAPGYEPPDYAARYKEWEAKIAAAND

TSLVEPTIKTKPSMEYKCSFPGNWTGSIAPCEPISCVEPPKRPHTTLNSTNRVFPSYIGYKCDHGYFYAEYNTTVMTA
WCGPDKQWIDVPKECDTVECGEFPIVFNATANAKKNHFKDTVVWQCNEGFMFPDGSVRQWVGCRENYVNNLIGR
WNVSKLDDCERLSCPPLPSPAYTFRDTNDSAYNTIVTYSCVPDDYEFPDGQPTLEVRCSEEAQWKPKVPACTKKPSK
LETKGKKYVPEPLEATAAESIGYSALIFVIIFLILIFLLDLTTIWKDIRMMKYNCRHFMKRVNSNKTPKRVNLQPIPKDMGK
DQ*
>C-lectin10 Complete protein sequence
MDYKFVLTVTIICAIAIATESKNITMRACGPRGKCTYDEIMMKVSKCFANHLTIDEEVHGKKPLTTLDALEARLTRQMEA
LSIRTLRGVRRIETKMQELTKSGMNRRLVRLDAEAQAGAQGGPQAGINEQRIISERSILCNERGESQKCPTLFTGIDG
WETCYLFSNFNATWQQARDYCAALGANLVVMETNKEHYLVNLLIKQKMTKANGTAWWTAGNYQVQSNTWMWATQ
LHLQPFRFHKWKNGTPKSETTKCMLLPGDNAHTWISDVCTEPYNFICETSKIMKMK*
>C-lectin11 Complete protein sequence
MKVLVVLTAFVALVSTNLLVKGQTNPNLILSGNECPNLWATGDQTCYRFFQNVKRTWQESKKYCEDIGAVLVNINTEN
KHQEVSAWLTGNDAQNLAWYTSGKRDETETTSPVFYWETLLPREAVKTFFFWTDPKNKDKPVPATVITYVTSGNTR
GWDLTDPSEAKAFICESPRSEVQKLSTDDRDFLFGQSDIGNIEKGPSFIQEPPRSVIFAASEAYKRVSFECVAEAKPLP
TYRWVKEEAKVGKTFEIKPNKKYTLSNGKLTIHNPQRTKQVDAQDPNAATNTDVGLYQCAATNKFGTILSNRVNFDFG
YINDFSKTKRDPIDAKLNFDTAIPCDAPPHFPSIRYQWYKDTPLEYIRTDLKPYIFISRNGKLYFSRVKNEDVGDYYCIAA
VPLPGFQESKTSMPIGLTVGTGGDERAPQIHSDFPAIYPGNPKKDDNVSIECFALGTGDLQYSWSRTNGLPMSKRVS
YMDNNRVLSINQIELEDAGGYQCTASATGGLGGTGTYSKVVKLDVQSIPFFTKPLQNHHVDVGNKFTLHCEADGKPE
VSYTWFKNGVPFNNATLEAEDRPRIKFDNTRVTIQAASAERDNGMYQCAAINQRGTSYSTAQVRVLSFAPSFAKRPM
APSTYAAIDGKATLVCTPEAAPEPIKIWYKNGVRLSVSEEPAARIHSLKNGNLELTKIVQEDEGIYRCEAENSLGKASSE
GNLLVLSKSVITVKPLSQIVRQNESVQLTCEAAVHPSLDFTYVWELNGFPIDLDFNEDWRLKQYRNLDKYERGTGAAW
GDLFIHNISYEEAGIYKCIARTPQDEAVAEANVTVIGVPGEPAGLIGIESSITSSSIQLTWSPGTNNGALLTSYTVEAFSNL
NPKWHVVKANICAWYGDPTQLTKGECREMTPLEKEESRPELLQVKGRIKTTVTGLLAYNKYNFRVFAYNTYGYGKPS
QPSSFYTTRSDKPFVIPRNVGGGGGKVGTLTITWDPLPINEHNGPNLKYRIYFKKADVSDQDKETKVEVEHPTSMYTE
TIGADNFYLPYDVRVQAYNDKGDGPVSKPKEIYSAEGMPVATPMNIRARPYNASSIEVLWDIVPDTRESVKGRVMGY
RINYWKWKIENEVKDYKRQTIVNQTDRGKIIGLSPNQYYQLNVMVFSNAGNGPKSALHEQRTWRKAPQDPPREIWVE
HLTDKSISVHWRGISIANDEEPLEGYIVRYWPAGEAMVNAIDHYAGKVVDTVLTNLESSPTRYYLRVLGFSRGGIGKM
SSPIIQFQIGPGPHLYVNPKPGSAGNLRPCLLLITLSIMALFKCLL*
>C-lectin12 Complete protein sequence
MDKLVLSFIAVWTVAWCGAHASIVRSKREISVQIGRSVYIHPDDLVLDGGTTCKVELDVFDPISQRVGTLEPKAFDCRF
QPKSVKYIHNGSPLLKRDSIKLRVHKFSNRETVTEIVYLKINILNEAYDVFVIGNPSPLAVPTFSGYSNAIDASILRFRYDM
RNNATCMVGFSHYGTNYPLVGQIVMGQDNTPIDALWRECHEFLFLGLKYQHLVPPTPDMDYIPLKVQVNDPLKGIIIER
LYLPIIITGAFPNVPPKAAFMKAYLMDVDQFVLSIVSPNIMGAEDDETPEPQLIYNISRPLGPGEGYLVNLADHTRPITSF
LQADLSNLNIAYQPPNISYTDRTNFIVEFTVYDSHFAKSAQPTVLHISVRPSETKAPRVALNAGLTLIEGHSATIRPDNLQ
IVDKDNIDRVHCLVKAGLLHGDILVNGRKQMSFSPRDIEMGHVVYYHDDTDTSRDQVLLRISDGLHSIQTTLFIHIIPKDD
SPPTLVNNIDLDVAQGELVQFKEQKLSAHDSDSNDYDIVYKITSAPAFGEIKRKISSDSPGHIASKFTEEELMKGFIYYK
QSKDTDSLTDVFKFTLLDQNQPPNESGEYVTMIHIHPKKNFPPRKVQNTEGSIVTKESDVVFIDKYHLGFEDPDSKDEE
IIYSITAQPYFVDSQETLDAGRIINTKNLSMVMKNPLIPGISSFTQAEINYMKIAYMPPMKDIGTDPRLVRFHYTVSDLSG
NVLTGLTFQITLMPVNDQIPQLMINPLSVKEGESVALTTNEIFVMDVDTKMENLVLTFDAMPSRGLLKKDGIVMAIGDQF
SVNDLKRRKIMYYHDGSDDSAEDSFGVTVSDGMFRLSKAIPIHIQLIDDQLPYLTSESSSQLIVTEGGETFITPAVLSAK
DQDTPASKITYIIVQPPTKGSILKGRQNINRFTQQEIDENQIKYMHRSGEIGYNSETDAFKFIIADKILTSMQDVSIYQLNV
TILPVNDRAPKIVLGNPLIVKEGGTSPITFDILGAQDVDTKSEELVFLIEKLPEWGFVENTRPNPGSEKSNAGISIQKFTIK
DLQEGKINYVQANHNKVEPLTDQMVIKVTDGEFFSRETLLTINIIPQNDETPVIEMTDIVVEEGHEMSIDNKILTISDLDEP
ADRLMVSVVQEPDYGKIKMMIERKQLGGEEMVEIPMHDFAARDMQKDMRLYYKHDGSETSDDQFTIEVSDGAHTVR
KTAKVHVKRMNDETPQVIKNAGLRILYGEAAFVSSAILKAWDIDNGPEEVNYILARKPKRGVLQKRIHSPLDTSQDGILD
PDVDDVNGEWEEISEGMNFTQLQINKNLIRYTHTGDMGLMQADNFRFIVTDGKNFSPEESFEIEIEHSQVSEIALLAKG
MKVKEGQRVVIATDILSAEDGTDLFNRIKFNILHPPLFGQIESLNAPGKALKEFSQLDLAARRIAYVHTSRSESIEDWFKF
KVSNGYQAKNGTLFIHIIPTDRITPTLLKNNVLFIKRGTERVISSTYLLVTDPDTKSTNLTFFIMKAPDHGELLNKGIKIIQK
FTQDDVDNERITYKNFGSQAEHDYFWFAMTDNKHIGFLVNGSLHLDPVKFTVALISLDPTPPIILRNKQPTMLDSFERK
KVGYIINSRFLKVTSSHEPAANLQFIITAKPKFGHLEDVARSKVIKKRFTQTDIDNGRVAYIIDSKNKSVTYDSFMFKVIDS
KRNELAPQRFGMSWPVIEFTRLEYIVCEDVGTLSVGLRRVGETSQSSYISVSVREMSAKAGEDFGSQTAAQIQFDPG
VKIVSWDIAIRDDGLPEGKEMFKLLLRSPVNALIGEYDRATIRIINRNNAKGRCIEVPGMIHKGGIVPENTNIKPSSILKTD
KETVYTFHGISGLKTDKKSSSRDENDDDEDAGDLALDAAGNMKPLNDGTGAGASSSGSGSTAPRSMKANRKRGRK
RKDKKKKNMTRKQKKIARFWERRDRMKKKQQKKLREKGERVALRKKKNGDGDDDDSEEVKAPKGKGSRRRRKEK
SESSRRQETPDASTSWMYNGGKGASSLSTDTDKDSGISSGRSSSYDRSSSRGSKSDRGSTSDQSAVTKTDSGRGL
TDDKMKEVFNSMDGRVMTTGEVDSGEGQKKKGKKNKRRRKKKDRKKKKGDEGTVDEKRVKGEVEGDLTESYGPA
DDNEGEDLDMTKISQSNKKVKGKRDKIKKEKLREKRRKIKEKKKRQRMKAQKQNERLKKQKEKERKQKQKNKKHVN
KKGDKKKNNKGNKRKNVNDNVGKDEDVNDNDNVDVNASNKGDVSDEVEVDTIDIDENVENNDKNGKRKRTKLDKK
EKNRKKKDKKKKEKNRVRHRNKNKGKRRGKKKQDVLDATPKGQLGVTLQPTGVTLQQPSVTQHTGVTLSTEQTRG
EKELDKNSKLDVGPTSLVLMPANPSEVPVMTQEVATSPSTYGQNTASADYGPTVASLDLNLENETDSIAGIPITDDRM
SVDMPAVDQQRPRPLSDDSTVKKPGSHGYKNSKLSADDLEKGLLSSRKPSESDLAEGSGAAGEIVYVNDCTPATKG
MLHLDPATSLIYKCDGYDWLSLEDEFRAAVKSTAPSTPTESPTVFKTVVSASECLPGWLKSNGRCYKPFDEQMTWS

MAQKQCRQFSADLPSIWAMQQLDWLWRLIGQDRFWIGLNCMQTFKKWEYSNGDPVTFTYWRTDYPRGGPLTGKN
CVLVREDKFWTNRPCSTLRAKFVCSKLPLSIHRTRN*
>C-lectin13 Partial protein sequence
FQKILKDLCNGNWDQGQGVSDWNTLLLEYSRVRTQAEAGKGFIAKFTSINPIKEQVFNYTEGHCEDGWLRFMSGCYK
YVKDDNNVSLTWTQAEEKCNSLAADSHLTSIQDEYEGDFILQSLFFQWLYDPNLKEIYIGLTDQQHDGRFIWTDGSPL
SYTSWDLGQPDGSVLEGCGMLRIDSLNRRRYWNDIPCAMKMTSQYICKTPAEAVTQYSITDLGGVIGAWDQSCDLN
RQFKCDNGNCIHNFLVCNTRNDCGDGSDEMDCQPINDVCPLDRFRCQNGDCILISKYCDHVLDCPGGEDEAGCV
>C-lectin14 Partial protein sequence
MDVSILKISFFLGLLAIDLHASTSPRMCPDNWIEHDPYCYYASTSQAEMREAAKSCNDSKATLLSINSQDENDWINNTL
LTDSGTSYWTGLMRITITKEHISNETFAWADGSLPDNPEAFIQWAPNEPNKGTDELCGEITGQGTFNDNHCGNDIKFIC
KKAKGCFTPSGGENCSKCHCLDQTCDPHYGNCSKGCDAQHLGFNCSTVLKIRKPEIKSVTDDGAVVRWRVWTGTTI
NSALGPGKIIYQYTVKRVDGDEHESKTKDHPEKKKKLTMKKTFRGLSPATNYSLCVKVSHDQLENNCSEQELVHFTTL
CPAPPPPVDVQVTVLARDSSTDLGFVKVKLAWKVTESPLDGCPQIVSYWLFSRETPTTNWAYHSQVNSSSNSSLHQ
GELTLRINGSYDFMVQAVDTNGNLSESNVTTANNIEPKECFLRYSDREKSGKNCTLCQCLNGVCNQYYGNCSEGCK
DSVIGFNCSTKFRYPTKPIISQVSSTLVTIRWKRWTENINSDKGPVEIIYGIKIKHDGYVFKMENVTFPSTANDDLEHTVQ
GLESSSDYVASIDLLRDDGAVHGRSQSYAKFKTLCNAPPRPMDIDLKSEVGSSIVNLTIRYRAERSGLCDNITRYVVRY
HKVNTQDQHKERRKSGADIQQTLKLEANHTYAIMLEVFNKDNVSSVSPTYWINFQGGNQSSELKKISHSATTAKLNWL
PRRTMLTKENYSVSCHAEGPSVGQKDRAVHPESEPE
>C-lectin15 Partial protein sequence
HYYGTGRTISWTITPPTGQFVRIYNVTLNSTLKNRVEEMKIGDWRHATNVLCQDVYTLYNNGQPIISDFGYYIHGDSQP
VMLKLHACFQSSYHPAGWFQAHVDFVDCPGFGFNQKTCKDTVSVDLNATQHTCGYIGSPLFPNVYPIAGTCNWNIAV
RWTSFIKLEFLNLDVAGDRVSCTGDQVVVTMLGLGVDAKNGPNRVVCNGNRAFTTMDSDWNKISVRYQRVRPEDG
AGKGFIAKFAEVNIEQRLINETYGVCPTGWLQHLEACYLFSKDNELITWTLAESKCDNMNAHLASVQDHDQHEFIKYTL
LTKWDWDWQNPGVYIGLTDQKSDGRYTWTDSSPVSYTNWGPGQPDGSSLEGCSIMEINTLSSGRYWNDIPCAIRKT
RQFICKKSAEGAQPKKAIDSKDIIKSWNGKCNPERQFTCSNGDCIQNMFVCDGYKNCRDGSDEVDCNQTPSQGGCG
INQFQCGSGSCISASGYCNFIDDCNDKTDESSCVWPECGVNEFRCLNKQCISNMSRVKNFVEDCIDGSDEMDKGCN
GFECYNGKCIPYKLHCDGIKDCAGTYSEDETDCTDIATKPLGEGYLNCITNSPYLATQHCIYDKDFLGYTTGCRDFSHL
ENCESFNCPPHYLKCSGSYCIPLRMICDDTADCASAEDELNCDSYTCPGMFRCEGNGNCIIQEKVCDGKKDCVGGD
DELSCSFCPDSCTCVGHAYRCSEKYLQAFPNLPQDA
>C-lectin16 Partial protein sequence
LIRLDTTNKQCLRLNKGASQWKWKDGTCDTGFNFVCKQAPEKVGSCPQGLRALQGFRYLYLNDPGRKYGVARSRC
RLMGGNVISMHSNAEITLAQEIACGAVLGTRVWIGINSGESMAAGWDDGTDFTFLTEQLQAKGFSSGRVVGVNQCAF
MKKGSNGWNLDKSTCEDRISYICKIDEDECIEASTLCPKPNRGVCRNIPGSYACVCDDGYYVEDDSNNCVDKNECAD
KPNKCSDLAYGVCRNTAGSYTCDCVSGYEKLGDACTDTDECTNPDICTANSSCQNYGGFHTCDCNNGFEKIGDVCK
DKDECTAPSNVCASLAN
>C-lectin17 Partial protein sequence
MVGKDSQGSVLGASCPTGWRQQARDQPRFLYVAESASFKDASKICVENGGHLASIHSEVENTFLTNWIEECEGTDF
DGVWIGAQDPNNNKFFGGLYWLDDSINDYTPDGSKIQLDSLPDECIRIKKKQWADTSCDDSNKKSFICKKAPLVGNSC
PTGLHTIGAYRYQYIQGDYNKQDFDGARRYCNLMGGDLASILSFSEEKAIFGMFCNFTKDFLMWLGYDGRTDSTKWT
DGQLLHSNYSSQFASGHADKCLVMELQTSLTWKFEDKNCGEKHSFLCKFSVTDPCSCENGECVKTDGSPSKCVCKV
GYMWSETLKNCVVSNCASEWKQGSQDGPKFLYVSTPKPLHNASQICAYYGGLLASIHSETEFKFITTMIETCSTASTD
VWIGAKDNEDSVNQLKWLDDSPFDYPPDSEKANLIRLDTT
>C-lectin18 Partial protein sequence
MAFVSSSGAFFITLISHLTRPVSGQGYQAKYYALKDGGCNSTHTAGSDAQNISSLNYPMEYFSNLRCVSIIIAEPGYRIL
LRILGLDVPGDASCENDKLVIKVFEEDPGTQLCGSMGNFSRDDLEFQSVSNLLSVAFISDYMLEGRGFYAQYTAVKSC
SNMTVSQKTGDFHSINWPNGYLNSENCQTTIVTNIAERVIQVKFSYLVLESATGGPELDYHCTKDYIEISDGVDVHRRC
GNWSGRESQLVFNSFTNKMVVRFSTNHNISRPGFKAEWSAVNNESAVLHVDCDSDWEESKRHCYKVFNEPRNWTE
AATDCDRRLAYLAKVDNEETMVVITRLVLQNKLHKDISSYWIGANDRQFENDFFWQDGTPVISPDWFPGWSDYRNFS
SQPNDDGMAEQDCVELRQSYPYPSKGIGVAQRLFWNDMSCSMKNSYICMRTKSGVHVPPLKTQKINCSSTLILSGQ
NNTGLLETPNFPKTYNNNDDCVIDISTLLGYRITLTFSHFQLEEDSSCRYDFLELTSPHSAPRTENTVRRCGEWEDKVK
LLHYASTNNHLQLTFHSDHSRVHKGFQVKYSVIAAPHRCDELGAVPWSRSCYYFSANDTADLANTSTKCRDVKSQPL
VIESQEELNFVTSYVNTHYAESQFSLGESLDFNATQMKSTAVVAVTLAASPSQTGTCYVFENSTILSWKGNECDIPLNF
ICKTPFPEHSTATLTITDATSGSLASADYHPSSQNYTNDEKHTVIITVPATHRIIFDIAFVDLEYQQQCLYDFLNFMDLHT
NTSKRFCGYNQRDLRYASDKSSVNVTFFTDSSRTGRGFSLTWKSVDISECNMAQVTGEFGAVSSLNSPFFYLDNLD
CKTWIVAATNNRLWLSFTKFELDYTDNICSDYLEITLSESTLSTTMQETSLKLCGDRNGRVNELRFVSFYNKILLQFHS
DNNARSIGYQAEYKSIPDGLYSHSDMITVNPGETGYVVSPNYGTDYPVDLDYTVRIKTKLQYQITLQPVDIDLRLSGDC
YYDSVTINDDIKTWQGEEGKMAVLCRNPTEATKWISHMNVISFRLQTKTRTNFGFKGTKLSYRIRQDEFFWNKTYRRL
NVTLEECSETTCKNNGTCRNTTSSGYQCDCKGIHTGLFCHITWCELRPSPCKEGKCEANGTYVCKCDQGYHGHDCD
MTLAQCNADKCNNNGWCKPISRYGDYQCQCHTGWDGKVCDESADGTKSLGEILIEEPFWIGIIIILVALFLTLFVWCFR
KKYAHNFHCFRRKKDKVLANHIHDISRRPSPGDLLVYNNPVFSREGSPRYTRSNSRSNSLSVDYKDTWGVPSRKGPE
QNAAAAARFFASLNDITREKSLMDTAGANYSTDSDANYMGIQGGQILLTPANLTINQVQGGSVTPGPSPFHAPLAARS
TDPGLKRVKSAREGGRLRRGHMMKASHKAHSFDYGIHHSDTEVVVDDMFNQFKIPNQRKVEATIEPHQNSLDFDLD

PEARSSARRRRAFATTPQITIQFSPGPGPGPDGNNEERFDFEDEFGHQDGIGQDAQGDHHEHPRRNEKRLSRQGSSGD
STSIRLSPHGSERSSRSGSVDVCGGSRRGSSLEPTPRESRRHSLEPSREGSGHSTRRHSLLPSRDHSARGSKRNSL
EPSRSTSRRGSAGHSPSSKSSRRSSIAVSRNVSQRSSRRYKKKEKRRNSDQPSPTGDQYSRRQSRDSYHGRSRDT
DKREHQSGSKQLQNRYLRDVLRHHLKNFAHRDLSFDSTTGSDVGSVRSLGMAYLTESLDLPCDCDKCIASSSKHRH
RQGHRRRNLEYYSPKDSFDSMPASCDVHSSDSLPSVHLGVRRTASAEPPGMRSSMRSSRSARGYDEEHVKQKSKK
HGSGHHRHRHRSDSKESKKVRHERWATPANRGYSSDKDVVHLIPSIVVDKHRPVMQKSHSEDDYLNSRSQSPVNK
MPAQVFLRVPEMFGEHDHSPSNESSSGIASSPSHSTAGLALSIPEGLDYNPVDYERRLFCDVMKTGARDRTDYEIIEL
ENMVPVNSNNQNSRPGELRKDSDKTDPSESDRSSSSSGLGLMQGKRRTSLGSTSAIEVSDPDDLLLRNSGVLPYKS
RSMPIVIRESHSMRRKHPFRDSAYQTKEHSMSKPPSNQVSPTLPNLIKIRPEPGISTKAQYTRPEDTAVTSQLFAAAKK
LEHLQSDLNSRNNFESETPTTSTQSGVIYHLPVSRKRSYGNGKPINISDIPESSQEIEDETQGATRTTTDDSSHRGSYY
SQESIYDRINGKMAPYNTFDTAGDSAESRDDEVFRTFKSGQNARSVLSVLERQRLSAEAE
>C-lectin19 Partial protein sequence
ETPCGSFQNEGKGRKGKCGGSKDLMDDSNGTSTSESQEVTETPCGSFQNEGKGRKGKCGGSKDLMDDSNGTSTS
ESQEVTEKPSKSPCECNSTDHVHTTTHCKKLVTRDPVKLRRCQGVRRGRYCYRIEHGAACWGEAKTMCEKKDAVLA
IINKKPVYRAIKKLIKRKANSNETFFWVGMSYATSLGLKMWVNNKQVDQSGYWNVHHILKEKEGPVYEGCFGIDSDS
WNLTTMDCATKFPFVCFYDITMNNQTEFNDAPAPKPTGAFGVVTEQNIKTGAIKTHHQIDIILTPEQHEIVKTGGDPSTS
TGAFRFKTATPTQNELQKLFILQMQTRRISRRRKRSTVLSKLWPNGVVNYIFHELTTSDEKSLIKAAMSVMEKRTSIRF
HEGLDSSKIGIMFNTHEARCASEVGRQRRLQYVFISRDSCLHTQGSILHELFHVLGFWHEHSRSDRDDYITINWENIDK
ANKVNFLIQSYSRYASFGYDYESIMHYPQNAMAINPNFDTIKPNPTKWQYQQLGQRFAPTEMDYKELNSLYSSSHPS
NGNWGIWSGWSQCPVTCGGGRQYRIRLCNNPAPANGGNGCPGEASQSQMCMGDSCPDPEIMYSWLGCWKDQG
FPEVLTLIEGLSSYLDEDYRIRANRIRKCALAAKERGAVIFALRFYGQCYVQSVDATEPAEDYKSDGPSSQCGREGHG
GMNEIDVYTFGNEPVQGQWSVWSDYTKCAKSCDVGVRYRQRECTNPPPSKSDDSKLACSGESTEKGECNTGPCPL
HGGWSDWSEWSKCPEEQTSQQTTPKEAAANETQIAYRYRSRRCSNPSPAHGGAQCLGELNEVEMCPTKNVQVPP
CEKTREPRICLIVLPNNVIASSYNNLDCSDDNFEGHNSNVWNKPHQMELVVY*
>C-lectin20 Partial protein sequence
VSCLGCAMSRLSLLLALVACFGAFQLADAKRKLYRKARSLGVEDCEDEGGFVLNDICYILGSDLGTWEEARQSCKDE
YDGDLAVVVKKVVYDYILDIMDTELDEGSVVWIGLSDHDTEGLFEWVDGFVDYEEKGDYWDIFWEWSYDSLGEYPN
KNCVIIDRETGDAKEEGCNERHFYVCEIVTYPDEWELEQMNEEIEWENDLVEMLSNASVMQQMEIGNKESLNDLYEF
DIKLNKEEAAIFESGGDPSSATGAFNFQGDQPTPEELMAIKELIEEYGIKSWDPATREYESRRRRKRSALGSDVKAWP
GGVVPYEFNPYYPAEAKGYVYQAMKVIEQDSCIKFKVRTTETKYMTIGNSDPSSCYTAGIGYPSTWYEQTREMNFGW
CWNKVPSIVHELYHALGFWHEMSRPDRDDYIEIIWKNVQPGENNKFQFQKAEKSTFSGYPYDYLSIMHYHLNAMAID
QNKYTMKPKKLDVKLFGKVGNGQMPTKLDKFELNTVYKCDPKYIQQISAMKGGAVELEEDQSWGPCSATCGIGQKR
KELKLCTKYQCYVQYTMSQCFAPVCPKPGEIIPPASSKGEVAVKQLGYTWGGCWNDDRFPSSLASLEGKSTFLDGKF
ADRSNKIQKCANAAKEQNSEVFALKFGGKCLAVPSTAKDKNVYKANGPSNDCNGMGTVSAMDVYLLNSKDVQGEW
GSWGEFDSCSKSCAGGVQTRKRVCNNPKPKGSGAQCKGSESGTAKCSTGSCPVDGGWGSWGPWDTQYCEDFG
FKMRERKCDNPKPQNGGNQCLADTNFLEVDMC*
>C-lectin21 Partial protein sequence
FNRISLSLVWILGLAQNLVQSQDLVERCVKPKYEFGSYCYVHSGSGPGAAWNFAWVDCTTRGGRLMTIADEATQNYI
VHNIIWKEFGKSAEVWLGATEHYSPWKWVNTNGLIGEPFLSLAGCYKIAGGPDLGPEVFTIDQMIPEKCVQKCLAYRF
AIVQGSRCWCKEKVGSYPKVSKVECSTPCSGDTQLRCGGESAIHVYHTHEREGTYFNWAKTSFTDGFNCVAYDKEI
KGWVNATCNSRNGYGYACHDKNFLHVELYDDSKTWFEARAACAKKGPSSDLAKINSSALSEDIAKKAMLRDRSKRF
WIGATNREWNWIPSGDPVIYSKWFDGYPQGEHKSCLMMKGHVGVVFSNPFSWENADCNETKPYICQHYKYKTTPA
PTQSTTTNAETSTFNEVTTTTSSAPTQSRDLSTIVTEITSAATNEKQTITVDASMADTGRQEDKGDPFPMTLIVVVTVILL
IAIALVIVIVYFVKRKRSQTPNGAKPSQDKSVHFSQNPLYKSSDDAADTDENLAVNIYGTQDDLASSLYDNPAHKAPNT
TGPPLSIQQSGPAGGSRYSIESSDLPSHYVGMTMKGPRSPSNDTLRSDDVEFLTDDTPDLYVSADQLEGVGGFSNPI
YDLPSNIR*
>C-lectin22 Partial protein sequence
LNSSFNRRNCHLDFSGSTIMRSLFFLALCAFVLLVRAEDGEQRQCPKGFTPLIMGCYKAEMTPMTFDDAEDKCLSYN
GVNETRFHLLSVQSVENQTHILKRLELGGKPVWVGILKIKGKPYQWAAKDGLSPDNAPWESTGPSKAYCVIIVNGKW
MKTKCKTPNIFICSGKQSQYIRKADIERWEYQDVGCPRMFALQNGHCYRFFYSKKSWGEAEARCQAVRQGYHLMSV
PSTRDNILMKAMFPYQIKKNRVWLGLRKVGGKDQWTDGSKYRKSRRNWDRHSKSKAKNCTALSKTSRGLPKWERV
KCAEPHEFICKGGTKAMVAITLAPTTTTTTTPAPYQGYG
>C-lectin23 Partial protein sequence
PSGINRTSLRFRLAISYWKRISKMRDCAFATLVVIGIIACCLLQCEAHPKGSRGRGPVRAIKRVEDVEALRARRSTSTET
VIELASGGQVTRLRGNSVKPNAGRVEVLSGGKWGVICDDYWDLRDANVTCKELGFPSGAIEATIESAHGTANIPDDHLA
MDNTLCDGPERSIHDCFHTSEHDCTTDESSGVVCKKTDGCPDGWVYNNGLCYILFEEGLDYQGAAETCTTEGARLL
DIESQKENDFISDWITNNYDLVDHSGILIGASRSSESNVWSWPESVLEFTFVKWFPGFESEFVNPPAATENCISLRNKY
FSSRSYSYANVNYFYWTPVNCDEPRAFICKKVASNVGQCYNGKGTDYRGTTFRTDKGSLCQRWSDTPVNVLTNPG
VGLGWHNFCRNPDGDKKPWCWTDHESNTFGFCALEECTSRPISPSTTQSRCPSTEQFDCYKSTFRLCIAVSWKCD
GERDCPNGEDELVCPLRLDEFTKIPDKGLPLAFVNIFYITLLDGCANLCLTRRDFVCRSFSYDRDDEMCYLSEHNTNTP
GLTLEDCAQCDAYETSSQAADACPGFRCDNGKCVAETDVCDGLDDCNDFTDERDCENKEPFEIRLAGGNANYKGSV
EVKYLGEWGTVCDDNWDMKDAKVVCKQLGFRDAKKAVSGAEFGKGVDNIILDEVKCVGDETSIENCPHDPWRKHD

CRSHEVAGVICESNRGCEAGEWQCSNKKCVALGWVCDGTDDCGDGSDENRCDRSEILVSLVGSSNPQEGRVEITR
NGIVGSICDDNWDDKDATVVCKMLGYKSGTATIEAKFGEGRGPIWLDEVDCTGSEDHIVNCPADNWGTTNCDHSED
AGVVCSNLDRTTRLPVPRPTRPVGPIDCGRRPIGQPSARVVGGFDAIYGAYPWQVGIRKNEDGLWVHWCGGTIINDQ
WIITAAHCFEDEPKEIFRVRTGDHDNEVLDASEEEFEIAELISHRQYNTGDFENDYDIALIRIRPNAQGNGITFNNFVQPA
CLPTASTPYTQGMKCFVTGWGDTGTDYPRILQVARVPVIAETECSRMYKSAVSPFMLCAGYRAGGVDTCQGDSGGP
LVCGVGGKYTIMGVTSWGNGCAEPESPGIYTKVGKLVDWIDRKMANT*
>C-lectin24 Partial protein sequence
GDEAVYATPCVMMDGLVDYRWVRVPCYQQHAYICEYEMESCPSKGVTVAEKCITISQTNETWFRARYDCIQNSGDL
VSIATEQVQTDLMTYLTGLNRTFWIGATSFKWKLKNDTLIFTDWGQGQPNPESEQGVYTQLYSKEGSPDLKWQLEHG
VFVGKYICKRAIEGTDQPGTTPALPSPHSSPGSTATVLPHGTTEENDPNALQQDDIPVVKIVIVAVIILVLSVIVAICIIVIKK
RRQRPDLYSDTDKLSTVSDIEDFTNPMNHVNHGYDNVTTTTTPNTVSPTADAKPVKAAKMDGMKYEKQKEKVQTLY
RTASGRPVNRTSTGDTIEPESSGLPAMDEDYHDPNEASLRTNPSNKSRESRSSFANPFYQGWGETDDEHYETPNEP
IMTKVFDFTGSMKSKGPSADGETPGLSVAKERPGTSPSREARKLESVPDRPAQPLPGDEPA
>C-lectin25 Partial protein sequence
TTQAPTSATPKSSIVTTCPATWKKYSGACYQMFTGSDEKNFKNAEDYCRAQKGNLASINDKKEYEYIMKEFMNTLPN
AGGITSLWIGLYCKDGKDFFWTDGSDTEGKAFYKPGLMYFKRHHKLATITLSKMGNMNGYFGVPMTKSLPFVCEITL
GGKKKNSKEAACGYGWRRFKGRCYRLFNEYERFFAAERQCNQYHGHLVSLNSKKEHKFVRGLFKTKHPVFGAWVG
MLHIKHDNARQPTFQWTDGSLTDDYTPPTMFNNLPAPNPNYGVMCKNSSILDVTPYYTLPFICEREMPKGCDPGWD
RFQDFCYLASKTPATFGDGMKTCRKLKATLASVCSFNEFKHMMRIRQLFGFQKVLTGARFLKSDKREILIWSDRRGCI
YNPPGKYANGLDKFGVLTQSGLTREPGTNKFPVICKKMAKNACENGFKKIYQHDDPLLLVTSASDIKECKKACLEKDG
CDGTTFAEYQGNKMCYLHHVDGGQLPMMTSVDLQHWKICRNY*
>C-lectin26 Partial protein sequence
MLQFESSIVLFVLCLSSDWLLIEGACRQGIWQQPGGSTGPMFRLSTSEATFDDAKQQCINLGGQLASIHSAEENSFVI
NMLRGCGKTRAWIGMVDPDNDMNTFHWIDGSDVDYSAEKTADNIKWFGYKIEECVNLRRSIFMSGGWRWNDAECD
KSILYVCRKYDTDECASASGNDCDTDATCTNTHGSYTCQCK
>C-lectin27 Partial protein sequence
FADKLNQADALEVCKSIKGDLISIQNPDEQQFLDTVLVFDYEDHWTLGKKDASGNFTYGGTEIAWTNWLADDLLTNVP
DGTEVCLSVAKNLFTKGFFLMKGKWDTKPCSEKKSFVCQTVAKRCLAMPTSGHVEPHPPKPYIELTIDYKCKPGYVM
PNYTARYLEWQQQEPSNATENMTEEWEASKPNITTMPSMKYECLFPGKWSDGSLAPCEPTRCKVPPDVKHADRNS
SERGFPSSIEYTCQHGWHWEFDNTSAKVAWCGPDSDWIDVPPDCDPVECEPFPRVGNATVNSANNRFKETVVWTC
NRGLMF
>C-lectin28 Partial protein sequence
VTTTEIRMGLRRIHLCILLFHLISKGSPRQNNYCLTDWQKPPGASGILYEYVRQSLSHQKAQESCATKGGNLASIHSKE
VNNWLINWVFRCDSRNGIAWIGLQDPNNTVEDLVWTDQSSFDFKYNDDHIQLNDQHGAE
>C-lectin29 Partial protein sequence
VITFHNETLDRTAASEKCHAEGSTLAKIASQRIQEFVLEKAQQLREVRFIKSKSLWFGYDCTFNAETKVKKFFWADGKD
AEYENCAKKSCWAKYCKSYTKWLFCVKPCSYKTSWLSPGEWVHAYITTKANFICTH
>C-lectin30 Partial protein sequence
KAEKKCELQGGHLLTLATDAQWEMMKDELVKAGQTKPVLIGYTDADMEGQFVWADNTAFYEKWAPNEPSTVDSQK
KDCVVIDPAKGFKFVTVPCNGKYAYACSQDLNPYRPCPPGWKYDYEKAGEDCYFEKPDKESFYDSYMTCIEHGADL
PKITSKEDEDKLMATFTMPEGVKM
>C-lectin31 Partial protein sequence
YRLFELSNPPREVTWNFAEASCLSLPYNKPNLACANTQGETDTVTSLVRRSKVERSWFGLRRNSHMDSWTWADNS
RMNMTNFHRFLNENIDYGNGFMDDSATPSGSWDVSTAKDGVRATAVFYVCQVEKSVN*
>C-lectin32 Partial protein sequence
KRFKDSCYRYVGGRFKFEWSEKWCRSHNAHLASVNSKQELNFLLKYAKPTLWTCGPWIGMQTRDTKTFTWFDHSS
TSYKPPTRVGERRGHH
>C-lectin33 Partial protein sequence
HYELFFVFKVILEGMGTRTGNIWIGLTDKTIPGQYVWSDNWPVTVTNWGPGEPTRGKDEGCVAMDKVSGVWTDIKC
AEALPVVCKISTDTPPTLPSVQIGQCDEGWIPHESKCYMYKLSDDKSKWAEGNFK
>C-lectin34 Partial protein sequence
MLRLALFLSIVLYANAFLANTNDECTQEAENYHAESNSCYMIPLGELQTWNGAQKFCQGRGGDLASIPSQSAQDFLE
VLIEKLNPVAKDNALFININSIGSQSEGWMDGETSTYRHWIGGRDKGPWSMSRP
>C-lectin35 Partial protein sequence
ELYRLELVGANSSTISGTCKLSFKYKDNTGEVHCIHRPTYFEILSKIPSAAGIISVPFNLIVKETTSIEDICVTTKMTTISAIS
NTLLNINGSSRHENITIKTVGSEPIIELYSRGKFPYIIEVVQKHYRMPQGDEECICPEGFLLSGCTCYGMFDADTFNVTS
FQEAEDACASLVPGGHLLSLTSSEEVENITHFLRSAWWNFTVGHDWITGLITENKTLMSRWWDEATFSTGNPPTRRM
GERYRLDEGDPLSGNTTHCGVIRVDRHRLLQNSTWFLVHCEHFDWNYSAADSGTVGFGCECSARRIEGLESQYKED
SVHKCHSGYYNCDRGQQVCEIQRGLSEQIIDEMIETNKTVCVGQACPNGWFCSKHVASYIRNTSAIVDLSVLPKRFKA
IDVGLSTGIVTLHEESFRGNSNIVWLRLIKQCNYSHPVGFNISSLSSLPGLLYLELSGYPITGLNLLEQFKQMQHFGLSR
SCRFVFSIPVYPPNLTYLDLSFIRIDVGFKGMLNLSQLRYLNLTNTSFKNYINISQVLVLTRLQQLVLKLCIPFRHANLDFV
TSLTNLRLLDIGSLLLYDI*

>C-lectin36 Partial protein sequence
MYYDDSHHARRKDNNILRNSAARWNNFTDMVLRVQVHLLLIFCSVATLTDASYLSIRLTKDPYLLLAQNQTGPLVPPR
PCSPNMLKGYQDPDNREKYPQCLGCHFNISFDWTVNATGPLICSIDLLNSYYPGCSTNRTYFNVEIRGLDLGKDDCITI
EDVIHKIGSCHSHGGVSSRPKTIKKKFCGPIQNISFRTLTNEPFLEFTQSSSPGNTTLLHDVIRMREIPFRKPLETESPEC
ECPESFTKHRCMCYGAFRGEYNDQTTWMTAEELCKDKGAHLLSIMDDEEMTDIKHFMLSVWMKTVIGWSRNQYEM
PWIYIGLTDQKTINNFRWTDGNPAVFAEWYHPAYAEHLPDLDKAIIKTSPQPTSLPTSRCTVLYISLETFWERTFWSKV
PCSYPIENSAYICKKSAMRHDGRHTDGLNKVDDKPTPSPTFEGGSAVPHNSTIQFPTQPVTPIVTKHVPQPNNSSLLE
FQGMAIVNSGADLTTCEPGWTFYKGTCRIMLTSLLPLNDHPMEDLLSKVAAVVNATIVWNSIVEPGEQDLNELLRMW
RHREEYGGSVLVVKNETQLGMIHYSRAEKNFQPILDYFIKLLNSDITPWSQGHSREARHMIF
>C-lectin37 Partial protein sequence
LRAAVIGVVIFLIILAVPALGYCCTWYQYHLDRSFSYEYAVVSPDTRGVGYTWYEAQDMCLANGGDLLSFATRAEKSW
IKRNHTAGLKLWTGLNRLDKIDKWEWTDKTNDTQQVPDLRFTERNKKCAGFVVGQGTWFIENCGNHQGKGHGLGFL
CKRLVRDSAPKFRPPKKLVSGFCPPCWTDAGTESSKCYGYFDHPHQREMTWKFAEASCLSLPFD*
>C-lectin38 Partial protein sequence
HAGYTRSSTPCQSEDWAYSNQTGHCYLFFLDTVTQDEAIKACKAKHSELLSIENQDEQNFVDTVLRFDFYDQWTLGR
KDEFGNFTYNGTEITYENWLADNLLTKVQEGEEVCIALVKIYYRGFYLLKWKWDTKPCGEKKSFACQHIANRCPPLPL
VDFVEPHPPKPFIEL
>C-lectin39 Partial protein sequence
GTQTNFTPSVRGQPWMEWVWDTTVPQLTLRPWFRMRPSGLVDDKYLALYFVNGRWWWYDEHRTSSLDYVCEKG
MQRALLGSEDEKECPQGWRTFQERCYHFSIHANKARQGAEEMCMEMGGRLVALETEAELEFLTTTMREVKHYDGV
KWWTGGIPIARQWVWSQDYMMFKDNRDQFLGWAPGEPNNWDDQENFITLLYNNTWGLNDVPNRDHISLPFICEKP
RKDNDGVYCPSKIGRTKLTKLPVTGDNWATLML
>Actin Gene sequence
CACCCCATGTCTAATCCAATACATAGTACCACTCTCCGATATTCTGATTTAGCAGTGATTCACCCCCAATGAGCAA
TACCACCAGATTCGTCAGAAAATTAAGAAATGCGGATATCTGGCAGCATGGCTAATCAACATGGCTAACTCATTC
TGATTCGCTACGACAGAATTCCTCGATCAAAGTTACCTTTTAAATCCTCTGTCCATTTCCACCCTGCAGACCTACA
TTTCAAGAATCTATAGCAAGGAAGTTGCGAGGTATCGCTTTGTCACCTCATGCATTGTTAATTGTCAAGGCGAGT
GGACTCGAGCTTGCCATATATGGGCATACAACTCCTTATATGGTAAACGGGACAGCCCGGACACTAGCAGCTGT
GTATGGGCCTACAGTGTGTGTGTGGTTTATAGTGTAGTGAGGAGAATTGCCAGCTCACTTGAAAGCCATCAGCC
GATCGCAGCAGACGCCTTGAGGAATTTTCGCCTAAAACTAAAATCTAACCGACAATATGTGTGACGACGACGTTG
CCGCTTTGGTCGTAGACAATGGATCTGGAATGGTAAAAGCTGGCTTTGCCGGTGATGATGCCCCACGAGCTGTC
TTCCCATCCATCGTTGGACGCCCAAGACATCAGGGTGTCATGGTTGGTATGGGCCAGAAGGACAGCTACGTCG
GCGATGAAGCCCAGAGCAAGAGAGGTATCCTCACTCTGAAATACCCCATTGAGCACGGTATCGTCACAAACTGG
GACGATATGGAGAAGATCTGGCATCACACCTTCTACAACGAGCTCCGTGTTGCCCCAGAGGAGCATCCAGTCCT
GCTGACCGAGGCTCCCCTCAACCCCAAGGCCAACAGGGAAAAGATGACACAGATCATGTTTGAGACCTTCAACA
GCCCCGCCATGTACGTCGCCATCCAGGCTGTCTTGTCCCTGTATGCATCCGGTCGTACTACTGGCATCGTCTTT
GACTCTGGTGATGGTGTGTCTCACTGCGTCCCAATCTACGAGGGATACGCCCTTCCCCATGCCATCCTCCGTCT
GGACTTGGCTGGCCGTGATCTTACAGATTACCTCATGAAGATCCTGACCGAGCGTGGCTATTCTTTCACCACCAC
TGCTGAGAGGGAAATCGTTCGTGACATCAAGGAGAAGCTCTGCTACGTCGCCCTTGACTTCGAACAGGAGATGC
AGACTGCTGCTTCATCCAGCTCATTGGAGAAGAGTTACGAACTTCCCGACGGACAGGTCATCACCATCGGAAAC
GAGAGATTTAGGTGTCCAGAGGCCCTCTACCAGCCCAGCTTCTTGGGAATGGAGTCTGCTGGTATCCACGAGAC
CACATACAACAGCATCATGAAATGTGATGTTGACATCAGGAAAGACTTGTACGCCAACACAGTGTTGTCTGGTGG
TTCCACCATGTATCCAGGCATCGCTGACCGTATGCAGAAGGAGATCACTGCCCTTGCTCCACCAACGATGAAGA
TCAAGATCATTGCTCCACCAGAGAGGAAATACTCCGTATGGATCGGTGGCTCCATCCTTGCCTCTCTCTCCACCT
TCCAACAGATGTGGATCAGCAAGCAGGAATACGACGAGTCCGGCCCATCCATCGTCCACAGGAAGTGCTTCTAA
ACTAACTGTCTTTCACAGCGAACTGACTTTCCGAATTGAAGAAACCGAAAACGAGCCAAGAACTTGCAGTTGATG
GACACTTAAAATGCAAATGGGGTGGGGGTTATCGTGCTAATCTAAAGTATCCAAGCGTCCGTGAAAAAAGCAATT
GGTGTCCACATTGAATTGTTCAATGCTCGTTTCCACATTCCAGATCGTTCCAGCTGATAAAAATGTGCTTGAAGAA
ATTAGAGAAGTTGAGTTGATGGCTTGAGTCCTCTATTCTAAGCATTTTTCTACTTTTTTTGTAACGGCCCGAAAGT
GAAATTACATAATCATGAAACAAACTACTGATTTTGTAAAAGAAATGCCCAAGGCTCTGCGCGATTGTAAAATAAA
TCTATTGTTAATATTTGTAAAATTAAAACTTTTAATATACGTTATCTTTGCTTTGCCTGCATGTGCTTGTCAAACGAA
AACTGAAAAGCACCTTCTTCAAATCACGGCGGAAAATGGTCAAGAACCTTACTGTTGTCGAAAAAGTGTGCAAAC
TGTTCATTACCCACCGATCAGAACTTGAAATTACTAGACCAGTTTGCCCACTTTTACTTCCCATACAAATGTTTGT
CCAGATGCATTTCAGTTTTGGGCGTTGTTGGCTAACCATGTCTACTTTACCTCGTGATATGTCGGCGTCTATAGC
TGTGGTTCCACTTATTCAAAACTCGCCAAAACCCC

**Supplementary Figure 1 – Sequences retrieved from the *Lineus ruber* transcriptome survey.** Next to the protein name, it is indicated if the protein sequence is complete or partial.
