## Supplementary Figure 2 for "The Toll and Imd pathway, the complement system and lectins during immune response of the nemertean *Lineus ruber*"

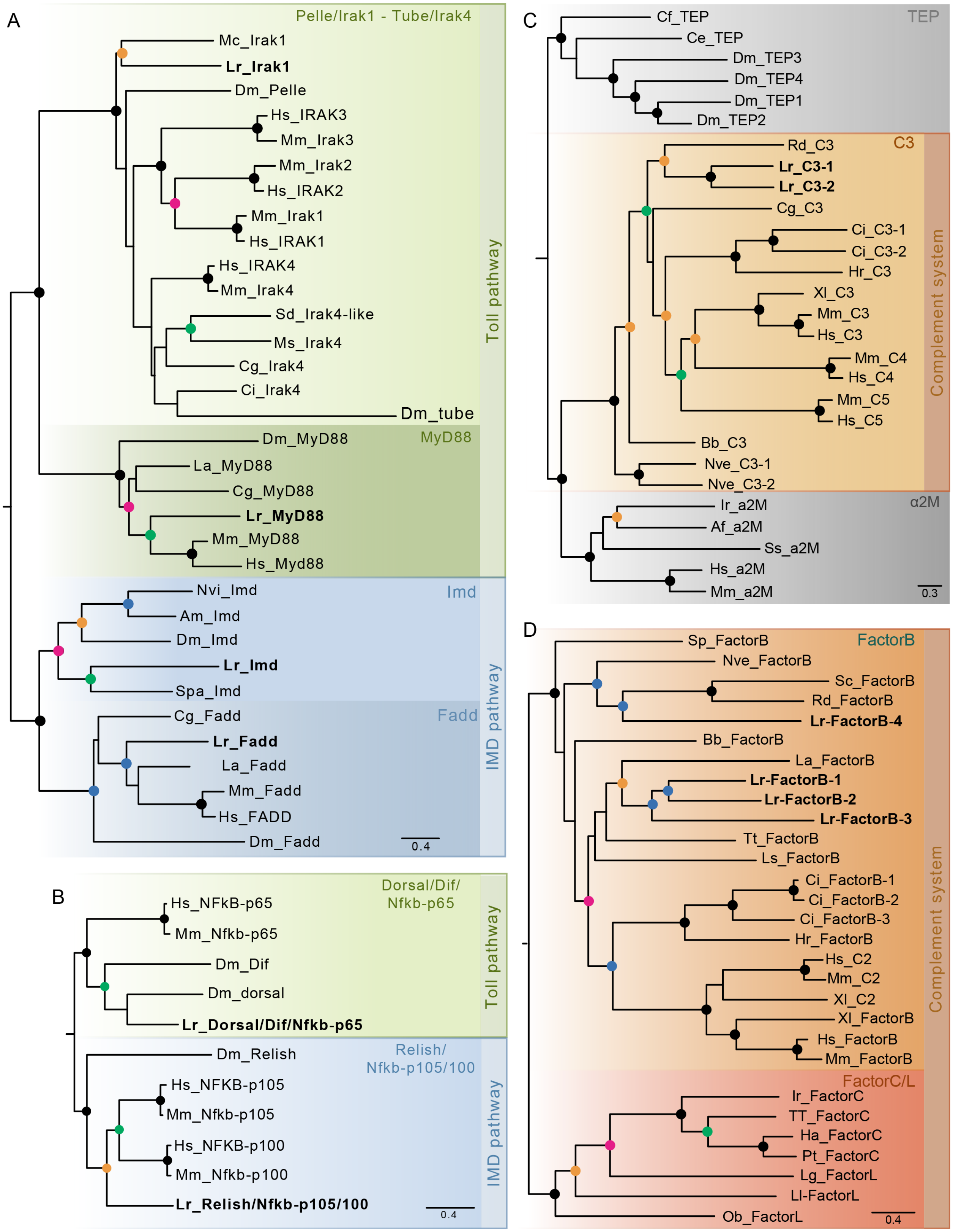
A
Pelle/Irak1 - Tube/Irak4
Mc_Irak1
Lr_Irak1
Dm_Pelle
Hs_IRAK3
Mm_Irak3
Mm_Irak2
Hs_IRAK2
Mm_Irak1
Hs_IRAK1
Hs_IRAK4
Mm_Irak4
Sd_Irak4-like
Ms_Irak4
Cg_Irak4
Ci_Irak4
Dm_tube
Toll pathway
MyD88
Dm_MyD88
La_MyD88
Cg_MyD88
Lr_MyD88
Mm_MyD88
Hs_Myd88
Imd
Nvi_Imd
Am_Imd
Dm_Imd
Lr_Imd
Spa_Imd
Fadd
Cg_Fadd
Lr_Fadd
La_Fadd
Mm_Fadd
Hs_FADD
Dm_Fadd
IMD pathway
0.4
B
Dorsal/Dif/
Nfkb-p65
Hs_NFkB-p65
Mm_Nfkb-p65
Dm_Dif
Dm_dorsal
Lr_Dorsal/Dif/Nfkb-p65
Toll pathway
Relish/
Nfkb-p105/100
Dm_Relish
Hs_NFKB-p105
Mm_Nfkb-p105
Hs_NFKB-p100
Mm_Nfkb-p100
Lr_Relish/Nfkb-p105/100
IMD pathway
0.4
C
TEP
Cf_TEP
Ce_TEP
Dm_TEP3
Dm_TEP4
Dm_TEP1
Dm_TEP2
C3
Rd_C3
Lr_C3-1
Lr_C3-2
Cg_C3
Ci_C3-1
Ci_C3-2
Hr_C3
Xl_C3
Mm_C3
Hs_C3
Mm_C4
Hs_C4
Mm_C5
Hs_C5
Bb_C3
Nve_C3-1
Nve_C3-2
Complement system
α2M
Ir_a2M
Af_a2M
Ss_a2M
Hs_a2M
Mm_a2M
0.3
D
FactorB
Sp_FactorB
Nve_FactorB
Sc_FactorB
Rd_FactorB
Lr-FactorB-4
Bb_FactorB
La_FactorB
Lr-FactorB-1
Lr-FactorB-2
Lr-FactorB-3
Tt_FactorB
Ls_FactorB
Ci_FactorB-1
Ci_FactorB-2
Ci_FactorB-3
Hr_FactorB
Hs_C2
Mm_C2
Xl_C2
Xl_FactorB
Hs_FactorB
Mm_FactorB
Complement system
FactorC/L
Ir_FactorC
TT_FactorC
Ha_FactorC
Pt_FactorC
Lg_FactorL
Ll-FactorL
Ob_FactorL
0.4

**Supplementary figure 2 – Phylogenetic analyses of the putative components of the Toll-, the Imd- pathways and the complement system. A.** Maximum likelihood phylogenetic analysis of DEATH domain containing proteins of the Toll and the Imd pathways. **B.** Maximum-likelihood phylogenetic analysis of Nfκb factors in *Lineus ruber*, *Homo sapiens*, *Mus musculus* and *Drosophila melanogaster.* **C.** Maximum-likelihood phylogenetic analysis of proteins belonging to the TEP family. TEP family is constituted by TEP, C3, and α2M proteins. **D.** Maximum-likelihood phylogenetic analysis of Factor B, C2, Factor C and Factor L proteins. For all trees, dots indicate support values ≥ 60 (black dots: 98-100%; blue dots: 90-97%; green dots: 80-89%; orange dots: 70-79%; pink dots: 60-69%). Tip labels indicate the species name abbreviation followed by the gene name. *Lineus ruber* proteins are labeled in bold. Species abbreviation: Af: *Azumapecten farreri;* Am: *Apis mellifera;* Bb: *Branchiostoma belcheri;* Ce: Caenorhabditis elegans; Cf: *Chlamis farreri*; Cg: *Crassostrea gigas;* Ci: *Ciona intestinalis*; Dm: *Drosophila melanogaster;* Ha: *Hasarius adansoni;* Hr: *Halocynthia roretzi;* Hs: *Homo sapiens;* Ir: *Ixodes ricinus;* La: *Lingula anatina;* Lg: *Lottia gigantea; Ll: Littorina littorea;* Lr: *Lineus ruber;* Ls: *Lepidonotus squamatus;* Mc: *Mytilus coruscus;* Mm: *Mus musculus;* Ms: *Melanaphis sacchari*; Nve: *Nematostella vectensis;* Nvi: Nasonia vitripennis; Ob: *Octopus bimaculoides;* Pt: *Parasteatoda tepidariorum;* Rd: *Ruditapes decussatus;* Sc: *Sinonovacula constricta;* Sd: *Suberites domuncula*; Spa: *Scylla paramosain;* Spu: *Strongylocentrotus purpuratus;* Ss: *Scylla serrata;* Tt: *Tachypleus tridentatus;* Xl: *Xenopus laevis*. Accession numbers: Dm_Imd: Q7K4Z4; Am_Imd: XP_016767530.2; Spa_Imd: AZK36044.1; Nvi_Imd: NP_001135910.1; Hs_MyD88: NP_001166037.2; Mm_MyD88: ID17874; Dm_MyD88: ID35956; Cg_MyD88: NP_001292287.1; La_MyD88: XP_013416180.1; Dm_Fadd: NP_651006.1; Hs_Fadd: NP_003815.1; Mm_Fadd: NP_034305.1; Cg_Fadd: NP_001295786.1; La_Fadd: XP_013392457.1; Dm_Tube: NP_001189164.1; Mm_Irak4: NP_084202.2; Hs_Irak4: NP_001107654.1; Cg_Irak4: XP_011428693.2; Ms_Irak4: XP_025204662.1; Dm_Pelle: NP_476971.1; Hs_Irak1: NP_001020413.1; Mm_Irak1: NP_001171444.1; Ci_Irak4: XP_009859368.1; Mm_Irak3: NP_001346113.1; Hs_Irak3: NP_001135995.1; Sd_Irak4-like: CAL36106.1; Hs_Irak2: NP_001561.3; Mm_Irak2: NP_001107025.1; Mc_Irak1: QCO95270.1; Dm_dorsal: P15330; Dm_Diff: P98149; Hs_NFkB-p65: Q04206; Mm_NFkB-p65: Q04207; Dm_Relish: Q94527; Hs_NFKB-p105: P19838; Mm_NFkB-p105: P25799; Hs_NFKB-p100: Q00653; Mm_Nfkb-p100: Q9WTK5; Dm_TEP1: CAB87807; Dm_TEP2: CAB87808; Dm_TEP3: CAB87809; Dm_TEP4: CAB87810; Cf_TEP: EF210036; Ce_TEP: CAB05007; Rd_C3: FJ392025; Cg_C3: NP_001292308.1 ; Nve_C3-1: AB450038; Nve_C3-2: AB450040; Ci_C3-1: Q8WPD8; Ci_C3-2: Q8WPD7; Hr_C3: AB006964; Bb_C3: AB050668; Xl_C3: AAB60608; Mm_C3: P01027; Hs_C3: P01024; Mm_C4: P01029; Hs_C4: AAB59537; Mm_C5: P06684; Hs_C5: P01031; Ir_a2M: EU835901; Hs_a2M: P01023; Mm_a2M: Q61838; Af_a2M: AAR39412; Ss_a2M: ABD61456; Hs_FactorB: CAA51389.1; Mm_FactorB: NP_001136178.1; Xl_FactorB: NP_001081234; Bb_FactorB: XP_019626103.1; Ci_FactorB-1: NP_001027973.1; Ci_FactorB-2: NP_001029011.1; Ci_FactorB-3: NP_001027974.1; Hr_FactorB: AAK00631; Spu_FactorB: NP_999700.1; La_FactorB: XP_013415956.1; Nve_FactorB: BAH22726.1; Sc_FactorB: QEX93860.1; Rd_FactorB: ACQ91095; Ls_FactorB: MG596914.1; Tt_FactorB: BAM15263; Hs_C2: NP_000054.2; Mm_C2: NP_038512.2; Xl_C2: NP_001116166.2; Ir_FactorC: AII02148.1; Tt_FactorC: P28175.1; Ha_FactorC: BAR45633.1; Pt_FactorC: XP_015930211.1; Lg_FactorL: XP_009058759; Ob_FactorL: XP_014774764; Ll_FactorL: MG596893.1.
