## Supplementary Figure 3 for "The Toll and Imd pathway, the complement system and lectins during immune response of the nemertean *Lineus ruber*"

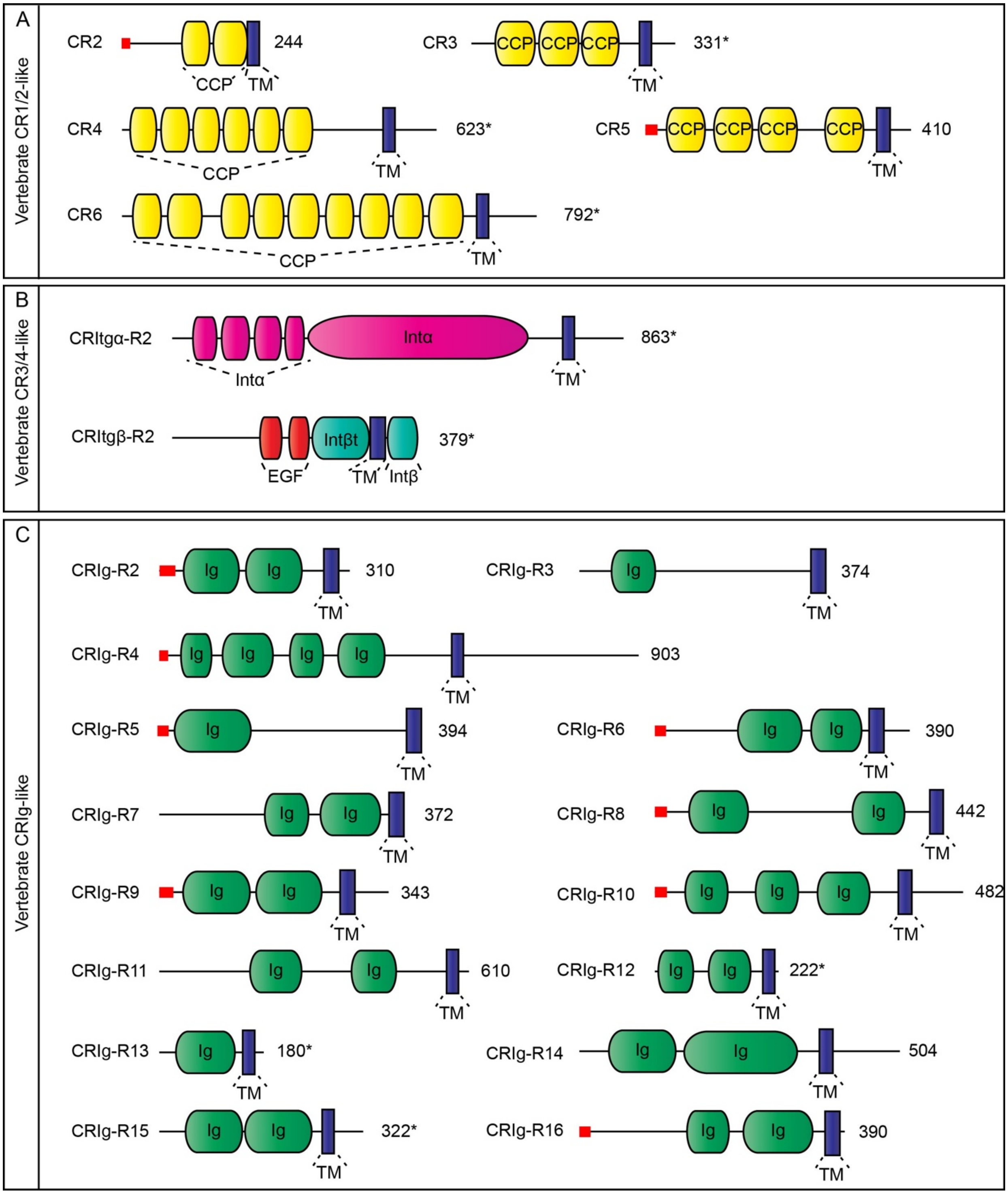


**Supplementary figure 3. Complement receptors in *Lineus ruber.* A.** Domain architecture analyses of *Lineus ruber* proteins with similar domain architecture than vertebrate CR1 and CR2. **B.** Domain architecture analyses of *Lineus ruber* proteins with similar domain architecture than vertebrate CR3 and CR4. **C.** Domain architecture analyses of *Lineus ruber* proteins with similar domain architecture than vertebrate CRIg. Red rectangles indicate signal peptides. Numbers adjacent to each protein indicate the length of the protein in amino acids and asterisks indicate partial proteins.
