## Supplementary Figure 4 for "The Toll and Imd pathway, the complement system and lectins during immune response of the nemertean *Lineus ruber*"

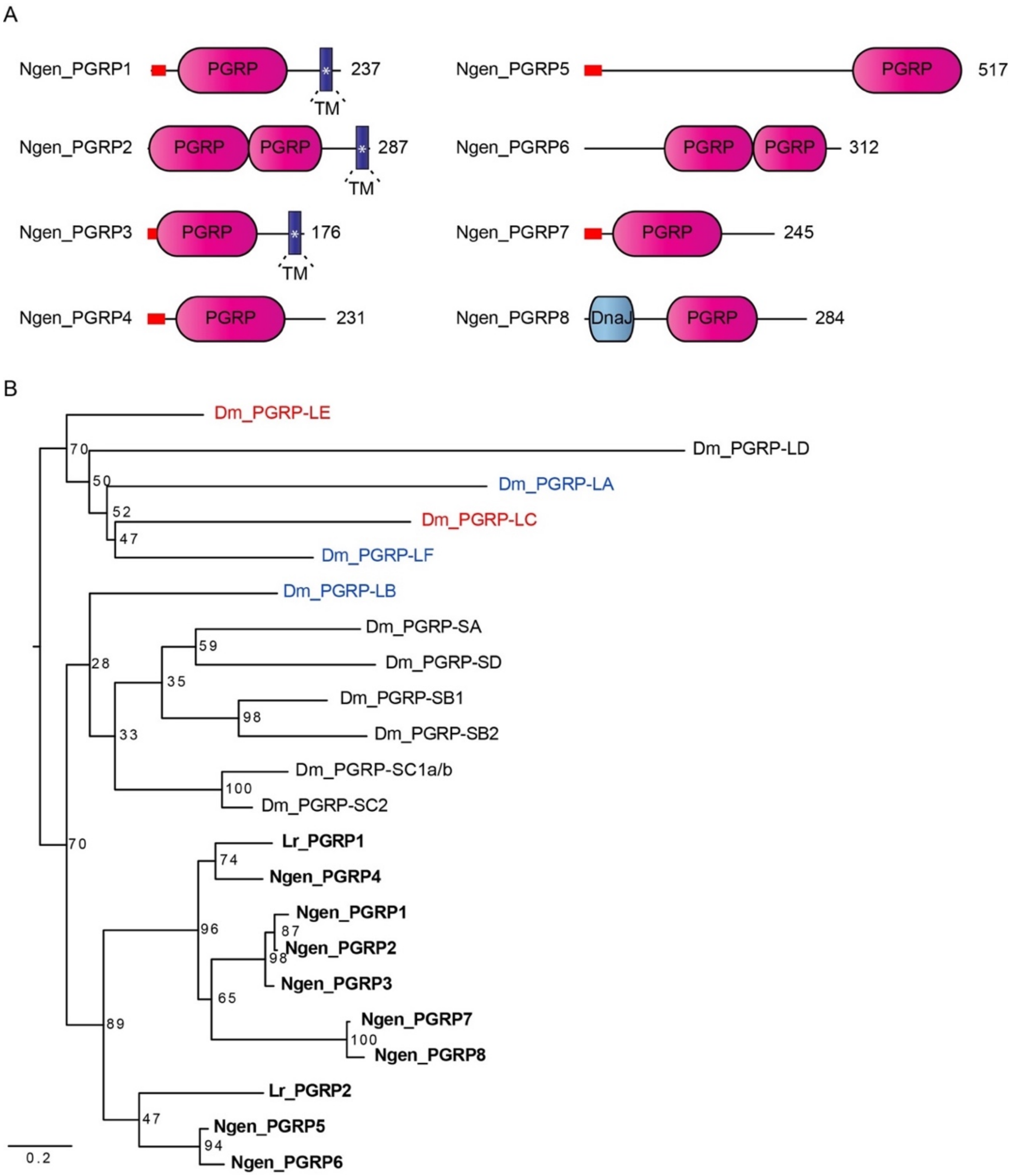


**Supplementary figure 2. PGRP proteins in *Notospermus geniculatus* and *Lineus ruber.* A.** Domain architecture analyses of PGRPs in *Notospermus geniculatus.* All the PGRPs in *Notospermus geniculatus* contain only one PGRP domain, with the exception of Ngen_PGRP2 and Ngen_PGRP6. White asterisk indicates that transmembrane domains for Ngen_PGRP1-3 were only detected by hmmer online software and not SMART online software. Red rectangles indicate signal peptides. Numbers adjacent to each protein indicate the length of the protein in aminoacids. **B.** Maximum-likelihood phylogenetic analysis of PGRP proteins in *Lineus ruber (Lr)*, *Notospermus geniculatus (Ngen)* and *Drosophila melanogaster (Dm).* Nemertean PGRP group forming an independent clade than *Drosophila melanogaster* PGRPs. *Drosophila melanogaster* sequences group forming two clades: a clade formed exclusively by long PGRPs and a clade formed by all short PGRPs and a non-transmembrane long PGRP (Dm_PGRP-LB). The later clade is the sister clade to the nemertean PGRPs. Numbers next to the tree nodes indicate bootstrap values. Red labels indicate *Drosophila* PGRPs involved in Imd pathway activation; Blue labels indicate *Drosophila* PGRPs involved in Imd pathway regulation. *Drosophila melanogaster* accession numbers: Dm_PGRP-LA: Q95T64; Dm_PGRP-LD: Q9GN97; Dm_PGRP-LE: Q9VXN9; Dm_PGRP-LF: Q8SXQ7; Dm_PGRP-SA: Q9VYX7; Dm_PGRP-SB1: Q70PY2; Dm_PGRP-SB2: Q9VV96; Dm_PGRP-SC1a/b: C0HK98; Dm_PGRP-SC2: Q9V4X2; Dm_PGRP-SD: Q9VS97.
