## Supplementary Figure 5 for "The Toll and Imd pathway, the complement system and lectins during immune response of the nemertean *Lineus ruber*"

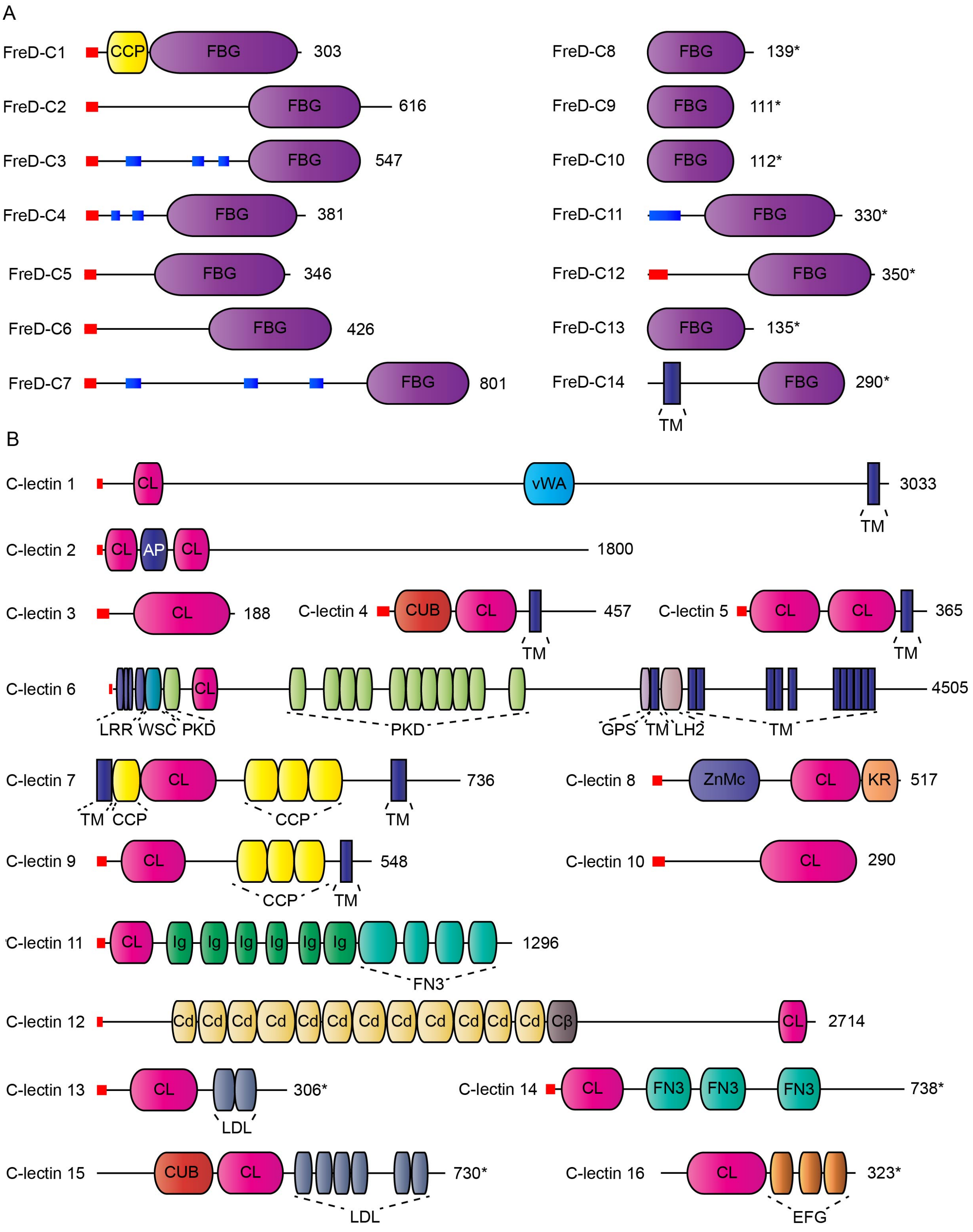

A
FreD-C1
CCP
FBG
303
FreD-C2
FBG
616
FreD-C3
FBG
547
FreD-C4
FBG
381
FreD-C5
FBG
346
FreD-C6
FBG
426
FreD-C7
FBG
801
FreD-C8
FBG
139*
FreD-C9
FBG
111*
FreD-C10
FBG
112*
FreD-C11
FBG
330*
FreD-C12
FBG
350*
FreD-C13
FBG
135*
FreD-C14
FBG
290*
TM
B
C-lectin 1
CL
vWA
3033
TM
C-lectin 2
CL
AP
CL
1800
C-lectin 3
CL
188
C-lectin 4
CUB
CL
457
TM
C-lectin 5
CL
CL
365
TM
C-lectin 6
CL
4505
LRR
WSC
PKD
PKD
GPS
TM
LH2
TM
C-lectin 7
CL
736
TM
CCP
CCP
TM
C-lectin 8
ZnMc
CL
KR
517
C-lectin 9
CL
548
CCP
TM
C-lectin 10
CL
290
C-lectin 11
CL
Ig
Ig
Ig
Ig
Ig
Ig
1296
FN3
C-lectin 12
Cd
Cd
Cd
Cd
Cd
Cd
Cd
Cd
Cd
Cd
Cd
Cd
Cβ
CL
2714
C-lectin 13
CL
306*
LDL
C-lectin 14
CL
FN3
FN3
FN3
738*
C-lectin 15
CUB
CL
730*
LDL
C-lectin 16
CL
323*
EFG

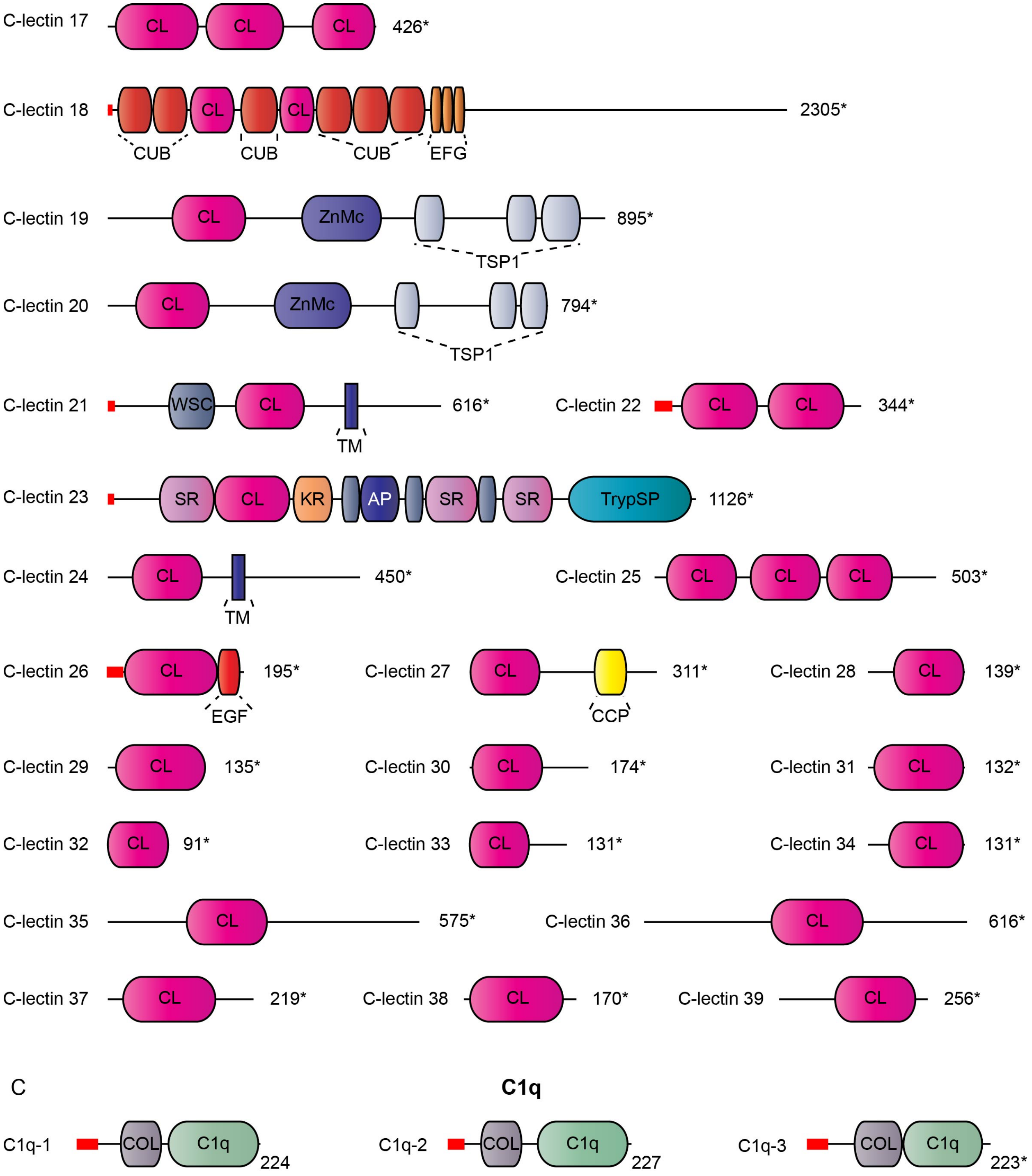


**Supplementary figure 5. Lectins and C1q in *Lineus ruber*. A.** Fibrinogen-related domain containing proteins (FreD-C). **B.** C-type lectins (C-lectins). **C.** C1q proteins. Numbers adjacent to each protein indicate the length of the protein in aminoacids. Asterisks next to the amino acid number indicate partial proteins. Red rectangles indicate signal peptides; blue small rectangles are coiled coils.
