## Supplementary Material and Methods for "The Toll and Imd pathway, the complement system and lectins during immune response of the nemertean *Lineus ruber*"

### Supplementary materials and methods

#### Animals and bacteria

Adult *Lineus ruber* (1) were collected during winter in a rocky beach in Bergen, Norway (coordinates: 60°15'06.6"N 5°19'15.4"E). The animals are kept in the animal facility in sea water tanks at 10-12°C and salinity 33 with constant air supply. Once per week, they were fed with mussels and the water was changed. During March-April, when oviposition occurred, cocoons were collected and cultured in the same salinity and temperature conditions than the adults, but they were never fed. Juveniles were fixed at 60 days after oviposition (dao) for whole mount *in situ* hybridization. First, animals were relaxed in 7.4% $MgCl_2$ and then fixed in 4% formaldehyde during 1h at room temperature. The fixative was washed repeatedly with phosphate buffer saline 0.1% Tween-20 (PTw) and specimens were stored at -20°C in 100% methanol.

The gram-negative bacteria *Vibrio diazotrophicus* were purchased from ATCC (catalog number: 33466). The bacteria were resuspended and cultured in Difco™ Marine Broth (Fisher Scientific) at 26°C overnight.

#### Bioinformatic survey of immune genes in *Lineus ruber*

Alignments for conserved domains of the proteins of interest were downloaded from Pfam database (2). When, alignments were not available for our protein of interest in pfam, orthologs of out proteins of interest were collected in NCBI database (www.ncbi.nlm.nih.gov) and alignments were built with MAFFT software version 7 (3). Hmmer profiles were built using HMMER software version 3.2.1 (*www.hmmer.org*) and blasted against the *Lineus ruber* transcriptome and/or the *Notospermus geniculatus* genome. For those proteins for which alignments were not available for their conserved domains, the full sequence of vertebrate and *Drosophila* orthologs were blasted into the *Lineus ruber* transcriptome. The sequences obtained from these surveys were validated by BLAST (4) (*www.blast.ncbi.nlm.nih.gov*). Domain architecture organization was analyzed with SMART (5,6) (*http://smart.embl.de*), hmmer (7) (http://hmmer.org), and NCBI Conserved Domains (8) (https://www.ncbi.nlm.nih.gov/Structure/cdd/wrpsb.cgi) online software.

#### Phylogenetic analyses

Amino acid sequences from *Lineus ruber* were obtained from the bioinformatic survey of immune genes. Sequences from other species were obtained from NCBI database (www.ncbi.nlm.nih.gov). Sequences were aligned using MAFFT software version 7 (3), using the L-INS-I algorithm. The alignment was trimmed with TrimAl software version 1.2 (9). Phylogenetic analyses were performed using the maximum likelihood IQ-TREE software (10) in the CIPRES Science Gateway V.3.3 (11) (*http://www.phylo.org*). For the phylogenetic analysis of DEATH domain containing proteins, LG+F+I+G4 was selected as the best-fit model according to Bayesian Information Criterion (BIC); whereas for the phylogenetic analysis of Nfκb factors VT+I+G4 was selected as the best-fit model. For both the phylogenetic analysis of proteins belonging to the TEP family and the one for Factor B, Factor C and Factor L proteins, LG+R4 was chosen as the best-fit model; whereas the LG+G4 model was chosen for the PGRP phylogenetic analysis. Bootstrap values were calculated running 1000 replicates using ultrafast bootstrap.

**Gene cloning and probe synthesis**

Specific primers for each gene were designed using the MacVector 10.6.0 software based on sequences obtained from transcriptomic surveys. Fragments of each gene were obtained by amplification of cDNA libraries from adult and juvenile stages. The fragments were inserted into pGEM-T Easy vectors (Promega, USA) and transformed into competent *E. coli* cells. Minipreps were prepared using NucleoSpin®Plasmid kit (Macherey-Nagel) and sequenced in the Sequencing facility of the University of Bergen. RNA probes were transcribed using digoxigenin-11-UTP (Roche, USA) with the MEGAscript™ kit (Invitrogen, Thermo Fisher).

**Whole mount *in situ* hybridization (WMISH)**

Whole mount *in sit*u hybridization (WMISH) were performed as described elsewhere (12). Proteinase K digestion was performed during 15 min. Probes were hybridized at a concentration of 1 ng/µl at 67°C during approximately 72h. Anti-digoxigenin-AP antibody (1:5000) was used for probe detection and *in situs* were developed using NBT/BCIP. Samples were washed twice in 100% ethanol and hydrated in descending ethanol steps (75%, 50% and 25%). Next, 3 washes in PBS were performed and they were incubated in 70% glycerol overnight. Samples were mounted in 70% glycerol.

**Imaging**

Histological sections and colorimetric in situ hybridization were imaged using Axiocam HRc camera connected to an Axioscope Ax10 (Zeiss, Oberkochen, Germany).

**Immune-challenge experiments in *Lineus ruber***

Immune-challenge experiments in *Lineus ruber* were designed using a similar approach to other previous studies in which immune-challenge experiments were performed (13–20). Immune-challenge experiments were performed in *Lineus ruber* adult specimens collected specifically for this experiment. The animals were acclimatized for two weeks in the animal facility, in the same conditions as described before, prior to the experiment. Bacterial concentration was assessed by monitoring animals for 48h at different concentrations ($10^6$ bacteria/ml, $10^7$ bacteria/ml, $10^8$ bacteria/ml and $7.6 \times 10^8$ bacteria/ml). The highest concentration was found to be lethal after approximately 3h of exposure, while animals in the remaining concentrations survived for 48h. Thus, we selected a concentration of $10^8$ bacteria/ml for the immune challenge experiments. 64 animals were randomly distributed into 8 groups of 8 animals each. 4 groups were exposed to *Vibrio diazothropic*us ($10^8$ bacteria/ml of sea water), while the other 4 groups were used as controls. Control animals were kept in autoclaved sea water. Prior to infection, both control and immune-challenged animals, were injured with a sterile needle in order to facilitate the penetration of the bacteria in the infected animals. The animals from one control group and one immune-challenged group were frozen in liquid nitrogen and stored individually at -80C.

**RNA extraction, DNA synthesis, qPCR, and data analysis**

mRNA extractions were performed individually for each animal using TRI Reagent™ Solution (ThermoFisher Scientific) and 1-bromo-3-chloropropane (Sigma). cDNA was synthetized using SuperScript™ III First-Strand Synthesis System (Invitrogen), following manufacturer's recommendations. Each reaction contained initially 1 µg of RNA. Specific primers for each gene were designed (MacVector 10.6.0 software) (Supplementary Table 4) and tested prior to the experiments. TLRs gene sequences were obtained from our previous study Orús-Alcalde et al., 2021 (21); whereas the remaining sequences were obtained from the transcriptome survey in this study. qPCRs were performed in Roche LightCycler 480 real-time PCR machine. The master mix contained 1µl of cDNA, 2 µl of primers (10 µM), 7 µl of sterile RNAse free water and 10 µl of mastermix Roche Diagnostics Lightcycler 480 Sybr Green I M (Fisher Scientific). *Actin* was searched in the transcriptome and used as a reference gene (Supplementary Figure 1). For each technical and biological replicate, the gene of interest was normalized with the actin expression levels. Next, each gene of interest was compared between the infected and control animals for each timepoint, to obtain the fold expression using the $2^{-\Delta\Delta CT}$ method (Supplementary Table 4) (22). Data was analyzed with the Light Cycler 480 SW 1.5.1, Microsoft Excel and StatPlus:mac LE v7.

**Illustrations**

All figures plates were assembled with Adobe Illustrator CS6.

**Bibliography**


1. Müller O. Vermivm Terrestrium et Fluviatilium, seu Animalium Infusoriorum, Helminthicorum et Testaceorum, Non Marinorum, Succincta Historia. Copenhagen, Leipzig: Heineck and Faber; 1774. Vol. 1, Part 2.
2. El-Gebali S, Mistry J, Bateman A, Eddy SR, Luciani A, Potter SC, et al. The Pfam protein families database in 2019. Nucleic Acids Research. 2019;47(D1):D427–32.
3. Katoh K, Standley DM. MAFFT Multiple Sequence Alignment Software Version 7: Improvements in Performance and Usability. Molecular Biology and Evolution [Internet]. 2013 Apr 1;30(4):772–80. Available from: https://academic.oup.com/mbe/article-lookup/doi/10.1093/molbev/mst010
4. Altschul S. Gapped BLAST and PSI-BLAST: a new generation of protein database search programs. Nucleic Acids Research [Internet]. 1997;25(17):3389–402. Available from: https://academic.oup.com/nar/article-lookup/doi/10.1093/nar/25.17.3389
5. Schultz J, Milpetz F, Bork P, Ponting CP. SMART, a simple modular architecture research tool: Identification of signaling domains. Proceedings of the National Academy of Sciences [Internet]. 1998 May 26;95(11):5857–64. Available from: http://www.pnas.org/cgi/doi/10.1073/pnas.95.11.5857
6. Letunic I, Doerks T, Bork P. SMART: recent updates, new developments and status in 2015. Nucleic Acids Research [Internet]. 2015 Jan 28;43(D1):D257–60. Available from: http://academic.oup.com/nar/article/43/D1/D257/2439521/SMART-recent-updates-new-developments-and-status

7. Finn RD, Clements J, Arndt W, Miller BL, Wheeler TJ, Schreiber F, et al. HMMER web server: 2015 update. Nucleic Acids Research [Internet]. 2015 Jul 1;43(W1):W30–8. Available from: https://academic.oup.com/nar/article-lookup/doi/10.1093/nar/gkv397
8. Lu S, Wang J, Chitsaz F, Derbyshire MK, Geer RC, Gonzales NR, et al. CDD/SPARCLE: the conserved domain database in 2020. Nucleic Acids Research [Internet]. 2020;48(D1):D265–8. Available from: https://academic.oup.com/nar/article/48/D1/D265/5645006
9. Capella-Gutierrez S, Silla-Martinez JM, Gabaldon T. trimAl: a tool for automated alignment trimming in large-scale phylogenetic analyses. Bioinformatics [Internet]. 2009 Aug 1;25(15):1972–3. Available from: https://academic.oup.com/bioinformatics/article-lookup/doi/10.1093/bioinformatics/btp348
10. Nguyen L-T, Schmidt HA, von Haeseler A, Minh BQ. IQ-TREE: A Fast and Effective Stochastic Algorithm for Estimating Maximum-Likelihood Phylogenies. Molecular Biology and Evolution [Internet]. 2015 Jan;32(1):268–74. Available from: https://academic.oup.com/mbe/article-lookup/doi/10.1093/molbev/msu300
11. Miller MA, Pfeiffer W, Schwartz T. Creating the CIPRES Science Gateway for inference of large phylogenetic trees. In: 2010 Gateway Computing Environments Workshop (GCE) [Internet]. IEEE; 2010. p. 1–8. Available from: http://ieeexplore.ieee.org/document/5676129/
12. Martín-Durán JM, Vellutini BC, Hejnol A. Evolution and development of the adelphophagic, intracapsular Schmidt's larva of the nemertean Lineus ruber. EvoDevo. 2015;6(1):1–18.
13. Deris ZM, Iehata S, Ikhwanuddin M, Sahimi MBMK, Dinh Do T, Sorgeloos P, et al. Immune and bacterial toxin genes expression in different giant tiger prawn, *Penaeus monodon* post-larvae stages following AHPND-causing strain of *Vibrio parahaemolyticus* challenge. Aquaculture Reports [Internet]. 2020 Mar;16:100248. Available from: https://doi.org/10.1016/j.aqrep.2019.100248
14. Lu Y, Li C, Zhang P, Shao Y, Su X, Li Y, et al. Two adaptor molecules of MyD88 and TRAF6 in *Apostichopus japonicus* Toll signaling cascade: Molecular cloning and expression analysis. Developmental & Comparative Immunology [Internet]. 2013 Dec;41(4):498–504. Available from: https://linkinghub.elsevier.com/retrieve/pii/S0145305X13001870
15. Lv Z, Zhang Z, Wei Z, Li C, Shao Y, Zhang W, et al. HMGB3 modulates ROS production via activating TLR cascade in *Apostichopus japonicus*. Developmental & Comparative Immunology [Internet]. 2017 Dec;77:128–37. Available from: https://doi.org/10.1016/j.dci.2017.07.026
16. Peng M, Niu D, Chen Z, Lan T, Dong Z, Tran T-N, et al. Expression of a novel complement C3 gene in the razor clam *Sinonovacula constricta* and its role in innate immune response and hemolysis. Developmental & Comparative Immunology [Internet]. 2017 Aug;73:184–92. Available from: http://dx.doi.org/10.1016/j.dci.2017.03.027
17. Ren Y, Pan H, Pan B, Bu W. Identification and functional characterization of three TLR signaling pathway genes in *Cyclina sinensis*. Fish & Shellfish Immunology [Internet]. 2016 Mar;50:150–9. Available from: http://dx.doi.org/10.1016/j.fsi.2016.01.025
18. Russo R, Chiaramonte M, Matranga V, Arizza V. A member of the *Tlr* family is involved in dsRNA innate immune response in *Paracentrotus lividus* sea urchin. Developmental & Comparative Immunology [Internet]. 2015 Aug;51(2):271–7. Available from: http://dx.doi.org/10.1016/j.dci.2015.04.007
19. Wang M, Yang J, Zhou Z, Qiu L, Wang L, Zhang H, et al. A primitive Toll-like receptor signaling pathway in mollusk Zhikong scallop *Chlamys farreri*. Developmental & Comparative Immunology [Internet]. 2011 Apr;35(4):511–20. Available from: http://dx.doi.org/10.1016/j.dci.2010.12.005
20. Zhu F, Sun B, Wang Z. The crab *Relish* plays an important role in white spot syndrome virus and *Vibrio alginolyticus* infection. Fish and Shellfish Immunology [Internet]. 2019;87(October 2018):297–306. Available from: https://doi.org/10.1016/j.fsi.2019.01.028
21. Orús-Alcalde A, Lu T, Hejnol A. The evolution of the metazoan Toll receptor family and its expression during protostome development. BioRxiv. 2021;
22. Livak KJ, Schmittgen TD. Analysis of relative gene expression data using real-time quantitative PCR and the 2-ΔΔCT method. Methods. 2001;25(4):402–8.
