## Supplementary Table 1 for "The Toll and Imd pathway, the complement system and lectins during immune response of the nemertean *Lineus ruber*"

**Supplementary Table 1. References from Figure 1.**

<table>
<tr><th></th><th>Poriferans</th><th>Cnidarians</th><th>Annelids</th><th>Mollusks</th><th>Nemerteans</th><th>Brachiopods</th><th>Platyhelminthes</th><th>Priapulids</th></tr>
<tr><td>**Spätzle**</td><td>X</td><td>X</td><td>(Davidson *et al.*, 2008)</td><td>(Gerdol and Venier, 2015; Yu *et al.*, 2015)</td><td>X</td><td>(Gerdol *et al.*, 2018)</td><td>X</td><td>(Mapalo *et al.*, 2020)</td></tr>
<tr><td>**TLR**</td><td>(Wiens *et al.*, 2006; Gauthier *et al.*, 2010)</td><td>(Bosch *et al.*, 2009; Gauthier *et al.*, 2010; Poole and Weis, 2014; Brennan *et al.*, 2017; Williams *et al.*, 2018; Leclère *et al.*, 2019)</td><td>(Davidson *et al.*, 2008; Halanych and Kocot, 2014; Orús-Alcalde *et al.*, 2021)</td><td>(Halanych and Kocot, 2014; Gerdol and Venier, 2015; Ren *et al.*, 2016; Orús-Alcalde *et al.*, 2021)</td><td>(Halanych and Kocot, 2014; Luo *et al.*, 2018; Orús-Alcalde *et al.*, 2021)</td><td>(Halanych and Kocot, 2014; Gerdol *et al.*, 2018; Orús-Alcalde *et al.*, 2021)</td><td>(Peiris *et al.*, 2014; Orús-Alcalde *et al.*, 2021)</td><td>(Halanych and Kocot, 2014; Mapalo *et al.*, 2020; Orús-Alcalde *et al.*, 2021)</td></tr>
<tr><td>**MyD88**</td><td>(Wiens *et al.*, 2005; Gauthier *et al.*, 2010)</td><td>(Sullivan *et al.*, 2007; Gauthier *et al.*, 2010; Brennan *et al.*, 2017)</td><td>(Davidson *et al.*, 2008)</td><td>(Toubiana *et al.*, 2014; Gerdol and Venier, 2015; Ren *et al.*, 2017)</td><td>X</td><td rowspan="3">(Gerdol *et al.*, 2018)</td><td rowspan="2">(Peiris *et al.*, 2014)</td><td>(Mapalo *et al.*, 2020)</td></tr>
<tr><td>**Irak**</td><td>(Wiens *et al.*, 2006; Gauthier *et al.*, 2010)</td><td>X</td><td>(Davidson *et al.*, 2008)</td><td>(Toubiana *et al.*, 2014; Gerdol and Venier, 2015)</td><td>X</td><td rowspan="2">(Mapalo *et al.*, 2020)</td></tr>
<tr><td>**NFκB-p65**</td><td>(Gauthier *et al.*, 2010)</td><td>(Sullivan *et al.*, 2007; Gauthier *et al.*, 2010; Brennan *et al.*, 2017)</td><td>(Davidson *et al.*, 2008)</td><td>(Gerdol and Venier, 2015)</td><td>X</td><td>(Forsthoefel *et al.*, 2012)</td></tr>
<tr><td>**Imd**</td><td>X</td><td>X</td><td>X</td><td>(Toubiana *et al.*, 2014)</td><td>X</td><td>(Gerdol *et al.*, 2018)</td><td>X</td><td rowspan="5">(Mapalo *et al.*, 2020)</td></tr>
<tr><td>**Fadd**</td><td>X</td><td>X</td><td>X</td><td>(Zhang *et al.*, 2011; Hu *et al.*, 2019)</td><td>X</td><td rowspan="4">(Gerdol *et al.*, 2018)</td><td>X</td></tr>
<tr><td>**Dredd**</td><td>X</td><td>X</td><td>X</td><td>(Romero *et al.*, 2011)</td><td>X</td><td>X</td></tr>
<tr><td>**Relish**</td><td>X</td><td>X</td><td>(Davidson *et al.*, 2008)</td><td>(Toubiana *et al.*, 2014)</td><td>X</td><td>X</td></tr>
<tr><td>**tPGRP**</td><td>X</td><td>X</td><td>X</td><td>(Gerdol and Venier, 2015)</td><td>X</td><td>X</td></tr>
<tr><td>**C3**</td><td>(Srivastava *et al.*, 2010)</td><td>(Dishaw *et al.*, 2005; Miller *et al.*, 2007; Kimura *et al.*,</td><td>X</td><td>(Castillo *et al.*, 2009; Prado-Alvarez *et al.*, 2009; Gerdol</td><td>X</td><td>(Gerdol *et al.*, 2018; Gorbushin, 2018)</td><td>(Peiris *et al.*, 2014)</td><td>X</td></tr>
</table>

| | | | | | | | | |
|---|---|---|---|---|---|---|---|---|
| | | 2009; Fujito *et al.*, 2010) | | and Venier, 2015; Wang *et al.*, 2017, 2019; Gorbushin, 2018) | | | | |
| **Factor B** | | (Kimura *et al.*, 2009; Poole *et al.*, 2016) | X | (Prado-Alvarez *et al.*, 2009; Wang *et al.*, 2017; Gorbushin, 2018) | X | | X | X |
| **CR** | | (Gorbushin, 2018) | (Altincicek and Vilcinskas, 2007) | (Wang *et al.*, 2017; Gorbushin, 2018) | X | (Gorbushin, 2018) | X | X |
| **C6** | | (Kimura *et al.*, 2009) | X | (Gerdol and Venier, 2015) | X | (Gerdol *et al.*, 2018) | X | X |
| **C1r/s** | | (Nonaka and Kimura, 2006) | | | X | (Nonaka and Kimura, 2006) | X | X |
| **MASP** | | (Kimura *et al.*, 2009; Poole *et al.*, 2016; Gorbushin, 2019) | (Nonaka and Kimura, 2006) | (Nonaka and Kimura, 2006; Wang *et al.*, 2017) | X | | X | X |
| **MreM** | | X | (Gorbushin, 2019) | (Gorbushin, 2019) | X | (Gorbushin, 2019) | X | X |
| **C1q** | X | X | (Gorbushin, 2019) | (Gerdol and Venier, 2015; Wang *et al.*, 2017) | X | (Gorbushin, 2019) | X | X |
| **FreDC2** | X | (Gorbushin, 2019) | | (Gerdol and Venier, 2015; Skazina and Gorbushin, 2016; Gorbushin, 2019) | X | (Gorbushin, 2019) | X | X |
| **CTLDC2** | X | (Gorbushin, 2019) | | (Gerdol and Venier, 2015; Gorbushin, 2019) | X | (Gerdol and Venier, 2015; Gorbushin, 2019) | X | X |

<table>
<tr><th></th><th>Nematodes</th><th>Arthropods</th><th>Tardigrades</th><th>Echinoderms</th><th>Amphioxus</th><th>Tunicates</th><th>Vertebrates</th></tr>
<tr><td>Spatzle</td><td>(Mapalo et al., 2020)</td><td rowspan="5">(Anderson and Nüsslein-Volhard, 1984; Anderson et al., 1985; Lemaitre et al., 1996; Hoffmann and Reichhart, 2002; Valanne et al., 2011; Palmer and Jiggins, 2015; Mapalo et al., 2020)</td><td>(Mapalo et al., 2020)</td><td>X</td><td>X</td><td>X</td><td>(Valanne et al., 2011)</td></tr>
<tr><td>TLR</td><td>(Mapalo et al., 2020; Orús-Alcalde et al., 2021)</td><td>(Mapalo et al., 2020; Orús-Alcalde et al., 2021)</td><td rowspan="3">(Hibino et al., 2006; Tassia et al., 2017)</td><td>(Tassia et al., 2017; Ji et al., 2018)</td><td>(Sasaki et al., 2009; Denoeud et al., 2010; Tassia et al., 2017)</td><td>(Medzhitov et al., 1997; Rock et al., 1998)</td></tr>
<tr><td>MyD88</td><td>(Mapalo et al., 2020)</td><td rowspan="3">(Mapalo et al., 2020)</td><td>(Yuan et al., 2009; Tassia et al., 2017)</td><td rowspan="2">(Azumi et al., 2003; Tassia et al., 2017)</td><td>(Wesche et al., 1997; Medzhitov et al., 1998)</td></tr>
<tr><td>Irak</td><td>(Mapalo et al., 2020)</td><td>(Tassia et al., 2017)</td><td rowspan="2">(Wesche et al., 1997)</td></tr>
<tr><td>NFκB-p65</td><td>(Mapalo et al., 2020)</td><td>(Hibino et al., 2006)</td><td>(Yuan et al., 2009; Tassia et al., 2017)</td><td>(Azumi et al., 2003)</td></tr>
<tr><td>Imd</td><td rowspan="4">(Mapalo et al., 2020)</td><td>(Kaneko et al., 2006)</td><td rowspan="5">(Mapalo et al., 2020)</td><td>X</td><td>X</td><td>X</td><td rowspan="4">(Myllymäki et al., 2014)</td></tr>
<tr><td>Fadd</td><td>(Hu and Yang, 2000; Naitza et al., 2002)</td><td>X</td><td>X</td><td>X</td></tr>
<tr><td>Dredd</td><td>(Hu and Yang, 2000)</td><td>X</td><td>X</td><td>X</td></tr>
<tr><td>Relish</td><td>(Dushay et al., 1996; Shin et al., 2002)</td><td>X</td><td>X</td><td>X</td></tr>
<tr><td>tPGRP</td><td>(Dziarski and Gupta, 2006; Mapalo et al., 2020)</td><td>(Werner et al., 2000, 2003; Choe et al., 2002; Gottar et al., 2002; Rämet et al., 2002; Takehana et al., 2002, 2004; Kaneko et al., 2006; Paik et al., 2017; Chevée et al., 2019)</td><td>X</td><td>X</td><td>X</td><td>(Kang et al., 1998)</td></tr>
<tr><td>C3</td><td rowspan="3">(The C.elegans Sequencing Consortium, 1998)</td><td>(Adams, 2000; Zhu et al., 2005; Ariki et al., 2008; Sekiguchi et al., 2012; Palmer and Jiggins, 2015; Gorbushin, 2018)</td><td>X</td><td>(Al-Sharif et al., 1998; Nair et al., 2005; Smith et al., 2006) (Hibino et al., 2006)</td><td>(Suzuki et al., 2002)</td><td>(Nonaka et al., 1999; Marino et al., 2002; Raftos et al., 2002; Azumi et al., 2003; Nair et al., 2005)</td><td>(Janssen et al., 2005)</td></tr>
<tr><td>Factor B</td><td>(Adams, 2000; Zhu et al., 2005; Palmer and Jiggins, 2015; Sekiguchi and Nonaka, 2015; Gorbushin, 2018)</td><td>X</td><td>(Nair et al., 2005; Smith et al., 2006)</td><td>(He et al., 2008)</td><td>(Smith et al., 1998; Nair et al., 2005)</td><td>(Milder et al., 2007)</td></tr>
<tr><td>CR</td><td>(Adams, 2000; Gorbushin, 2018)</td><td>X</td><td>(Gorbushin, 2018)</td><td>X</td><td>(Gorbushin, 2018)</td><td>(Ahearn and Fearon, 1989;</td></tr>
</table>

<table>
<tr><td></td><td rowspan="5"></td><td></td><td></td><td></td><td></td><td></td><td>Helmy et al., 2006; Vorup-Jensen and Jensen, 2018)</td></tr>
<tr><td>C6</td><td>X</td><td>X</td><td>X</td><td>(Suzuki et al., 2002)</td><td>(Azumi et al., 2003)</td><td>(Bhakdi and Tranum-Jensen, 1978; Rosado et al., 2008)</td></tr>
<tr><td>C1r/s</td><td>(Nonaka and Kimura, 2006)</td><td>X</td><td>(Nonaka and Kimura, 2006)</td><td rowspan="2">(Nonaka and Kimura, 2006)</td><td>(Nonaka and Kimura, 2006)</td><td>(Arlaud et al., 2002; Girija et al., 2013)</td></tr>
<tr><td>MASP</td><td>X</td><td>X</td><td rowspan="2">(Gorbushin, 2019)</td><td>(Ji et al., 1997; Azumi et al., 2003; Nonaka and Kimura, 2006; Gorbushin, 2019)</td><td>(Matsushita and Fujita, 1992)</td></tr>
<tr><td>MreM</td><td>(Gorbushin, 2019)</td><td>X</td><td>X</td><td>X</td><td>X</td></tr>
<tr><td>C1q</td><td></td><td>X</td><td>X</td><td>(Hibino et al., 2006)</td><td>X</td><td rowspan="2">(Azumi et al., 2003)</td><td>(Svehag et al., 1972; Carland and Gerwick, 2010)</td></tr>
<tr><td>FreDC2</td><td></td><td>(Han et al., 2018)</td><td>X</td><td>X</td><td>(Huang et al., 2011)</td><td>(Ichijo et al., 1993)</td></tr>
<tr><td>CTLDC2</td><td></td><td>X</td><td>X</td><td>X</td><td>X</td><td>(Sekine et al., 2001; Azumi et al., 2003)</td><td>(Sastry et al., 1989)</td></tr>
</table>

**Bibliography Supplementary Table 1.**


Adams, M. D. (2000) ‘The genome sequence of *Drosophila melanogaster*’, *Science*, 287(5461), pp. 2185–2195. doi: 10.1126/science.287.5461.2185.

Ahearn, J. M. and Fearon, D. T. (1989) ‘Structure and Function of the Complement Receptors, CR1 (CD35) and CR2 (CD21)’, *Advances in Immunology*, 46(C), pp. 183–219. doi: 10.1016/S0065-2776(08)60654-9.

Al-Sharif, W. Z. *et al.* (1998) ‘Sea urchin coelomocytes specifically express a homologue of the complement component C3.’, *Journal of immunology (Baltimore, Md. : 1950)*, 160(6), pp. 2983–97. Available at: http://www.ncbi.nlm.nih.gov/pubmed/9510203.

Altincicek, B. and Vilcinskas, A. (2007) ‘Analysis of the immune-related transcriptome of a lophotrochozoan model, the marine annelid *Platynereis dumerilii*’, *Frontiers in Zoology*, 4(1), p. 18. doi: 10.1186/1742-9994-4-18.

Anderson, K. V., Jürgens, G. and Nüsslein-Volhard, C. (1985) ‘Establishment of dorsal-ventral polarity in the *Drosophila* embryo: Genetic studies on the role of the *Toll* gene product’, *Cell*, 42(3), pp. 779–789. doi: 10.1016/0092-8674(85)90274-0.

Anderson, K. V. and Nüsslein-Volhard, C. (1984) ‘Information for the dorsal–ventral pattern of the *Drosophila* embryo is stored as maternal mRNA’, *Nature*, 311(5983), pp. 223–227. doi: 10.1038/311223a0.

Ariki, S. *et al.* (2008) ‘Factor C Acts as a lipopolysaccharide-responsive C3 convertase in horseshoe crab complement activation’, *The Journal of Immunology*, 181(11), pp. 7994–8001. doi: 10.4049/jimmunol.181.11.7994.

Arlaud, G. J. *et al.* (2002) ‘Structural biology of the C1 complex of complement unveils the mechanisms of its activation and proteolytic activity’, *Molecular Immunology*, 39(7–8), pp. 383–394. doi: 10.1016/S0161-5890(02)00143-8.

Azumi, K. *et al.* (2003) ‘Genomic analysis of immunity in a Urochordate and the emergence of the vertebrate immune system: “waiting for Godot”’, *Immunogenetics*, 55(8), pp. 570–581. doi: 10.1007/s00251-003-0606-5.

Bhakdi, S. and Tranum-Jensen, J. (1978) ‘Molecular nature of the complement lesion.’, *Proceedings of the National Academy of Sciences*, 75(11), pp. 5655–5659. doi: 10.1073/pnas.75.11.5655.

Bosch, T. C. G. *et al.* (2009) ‘Uncovering the evolutionary history of innate immunity: The simple metazoan *Hydra* uses epithelial cells for host defence’, *Developmental & Comparative Immunology*, 33(4), pp. 559–569. doi: 10.1016/j.dci.2008.10.004.

Brennan, J. J. *et al.* (2017) ‘Sea anemone model has a single Toll-like receptor that can function in pathogen detection, NF-κB signal transduction, and development’, *Proceedings of the National Academy of Sciences*, 114(47), pp. E10122–E10131. doi: 10.1073/pnas.1711530114.

Carland, T. M. and Gerwick, L. (2010) ‘The C1q domain containing proteins: Where do they come from and what do they do?’, *Developmental and Comparative Immunology*. Elsevier Ltd, 34(8), pp. 785–790. doi: 10.1016/j.dci.2010.02.014.

Castillo, M. G., Goodson, M. S. and McFall-Ngai, M. (2009) ‘Identification and molecular characterization of a complement C3 molecule in a lophotrochozoan, the Hawaiian bobtail squid *Euprymna scolopes*’, *Developmental & Comparative Immunology*, 33(1), pp. 69–76. doi: 10.1016/j.dci.2008.07.013.

Chevée, V. *et al.* (2019) ‘The peptidoglycan recognition protein PGRP-LE regulates the *Drosophila* immune response against the pathogen *Photorhabdus*’, *Microbial Pathogenesis*. Elsevier Ltd, 136(August), p. 103664. doi: 10.1016/j.micpath.2019.103664.

Choe, K. *et al.* (2002) ‘Requirement for a Peptidoglycan Recognition Protein (PGRP) in Relish Activation and Antibacterial Immune Responses in Drosophila’, *Science*, 296(5566), pp. 359–362. doi: 10.1126/science.1070216.

Davidson, C. R. *et al.* (2008) ‘*Toll-like receptor* genes (TLRs) from *Capitella capitata* and *Helobdella robusta* (Annelida)’, *Developmental & Comparative Immunology*, 32(6), pp. 608–612. doi: 10.1016/j.dci.2007.11.004.

Denoeud, F. *et al.* (2010) ‘Plasticity of animal genome architecture unmasked by rapid evolution of a pelagic tunicate’, *Science*, 330(6009), pp. 1381–1385. doi: 10.1126/science.1194167.

Dishaw, L. J., Smith, S. L. and Bigger, C. H. (2005) ‘Characterization of a C3-like cDNA in a coral: phylogenetic implications’, *Immunogenetics*, 57(7), pp. 535–548. doi: 10.1007/s00251-005-0005-1.

Dushay, M. S., Asling, B. and Hultmark, D. (1996) ‘Origins of immunity: Relish, a compound <i>Rel<i/>-like gene in the antibacterial defense of *Drosophila*.’, *Proceedings of the National Academy of Sciences*, 93(19), pp. 10343–10347. doi: 10.1073/pnas.93.19.10343.

Dziarski, R. and Gupta, D. (2006) ‘The peptidoglycan recognition proteins (PGRPs)’, *Genome Biology*, 7(8), pp. 1–13. doi: 10.1186/gb-2006-7-8-232.

Forsthoefel, D. J. *et al.* (2012) ‘An RNAi Screen Reveals Intestinal Regulators of Branching Morphogenesis, Differentiation, and Stem Cell Proliferation in Planarians’, *Developmental Cell*, 23(4), pp. 691–704. doi: 10.1016/j.devcel.2012.09.008.

Fujito, N. T., Sugimoto, S. and Nonaka, M. (2010) ‘Evolution of thioester-containing proteins revealed by cloning and characterization of their genes from a cnidarian sea anemone, *Haliplanella lineate*’, *Developmental & Comparative Immunology*. Elsevier Ltd, 34(7), pp. 775–784. doi: 10.1016/j.dci.2010.02.011.

Gauthier, M. E. A., Du Pasquier, L. and Degnan, B. M. (2010) ‘The genome of the sponge *Amphimedon queenslandica* provides new perspectives into the origin of Toll-like and interleukin 1 receptor pathways’, *Evolution & Development*, 12(5), pp. 519–533. doi: 10.1111/j.1525-142X.2010.00436.x.

Gerdol, M. *et al.* (2018) ‘Genetic and molecular basis of the immune system in the brachiopod *Lingula anatina*’,

*Developmental & Comparative Immunology*, 82, pp. 7–30. doi: 10.1016/j.dci.2017.12.021.
Gerdol, M. and Venier, P. (2015) ‘An updated molecular basis for mussel immunity’, *Fish & Shellfish Immunology*. Elsevier Ltd, 46(1), pp. 17–38. doi: 10.1016/j.fsi.2015.02.013.
Girija, V. *et al.* (2013) ‘Structural basis of the C1q/C1s interaction and its central role in assembly of the C1 complex of complement activation’, *Proceedings of the National Academy of Sciences*, 110(34), pp. 13916–13920. doi: 10.1073/pnas.1615704113.
Gorbushin, A. M. (2018) ‘Immune repertoire in the transcriptome of *Littorina littorea* reveals new trends in lophotrochozoan proto-complement evolution’, *Developmental & Comparative Immunology*. Elsevier Ltd, 84, pp. 250–263. doi: 10.1016/j.dci.2018.02.018.
Gorbushin, A. M. (2019) ‘Derivatives of the lectin complement pathway in Lophotrochozoa’, *Developmental & Comparative Immunology*. Elsevier, 94(November 2018), pp. 35–58. doi: 10.1016/j.dci.2019.01.010.
Gottar, M. *et al.* (2002) ‘The *Drosophila* immune response against Gram-negative bacteria is mediated by a peptidoglycan recognition protein’, *Nature*, 416(6881), pp. 640–644. doi: 10.1038/nature734.
Halanych, K. M. and Kocot, K. M. (2014) ‘Repurposed transcriptomic data facilitate discovery of innate immunity *Toll-Like Receptor (TLR)* genes across Lophotrochozoa’, *The Biological Bulletin*, 227(2), pp. 201–209. doi: 10.1086/BBLv227n2p201.
Han, K. *et al.* (2018) ‘Novel fibrinogen-related protein with single FReD contributes to the innate immunity of <i>Macrobrachium rosenbergii</i >’, *Fish & Shellfish Immunology*. Elsevier, 82(June), pp. 350–360. doi: 10.1016/j.fsi.2018.08.036.
He, Y. *et al.* (2008) ‘Molecular and immunochemical demonstration of a novel member of Bf/C2 homolog in amphioxus *Branchiostoma belcheri*: Implications for involvement of hepatic cecum in acute phase response’, *Fish & Shellfish Immunology*, 24(6), pp. 768–778. doi: 10.1016/j.fsi.2008.03.004.
Helmy, K. Y. *et al.* (2006) ‘CRIg: A macrophage complement receptor required for phagocytosis of circulating pathogens’, *Cell*, 124(5), pp. 915–927. doi: 10.1016/j.cell.2005.12.039.
Hibino, T. *et al.* (2006) ‘The immune gene repertoire encoded in the purple sea urchin genome’, *Developmental Biology*, 300(1), pp. 349–365. doi: 10.1016/j.ydbio.2006.08.065.
Hoffmann, J. A. and Reichhart, J.-M. (2002) ‘*Drosophila* innate immunity: an evolutionary perspective’, *Nature Immunology*, 3(2), pp. 121–126. doi: 10.1038/ni0202-121.
Hu, G. *et al.* (2019) ‘Molecular cloning and characterization of FADD from the manila clam *Ruditapes philippinarum*’, *Fish & Shellfish Immunology*. Elsevier, 88(March), pp. 556–566. doi: 10.1016/j.fsi.2019.03.033.
Hu, S. and Yang, X. (2000) ‘dFADD, a novel death domain-containing adapter protein for the *Drosophila* caspase DREDD’, *Journal of Biological Chemistry*, 275(40), pp. 30761–30764. doi: 10.1074/jbc.C000341200.
Huang, H. *et al.* (2011) ‘Functional characterization of a Ficolin-mediated complement pathway in amphioxus’, *Journal of Biological Chemistry*, 286(42), pp. 36739–36748. doi: 10.1074/jbc.M111.245944.
Ichijo, H. *et al.* (1993) ‘Molecular cloning and characterization of ficolin, a multimeric protein with fibrinogen- and collagen-like domains’, *Journal of Biological Chemistry*. Â© 1993 ASBMB. Currently published by Elsevier Inc; originally published by American Society for Biochemistry and Molecular Biology., 268(19), pp. 14505–14513. doi: 10.1016/S0021-9258(19)85267-5.
Janssen, B. J. C. *et al.* (2005) ‘Structures of complement component C3 provide insights into the function and evolution of immunity’, *Nature*, 437(7058), pp. 505–511. doi: 10.1038/nature04005.
Ji, J. *et al.* (2018) ‘Characterization of the TLR family in *Branchiostoma lanceolatum* and discovery of a novel TLR22-like involved in dsRNA recognition in amphioxus’, *Frontiers in Immunology*, 9, pp. 1–15. doi: 10.3389/fimmu.2018.02525.
Ji, X. *et al.* (1997) ‘Ancient origin of the complement lectin pathway revealed by molecular cloning of mannan binding protein-associated serine protease from a urochordate, the Japanese ascidian, *Halocynthia roretzi*’, *Proceedings of the National Academy of Sciences*, 94(12), pp. 6340–6345. doi: 10.1073/pnas.94.12.6340.
Kaneko, T. *et al.* (2006) ‘PGRP-LC and PGRP-LE have essential yet distinct functions in the *Drosophila* immune response to monomeric DAP-type peptidoglycan’, *Nature Immunology*, 7(7), pp. 715–723. doi: 10.1038/ni1356.
Kang, D. *et al.* (1998) ‘A peptidoglycan recognition protein in innate immunity conserved from insects to humans’, *Proceedings of the National Academy of Sciences*, 95(17), pp. 10078–10082. doi: 10.1073/pnas.95.17.10078.
Kimura, A., Sakaguchi, E. and Nonaka, M. (2009) ‘Multi-component complement system of Cnidaria: C3, Bf, and MASP genes expressed in the endodermal tissues of a sea anemone, *Nematostella vectensis*’, *Immunobiology*, 214(3), pp. 165–178. doi: 10.1016/j.imbio.2009.01.003.
Leclère, L. *et al.* (2019) ‘The genome of the jellyfish *Clytia hemisphaerica* and the evolution of the cnidarian life-cycle’, *Nature Ecology & Evolution*. Springer US, 3(5), pp. 801–810. doi: 10.1038/s41559-019-0833-2.
Lemaitre, B. *et al.* (1996) ‘The dorsoventral regulatory gene cassette *Spätzle/Toll/Cactus* controls the potent antifungal response in *Drosophila* adults’, *Cell*, 86(6), pp. 973–983. doi: 10.1016/S0092-8674(00)80172-5.
Luo, Y.-J. *et al.* (2018) ‘Nemertean and phoronid genomes reveal lophotrochozoan evolution and the origin of bilaterian heads’, *Nature Ecology & Evolution*. Springer US, 2(1), pp. 141–151. doi: 10.1038/s41559-017-0389-y.
Mapalo, M. A. *et al.* (2020) ‘The unique antimicrobial recognition and signaling pathways in tardigrades with a comparison across Ecdysozoa’, *G3 Genes|Genomes|Genetics*, 10(3), pp. 1137–1148. doi: 10.1534/g3.119.400734.
Marino, R. *et al.* (2002) ‘Complement in urochordates: cloning and characterization of two *C3-like* genes in the

ascidian *Ciona intestinalis*', *Immunogenetics*, 53(12), pp. 1055–1064. doi: 10.1007/s00251-001-0421-9.
Matsushita, M. and Fujita, T. (1992) 'Activation of the classical complement pathway by mannose-binding protein in association with a novel C1s-like serine protease.', *Journal of Experimental Medicine*, 176(6), pp. 1497–1502. doi: 10.1084/jem.176.6.1497.
Medzhitov, R. *et al.* (1998) 'MyD88 Is an adaptor protein in the hToll/IL-1 receptor family signaling pathways', *Molecular Cell*, 2(2), pp. 253–258. doi: 10.1016/S1097-2765(00)80136-7.
Medzhitov, R., Preston-Hurlburt, P. and Janeway, C. A. (1997) 'A human homologue of the *Drosophila* Toll protein signals activation of adaptive immunity', *Nature*, 388(6640), pp. 394–397. doi: 10.1038/41131.
Milder, F. J. *et al.* (2007) 'Factor B structure provides insights into activation of the central protease of the complement system', *Nature Structural & Molecular Biology*, 14(3), pp. 224–228. doi: 10.1038/nsmb1210.
Miller, D. J. *et al.* (2007) 'The innate immune repertoire in Cnidaria - ancestral complexity and stochastic gene loss', *Genome Biology*, 8(4), p. R59. doi: 10.1186/gb-2007-8-4-r59.
Myllymäki, H., Valanne, S. and Rämet, M. (2014) 'The *Drosophila* Imd signaling pathway', *The Journal of Immunology*, 192(8), pp. 3455–3462. doi: 10.4049/jimmunol.1303309.
Nair, S. V., Ramsden, A. and Raftos, D. A. (2005) 'Ancient origins: Complement in invertebrates', *Invertebrate Survival Journal*, 2(2), pp. 114–123.
Naitza, S. *et al.* (2002) 'The *Drosophila* immune defense against gram-negative infection requires the Death protein dFADD', *Immunity*, 17(5), pp. 575–581. doi: 10.1016/S1074-7613(02)00454-5.
Nonaka, M. *et al.* (1999) 'Opsonic complement C3 in the solitary ascidian, Halocynthia roretzi', *Molecular Immunology*, 35(6–7), p. 363. doi: 10.1016/S0161-5890(98)90662-9.
Nonaka, M. and Kimura, A. (2006) 'Genomic view of the evolution of the complement system', *Immunogenetics*, 58(9), pp. 701–713. doi: 10.1007/s00251-006-0142-1.
Orús-Alcalde, A., Lu, T. and Hejnol, A. (2021) 'The evolution of the metazoan Toll receptor family and its expression during protostome development', *BioRxiv*. doi: 10.1101/2021.02.01.429095.
Paik, D. *et al.* (2017) 'SLC46 Family Transporters Facilitate Cytosolic Innate Immune Recognition of Monomeric Peptidoglycans', *The Journal of Immunology*, 199(1), pp. 263–270. doi: 10.4049/jimmunol.1600409.
Palmer, W. J. and Jiggins, F. M. (2015) 'Comparative genomics reveals the origins and diversity of arthropod immune systems', *Molecular Biology and Evolution*, 32(8), pp. 2111–2129. doi: 10.1093/molbev/msv093.
Peiris, T. H., Hoyer, K. K. and Oviedo, N. J. (2014) 'Innate immune system and tissue regeneration in planarians: An area ripe for exploration', *Seminars in Immunology*. Elsevier Ltd, 26(4), pp. 295–302. doi: 10.1016/j.smim.2014.06.005.
Poole, A. Z., Kitchen, S. A. and Weis, V. M. (2016) 'The role of complement in cnidarian-dinoflagellate symbiosis and immune challenge in the sea anemone *Aiptasia pallida*', *Frontiers in Microbiology*, 7, pp. 1–18. doi: 10.3389/fmicb.2016.00519.
Poole, A. Z. and Weis, V. M. (2014) 'TIR-domain-containing protein repertoire of nine anthozoan species reveals coral–specific expansions and uncharacterized proteins', *Developmental & Comparative Immunology*. Elsevier Ltd, 46(2), pp. 480–488. doi: 10.1016/j.dci.2014.06.002.
Prado-Alvarez, M. *et al.* (2009) 'Characterization of a C3 and a factor B-like in the carpet-shell clam, *Ruditapes decussatus*', *Fish & Shellfish Immunology*. Elsevier Ltd, 26(2), pp. 305–315. doi: 10.1016/j.fsi.2008.11.015.
Raftos, D. A. *et al.* (2002) 'A complement component C3-like protein from the tunicate, *Styela plicata*', *Developmental & Comparative Immunology*, 26(4), pp. 307–312. doi: 10.1016/S0145-305X(01)00080-5.
Rämet, M. *et al.* (2002) 'Functional genomic analysis of phagocytosis and identification of a *Drosophila* receptor for *E. coli*', *Nature*, 416(6881), pp. 644–648. doi: 10.1038/nature735.
Ren, Y. *et al.* (2016) 'Identification and functional characterization of three TLR signaling pathway genes in *Cyclina sinensis*', *Fish & Shellfish Immunology*. Elsevier Ltd, 50, pp. 150–159. doi: 10.1016/j.fsi.2016.01.025.
Ren, Y. *et al.* (2017) 'Comparative and evolutionary analysis of an adapter molecule MyD88 in invertebrate metazoans', *Developmental & Comparative Immunology*. Elsevier Ltd, 76, pp. 18–24. doi: 10.1016/j.dci.2017.05.007.
Rock, F. L. *et al.* (1998) 'A family of human receptors structurally related to *Drosophila* Toll', *Proceedings of the National Academy of Sciences*, 95(2), pp. 588–593. doi: 10.1073/pnas.95.2.588.
Romero, A. *et al.* (2011) 'New Insights into the Apoptotic Process in Mollusks: Characterization of Caspase Genes in *Mytilus galloprovincialis*', *PLoS ONE*. Edited by A. Bergmann, 6(2), p. e17003. doi: 10.1371/journal.pone.0017003.
Rosado, C. J. *et al.* (2008) 'The MACPF/CDC family of pore-forming toxins', *Cellular Microbiology*, 10(9), pp. 1765–1774. doi: 10.1111/j.1462-5822.2008.01191.x.
Sasaki, N. *et al.* (2009) 'Toll-like Receptors of the ascidian *Ciona intestinalis*', *Journal of Biological Chemistry*, 284(40), pp. 27336–27343. doi: 10.1074/jbc.M109.032433.
Sastry, K. *et al.* (1989) 'The human mannose-binding protein gene. Exon structure reveals its evolutionary relationship to a human pulmonary surfactant gene and localization to chromosome 10.', *Journal of Experimental Medicine*, 170(4), pp. 1175–1189. doi: 10.1084/jem.170.4.1175.
Sekiguchi, R., Fujito, N. T. and Nonaka, M. (2012) 'Evolution of the thioester-containing proteins (TEPs) of the arthropoda, revealed by molecular cloning of TEP genes from a spider, *Hasarius adansoni*', *Developmental & Comparative Immunology*. Elsevier Ltd, 36(2), pp. 483–489. doi: 10.1016/j.dci.2011.05.003.
Sekiguchi, R. and Nonaka, M. (2015) 'Evolution of the complement system in protostomes revealed by de novo

transcriptome analysis of six species of Arthropoda', *Developmental & Comparative Immunology*. Elsevier Ltd, 50(1), pp. 58–67. doi: 10.1016/j.dci.2014.12.008.
Sekine, H. *et al.* (2001) 'An ancient lectin-dependent complement system in an ascidian: Novel lectin isolated from the plasma of the solitary ascidian, *Halocynthia roretzi*', *The Journal of Immunology*, 167(8), pp. 4504–4510. doi: 10.4049/jimmunol.167.8.4504.
Shin, S. W. *et al.* (2002) 'Characterization of three alternatively spliced isoforms of the Rel/NF- B transcription factor Relish from the mosquito *Aedes aegypti*', *Proceedings of the National Academy of Sciences*, 99(15), pp. 9978–9983. doi: 10.1073/pnas.162345999.
Skazina, M. A. and Gorbushin, A. M. (2016) 'Characterization of the gene encoding a fibrinogen-related protein expressed in *Crassostrea gigas* hemocytes', *Fish & Shellfish Immunology*. Elsevier Ltd, 54, pp. 586–588. doi: 10.1016/j.fsi.2016.05.017.
Smith, L. C. *et al.* (2006) 'The sea urchin immune system', *Invertebrate Survival Journal*, 3(1), pp. 25–39.
Smith, L. C., Shih, C. S. and Dachenhausen, S. G. (1998) 'Coelomocytes express SpBf, a homologue of factor B, the second component in the sea urchin complement system.', *Journal of immunology (Baltimore, Md. : 1950)*, 161(12), pp. 6784–93. Available at: http://www.ncbi.nlm.nih.gov/pubmed/9862709.
Srivastava, M. *et al.* (2010) 'The Amphimedon queenslandica genome and the evolution of animal complexity', *Nature*. Nature Publishing Group, 466(7307), pp. 720–726. doi: 10.1038/nature09201.
Sullivan, J. C. *et al.* (2007) 'Rel homology domain-containing transcription factors in the cnidarian *Nematostella vectensis*', *Development Genes and Evolution*, 217(1), pp. 63–72. doi: 10.1007/s00427-006-0111-6.
Suzuki, M. M., Satoh, N. and Nonaka, M. (2002) 'C6-like and C3-like molecules from the cephalochordate, amphioxus, suggest a cytolytic complement system in invertebrates', *Journal of Molecular Evolution*, 54(5), pp. 671–679. doi: 10.1007/s00239-001-0068-z.
Svehag, S., Manhem, L. and Bloth, B. (1972) 'Ultrastructure of Human C1q Protein', *Nature New Biology*, 238(82), pp. 117–118. doi: 10.1038/newbio238117a0.
Takehana, A. *et al.* (2002) 'Overexpression of a pattern-recognition receptor, peptidoglycan-recognition protein-LE, activates imd/relish-mediated antibacterial defense and the prophenoloxidase cascade in *Drosophila* larvae', *Proceedings of the National Academy of Sciences*, 99(21), pp. 13705–13710. doi: 10.1073/pnas.212301199.
Takehana, A. *et al.* (2004) 'Peptidoglycan recognition protein (PGRP)-LE and PGRP-LC act synergistically in *Drosophila* immunity', *The EMBO Journal*, 23(23), pp. 4690–4700. doi: 10.1038/sj.emboj.7600466.
Tassia, M. G., Whelan, N. V. and Halanych, K. M. (2017) 'Toll-like receptor pathway evolution in deuterostomes', *Proceedings of the National Academy of Sciences*, 114(27), pp. 7055–7060. doi: 10.1073/pnas.1617722114.
The C.elegans Sequencing Consortium (1998) 'Genome sequence of the nematode *C. elegans*: A platform for investigating biology', *Science*, 282(5396), pp. 2012–2018. doi: 10.1126/science.282.5396.2012.
Toubiana, M. *et al.* (2014) 'Toll signal transduction pathway in bivalves: Complete cds of intermediate elements and related gene transcription levels in hemocytes of immune stimulated *Mytilus galloprovincialis*', *Developmental & Comparative Immunology*, 45(2), pp. 300–312. doi: 10.1016/j.dci.2014.03.021.
Valanne, S., Wang, J.-H. and Rämet, M. (2011) 'The *Drosophila* Toll signaling pathway', *The Journal of Immunology*, 186(2), pp. 649–656. doi: 10.4049/jimmunol.1002302.
Vorup-Jensen, T. and Jensen, R. K. (2018) 'Structural immunology of complement receptors 3 and 4', *Frontiers in Immunology*, 9(NOV), pp. 1–20. doi: 10.3389/fimmu.2018.02716.
Wang, Lingling *et al.* (2017) 'The RNA-seq analysis suggests a potential multi-component complement system in oyster *Crassostrea gigas*', *Developmental & Comparative Immunology*. Elsevier Ltd, 76, pp. 209–219. doi: 10.1016/j.dci.2017.06.009.
Wang, N. *et al.* (2019) 'Molecular cloning of complement component *C3* gene from pearl mussel, *Hyriopsis cumingii* and analysis of the gene expression in response to tissue transplantation', *Fish & Shellfish Immunology*. Elsevier, 94(301), pp. 288–293. doi: 10.1016/j.fsi.2019.09.010.
Werner, T. *et al.* (2000) 'A family of peptidoglycan recognition proteins in the fruit fly *Drosophila melanogaster*', *Proceedings of the National Academy of Sciences*, 97(25), pp. 13772–13777. doi: 10.1073/pnas.97.25.13772.
Werner, T. *et al.* (2003) 'Functional diversity of the *Drosophila PGRP-LC* gene cluster in the response to lipopolysaccharide and peptidoglycan', *Journal of Biological Chemistry*, 278(29), pp. 26319–26322. doi: 10.1074/jbc.C300184200.
Wesche, H. *et al.* (1997) 'MyD88: An adapter that recruits IRAK to the IL-1 receptor complex', *Immunity*, 7(6), pp. 837–847. doi: 10.1016/S1074-7613(00)80402-1.
Wiens, M. *et al.* (2005) 'Innate Immune Defense of the Sponge *Suberites domuncula* against Bacteria Involves a MyD88-dependent Signaling Pathway', *Journal of Biological Chemistry*, 280(30), pp. 27949–27959. doi: 10.1074/jbc.M504049200.
Wiens, M. *et al.* (2006) 'Toll-like receptors are part of the innate immune defense system of sponges (Demospongiae: Porifera)', *Molecular Biology and Evolution*, 24(3), pp. 792–804. doi: 10.1093/molbev/msl208.
Williams, L. M. *et al.* (2018) 'A conserved Toll-like receptor-to-NF-κB signaling pathway in the endangered coral *Orbicella faveolata*', *Developmental & Comparative Immunology*, 79, pp. 128–136. doi: 10.1016/j.dci.2017.10.016.
Yu, M. *et al.* (2015) 'The first mollusk spätzle homolog gene in the clam, Paphia undulate', *Fish & Shellfish Immunology*. Elsevier, 47(2), pp. 712–716. doi: 10.1016/j.fsi.2015.10.017.

Yuan, S. *et al.* (2009) ‘An amphioxus TLR with dynamic embryonic expression pattern responses to pathogens and activates NF-κB pathway via MyD88’, *Molecular Immunology*, 46(11–12), pp. 2348–2356. doi: 10.1016/j.molimm.2009.03.022.
Zhang, L., Li, L. and Zhang, G. (2011) ‘Gene discovery, comparative analysis and expression profile reveal the complexity of the *Crassostrea gigas* apoptosis system’, *Developmental & Comparative Immunology*, 35(5), pp. 603–610. doi: 10.1016/j.dci.2011.01.005.
Zhu, Y. *et al.* (2005) ‘The ancient origin of the complement system’, *The EMBO Journal*, 24(2), pp. 382–394. doi: 10.1038/sj.emboj.7600533.
