## Supplementary Table 2 for "The Toll and Imd pathway, the complement system and lectins during immune response of the nemertean *Lineus ruber*"

**Supplementary Table 2. BLAST hits from the components of the *Lineus ruber* Toll and Imd pathways and C3 and Factor B proteins from the complement system.** The ordinal numbers before each hit indicate the position of each hit. All 1rst positions are indicated, but hits for uncharacterized proteins and repeated proteins are omitted.

| Protein | Best hits in BLAST | E-value | Identity % |
|---|---|---|---|
| **MyD88** | *1st: Myeloid differentiation primary response protein 88b [Cyprinus carpio]* | $5e^{-53}$ | 38.93% |
| **Irak** | *1st: Receptor-like kinase LIP2 isoform X1 [Mizuhopecten yessoensis]*<br>*3rd: Interleukin-1 receptor-associated kinase 4-like [Lingula anatina]* | $1e^{-66}$<br>$1e^{-54}$ | 31.58%<br>37.50% |
| **Dorsal/NFkB-p65** | *1st: Nuclear factor-kB [Cyclina sinensis]*<br>*2nd: Rel/NF-kB [Haliotis discus discus]* | $1e^{-144}$<br>$1e^{-137}$ | 58.58%<br>55.01% |
| **PGRP-1** | *1st: Peptidoglycan recognition protein S2 [Hyriopsis cumingii]* | $1e^{-70}$ | 58.43% |
| **PGRP-2** | *1st: Peptidoglycan recognition protein 1 [Gopherus evgoodei]* | $4e^{-30}$ | 59.74% |
| **Fadd** | *1st: FAS-associated death domain protein [Vombatus ursinus]* | $5e^{-22}$ | 30.14% |
| **Imd** | *1st: Imd [Scylla paramamosain]* | $2e^{-6}$ | 31.33% |
| **Dredd** | *1st: Caspase-8 [Lingula anatina]* | $2e^{-81}$ | 52.87% |
| **Relish/Nfkb-p105/100** | *1st: Hypothetical protein CAPTEDRAFT_181359 [Capitella teleta]*<br>*2nd: Nuclear factor NF-kappa-B p105 subunit [Mus musculus]* | $6e^{-165}$<br>$1e^{-159}$ | 35.18%<br>37.20% |
| **C3-1** | *1st: Complement C3 [Branchiostoma belcheri]* | 0 | 33.43% |
| **C3-2** | *1st Venom factor [Lingula anatina]*<br>*Many complement C3 hits* | 0 | 35.36% |
| **Factor B-1** | *1st: complement component 2/factor B variant 2 [Tachypleus tridentatus]* | $1e^{-117}$ | 30.63% |
| **Factor B-2** | *1st: complement factor B-like isoform X1 [Centruroides sculpturatus]* | $1e^{-111}$ | 29.39% |
| **Factor B-3** | *1st: complement factor B-like isoform X2 [Centruroides sculpturatus]* | $5e^{-73}$ | 26.93% |
| **Factor B-4** | *1st: complement C2-like [Branchiostoma floridae]* | $1e^{-67}$ | 27.99% |
