## Supplementary Table 3 for "The Toll and Imd pathway, the complement system and lectins during immune response of the nemertean *Lineus ruber*"

**Supplementary Table 3 – qPCR primers**

| Gene | Forward | Reverse |
|---|---|---|
| ***TLRα3*** | TCCTTTCCAATGTCACACACC | CAGTTCACTCCCACGAAGTTG |
| ***TLRα4*** | CACGGGGAAGACCAATCAATG | GGCGTTCATCCATACTCCAGTG |
| ***TLRβ1*** | CAACCAAACACGACTGTCAATGC | ACCACGAAACCCGCCTTTACTG |
| ***TLRβ2*** | GGTCGGTGCTAATGGACGATTC | CGCAATGGGTGTCAAACAGAC |
| ***imd*** | TGCTGGAAGTTGATTCAGTCGTC | GAGTAAGTTCACCAATGTCGCTACC |
| ***C3-1*** | AACTGAGGTTTGCGGACCACTG | CCCATCCTTTCCCATCACAAG |
| ***fred-c5*** | AGCGACAATGACCTCTGGTTTG | ATCCTGTTTCCACGAGTGCCAC |
| ***c-lectin2*** | GCAGGATGAAGCGAATGAAGAGAC | TTGGTTTCCCTTGCTCCACC |
