## Supplementary Table 4 for "The Toll and Imd pathway, the complement system and lectins during immune response of the nemertean *Lineus ruber*"

**Supplementary Table 4 – qPCR data.** The fold expression (infected vs control animals) was calculated for two or three biological replicates using the $2^{-\Delta\Delta CT}$ method (Livak and Schmittgen, 2001) and the average fold for each gene and timepoint was calculated. ANOVA tests were performed to test if the expression changes were significant (being p-value<0.05 significant, and p-value<0.01 very significant).

| | | Biol rep1 fold | Biol rep2 fold | Biol rep3 fold | average fold | ANOVA (p-value) |
|---|---|---|---|---|---|---|
| *TLRβ1* | 3h | 1,424618706 | 0,872725273 | 1,025273273 | 1,107539084 | 0,72353 |
| | 6h | 1,888244099 | 1,902122988 | - | 1,895183544 | 0,02267 |
| | 12h | 2,269780114 | 1,687290681 | - | 1,978535397 | 0,09720 |
| | 24h | 1,977314653 | 2,427317932 | - | 2,202316293 | 0,03501 |
| *TLRβ2* | 3h | 1,310207569 | 1,084141059 | 1,82914563 | 1,407831419 | 0,25090 |
| | 6h | 1,511413211 | 1,625168555 | - | 1,568290883 | 0,03098 |
| | 12h | 1,711460603 | 2,073969913 | - | 1,892715258 | 0,04234 |
| | 24h | 0,397526706 | 0,322693018 | 0,296743892 | 0,338987872 | 0,03408 |
| *TLRα4* | 3h | 1,720083203 | 0,721480929 | 1,151228819 | 1,19759765 | 0,69964 |
| | 6h | 0,694941696 | 0,549138062 | - | 0,622039879 | 0,18294 |
| | 12h | 0,445139487 | 0,685608491 | 0,486936057 | 0,539228011 | 0,01710 |
| | 24h | 0,940432402 | 0,982833302 | 0,931507036 | 0,951590913 | 0,15524 |
| *TLRα3* | 3h | 0,770712666 | 0,88612137 | 0,509016228 | 0,721950088 | 0,19115 |
| | 6h | 0,991771835 | 0,655044145 | 0,569710196 | 0,738842059 | 0,23204 |
| | 12h | 5,09707983 | 4,721579974 | - | 4,909329902 | 0,00061 |
| | 24h | 2,080604352 | 3,037394612 | - | 2,558999482 | 0,02242 |
| *imd* | 1h | 1,268955589 | 1,379070073 | - | 1,324012831 | 0,06202 |
| | 3h | 1,220337336 | 1,431148335 | - | 1,325742836 | 0,10270 |
| | 6h | 0,84933177 | 0,840361226 | - | 0,844846498 | 0,07246 |
| | 12h | 0,980954942 | 0,958883163 | - | 0,969919053 | 0,82726 |
| | 24h | 0,979199504 | 1,027469348 | - | 1,003334426 | 0,95341 |
| *fred-c5* | 3h | 0,507040267 | 0,549820852 | 0,422012607 | 0,492957908 | 0,00820 |
| | 6h | 0,702191241 | 0,363793956 | 0,54129213 | 0,535759109 | 0,06982 |
| | 12h | 1,519471342 | 1,613763518 | - | 1,56661743 | 0,01575 |
| | 24h | 0,712430328 | 0,742843412 | - | 0,72763687 | 0,23309 |
| *C3-1* | 3h | 0,570733533 | 0,411926117 | 0,907594136 | 0,630084595 | 0,19045 |
| | 6h | 0,72536245 | 0,823001108 | - | 0,774181779 | 0,30180 |
| | 12h | 0,470388394 | 0,352462282 | - | 0,411425338 | 0,02743 |
| | 24h | 1,376087528 | 1,9831417 | 1,748147014 | 1,702458747 | 0,02736 |
| *c-lectin2* | 3h | 0,919192617 | 0,672750036 | - | 0,795971327 | 0,37113 |
| | 6h | 1,110689559 | 1,041811505 | - | 1,076250532 | 0,75218 |
| | 12h | 1,634192621 | 1,465888482 | - | 1,550040551 | 0,00686 |
| | 24h | 1,738981084 | 1,039868357 | - | 1,389424721 | 0,38358 |
